# Multiplexed CRISPRi Reveals a Transcriptional Switch Between KLF Activators and Repressors in the Maturing Neocortex

**DOI:** 10.1101/2025.02.07.636951

**Authors:** Ryan W Kirk, Liwei Sun, Ruixuan Xiao, Erin A Clark, Sacha Nelson

## Abstract

A critical phase of mammalian brain development takes place after birth. Neurons of the mouse neocortex undergo dramatic changes in their morphology, physiology, and synaptic connections during the first postnatal month, while properties of immature neurons, such as the capacity for robust axon outgrowth, are lost. The genetic and epigenetic programs controlling prenatal development are well studied, but our understanding of the transcriptional mechanisms that regulate postnatal neuronal maturation is comparatively lacking. By integrating chromatin accessibility and gene expression data from two subtypes of neocortical pyramidal neurons in the neonatal and maturing brain, we predicted a role for the Krüppel-Like Factor (KLF) family of Transcription Factors in the developmental regulation of neonatally expressed genes. Using a multiplexed CRISPR Interference (CRISPRi) knockdown strategy, we found that a shift in expression from KLF activators (Klf6, Klf7) to repressors (Klf9, Klf13) during early postnatal development functions as a transcriptional ‘switch’ to first activate, then repress a set of shared targets with cytoskeletal functions including *Tubb2b* and *Dpysl3*. We demonstrate that this switch is buffered by redundancy between KLF paralogs, which our multiplexed CRISPRi strategy is equipped to overcome and study. Our results indicate that competition between activators and repressors within the KLF family regulates a conserved component of the postnatal maturation program that may underlie the loss of intrinsic axon growth in maturing neurons. This could facilitate the transition from axon growth to synaptic refinement required to stabilize mature circuits.

## Introduction

From birth onwards, neurons in the cortex continue to mature and refine their prenatally acquired identity (Molyneaux et al., 2015; Prince et al., 2024; Wallace & Pollen, 2024; Yuan et al., 2022). This is especially pronounced in the first postnatal month of rodent brain development, which is characterized by sequential overlapping periods of axon growth, synaptogenesis, and synapse refinement (Faust et al., 2021; Price et al., 2006; Südhof, 2018; Turrigiano & Nelson, 2004). Many human neuropsychiatric and neurodevelopmental disorders result from disruptions in one or more of these processes, but how these processes are regulated at a molecular level remains incompletely understood (Gonçalves et al., 2013; Krishnan et al., 2015; Marín, 2016; Sekar et al., 2016). While the precise timing and nuances of the postnatal maturation program are tailored to the circuit-specific needs of cell types in different cortical layers, many physiological outcomes are remarkably conserved across cortical cell types and species (Gonzalez-Burgos et al., 2008; Kroon et al., 2019; Marchetto et al., 2019; Shi et al., 2012). These include regulated increases in the size of the soma (Benedetti et al., 2020), the number and strength of inhibitory and excitatory presynaptic connections (Desai et al., 2002; Etherington & Williams, 2011), expansion of the dendritic arbor (Koenderink & Uylings, 1995; Wu et al., 1999), and a decline in intrinsic excitability through rebalancing of membrane ion channels (McCormick & Prince, 1987; Okaty et al., 2009; Zhang 2004). A number of stereotyped postnatal switches in the expression of individual ion channel and neurotransmitter receptor subunits have been characterized, such as an increase in the Ca^2+^-permeable AMPA receptor subunit GLUA2 (Sheng et al., 1994; Williams et al., 1993), an exchange of the prenatal GABRA2 receptor for GABRA1 subtype (Fritschy et al., 1994), and a transition from GLUN2B-containing NMDA receptors to the GLUN2A-containing variety found in the adult (Brill & Huguenard, 2008; Kumar et al., 2002). Such switches appear to be conserved properties of postnatal development, as they have been identified in multiple cortical regions, cell types, and species albeit with modestly differing degrees and kinetics (Ciceri & Studer, 2024; Law et al., 2003). It has been proposed that such switches reflect functional adaptations to changing requirements for sensitivity, plasticity, and stability at maturing cortical synapses, but in many cases the precise function of the switch is still unclear (Cho et al., 2009; Philpot et al., 2007).

In the neocortex of the mouse and other altricial mammals, prenatally initiated programs of stereotyped axon guidance and growth continue throughout the first two weeks of postnatal life, resulting in the formation of the corpus callosum and major subcortical tracts (De León Reyes et al., 2020; Gianino et al., 1999). Following this period of axon outgrowth, cortical neurons uniformly relinquish their capacity for robust axon growth to give way to a period of axon retraction and synaptic pruning (Riccomagno & Kolodkin, 2015). This transition is terminal, as mature neurons remain incapable of re-growing their axons following injury or insult (Bradke, 2022; Bregman et al., 1989; Ramon y Cajal, 1928). The loss of regenerative capacity has been attributed to a variety of cell-extrinsic and -intrinsic factors inherent to the central nervous system, as peripheral neurons do not exhibit such a dramatic loss of growth potential (Huebner & Strittmatter, 2009). Extrinsically, developmental increases in extracellular matrix proteins and components of the myelin sheath prevent axon growth through both physical and molecular mechanisms (Laabs et al., 2005; Yiu & He, 2006). However, mature neurons still struggle to regrow damaged axons even when these impediments are removed and young neurons can still extend axons through the inhibitory postnatal extracellular environment, exposing the influence of a powerful intrinsic brake on axon growth (Blackmore & Letourneau, 2006; Li et al., 1995; Steinmetz et al., 2005). The biological nature of such a brake is still under investigation, but many therapeutic strategies for restoring axon growth rely on restoring specific neonatal attributes to adult neurons (Filbin, 2006; Hilton & Bradke, 2017; Mahar & Cavalli, 2018). Thus, neuronal maturation is characterized by both the acquisition of specialized functions and loss of shared properties of immature neurons that were critical for circuit development. Viewed differently, terminal differentiation of cortical neuron subtypes can also be conceived of as a generalized process characterized by the repression of neonatal identity.

Studies of genetically or anatomically defined cell types over the first 1-2 months of postnatal development have characterized the genetic and electrophysiological maturation of cortical neurons and interneurons in exquisite detail (Ciceri et al., 2024; Kroon et al., 2019; Okaty et al., 2009; Patel et al., 2022; Yuan et al., 2022). While these genetic and physiological parameters change in parallel, surprisingly little is known about the Transcription Factors (TFs) responsible. Unraveling transcriptional circuits in higher eukaryotes is often confounded by the presence of gene paralogs able to compensate or compete with one another by binding the same DNA sequence (Badis et al., 2009; Jolma et al., 2013; Majidi et al., 2019; Weirauch et al., 2014). One such family is the diverse Krüppel-like Factor (KLF) family of TFs, which contains 17 members that share a preference for a GC-rich DNA motif frequently located in promoter regions (Kaczynski et al., 2003; Moore et al., 2011). Members of this family are expressed in unique combinations in most major cell types of the body, and are known to differentially compensate and compete with one another to affect gene expression in surprising, often non-linear, ways in non-neuronal tissues (Eaton et al., 2008; Jiang et al., 2008). Therefore, ascribing a unitary role to any individual member of this diverse family is complicated by intrafamilial interactions. Despite this complexity, several KLF paralogs have been shown to play roles in promoting or repressing CNS axon growth (Moore et al., 2009). Specifically, overexpression of the pro-growth TFs Klf6 or Klf7 has been shown to promote regeneration of Corticospinal Tract axons *in vivo* after spinal cord injury (Blackmore et al., 2012; Wang et al., 2018), while removal of growth-inhibiting Klf9 has been shown to enhance Retinal Ganglion Cell axon growth after optic nerve crush (Apara et al., 2017). The convergence of opposing members of the KLF family on a singular biological function has generated speculation that they might regulate the same set of genes in opposite directions, but experimental evidence for this hypothesis is scarce (Galvao et al., 2018). Such a detailed dissection of KLF family function in neurons would require methods robust to potential compensation by paralogues.

Multiplexed gene loss-of-function approaches offer an experimental workaround to the problem of TF redundancy: by eliminating a TF and its known or hypothesized compensator, previously obscured phenotypes can be unmasked (Ahmed et al., 2024; Kim et al., 2021; Majidi et al., 2019). RNA-interference has been used as an alternative to intersectional transgenic strategies to demonstrate genetic redundancy and study genetic epistasis *in vitro*, yet it is prone to off-target effects and shows variable potency of knockdown *in vivo* (Chen et al., 2021; Horn et al., 2011; Sawyer et al., 2011). CRISPR-based loss-of-function is an emerging alternative to RNAi, but CRISPR-mediated double-strand break repair in post-mitotic cells leads to heterogenous outcomes complicating interpretation (González et al., 2014; Mandegar et al., 2016). CRISPR Interference (CRISPRi) has recently gained traction as a versatile tool for repressing gene expression. This approach targets a nuclease-dead Cas9 fused to a KRAB effector domain (dCas9-KRAB) to genomic loci of interest using sequence-specific single guide RNAs (sgRNAs) (Gilbert et al., 2013; Qi et al., 2013). Multiplexed gene knockdown using CRIPSRi has been achieved by expressing multiple sgRNAs targeting different genes (Gilbert et al., 2014; McCarty et al., 2020; Zheng et al., 2018). CRISPRi has higher efficacy and specificity than RNAi, but *in vivo* applications have been limited by the large size of the dCas9-KRAB effector (Lau et al., 2019; Li et al., 2020; Zheng et al., 2018). The recent creation of a mouse containing a floxed dCas9-KRAB allele enables multiplexed and cell type-specific loss-of-function without the AAV payload restrictions surrounding delivery of the fusion enzyme (Gemberling et al., 2021). However, this system has not yet been utilized to study the regulatory logic of neuronal TFs *in vivo*.

In this study, we used bioinformatics to identify putative TFs regulating postnatal development in excitatory neurons of the mouse cortex. By focusing on the program shared by upper and deep layer neurons to silence genes related to pre- and neonatal functions, we identified potential roles for developmentally regulated KLF paralogs in regulating this process. Using a novel *in vivo* multiplexed CRISPRi approach, we demonstrate that the postnatally upregulated TFs Klf9 and Klf13 are redundant repressors of a subset of developmentally downregulated genes. Conversely, we demonstrated that the neonatally-expressed transcriptional activators Klf6 and Klf7 are required for early postnatal expression of developmentally regulated genes with cytoskeletal functions. The high degree of overlap observed among bidirectionally regulated targets of the KLF repressor and activator pairs suggests that the KLF family functions as a ‘transcriptional switch’ to turn off neonatally expressed genes with known or suggested roles in axon growth, possibly through competition among family members for the same GC-rich motifs in shared targets’ promoters. Furthermore, the timing of this switch coincides with the transition from a period of robust axon growth to one of synaptogenesis in the mouse CNS (Kiyoshi & Tedeschi, 2020; Lewis et al., 2013), suggesting a possible transcriptional mechanism contributing to this developmental process. Moreover, this study provides proof-of-concept for the utility of multiplexed CRISPR interference for studying redundancy and competition within large TF families in the mammalian brain.

## Results

### Pyramidal cells in Layers 4 and 6 share a common Gene Regulatory Program in early postnatal development

To identify putative transcriptional regulators of early postnatal development, we used RNA sequencing (RNA-seq) and Assay for Transposase-Accessible Chromatin with Sequencing (ATAC-seq) to profile changes in gene expression and chromatin accessibility between P2 and P30 in two distinct excitatory cortical cell types: RORβ^+^ Layer 4 neurons and Bmp3^+^ Layer 6 neurons (**Figure 1A**). Layer 4 neurons were purified by Fluorescence Activated Cell Sorting (FACS) from Rorb^GFP/+^ knock-in mice, which express GFP primarily in excitatory thalamocortical input neurons of layer 4 (<u>Liu et al., 2013</u>; Clark et al., 2020). Layer 6 neurons expressing mCitrine were purified by manual sorting or FACS from the 56L mouse line, which labels a unique population of excitatory corticothalamic neurons in lower layer 6 that project to non-primary thalamic nuclei and subcortical targets (Shima et al., 2016). Layer 4 ATAC-seq data from P30 and RNA-seq data from P2 were previously published in Clark et al., 2020, and Layer 6 ATAC-seq and RNA-seq data from P30 were published in Sugino et al., 2019.

**Figure 1.**
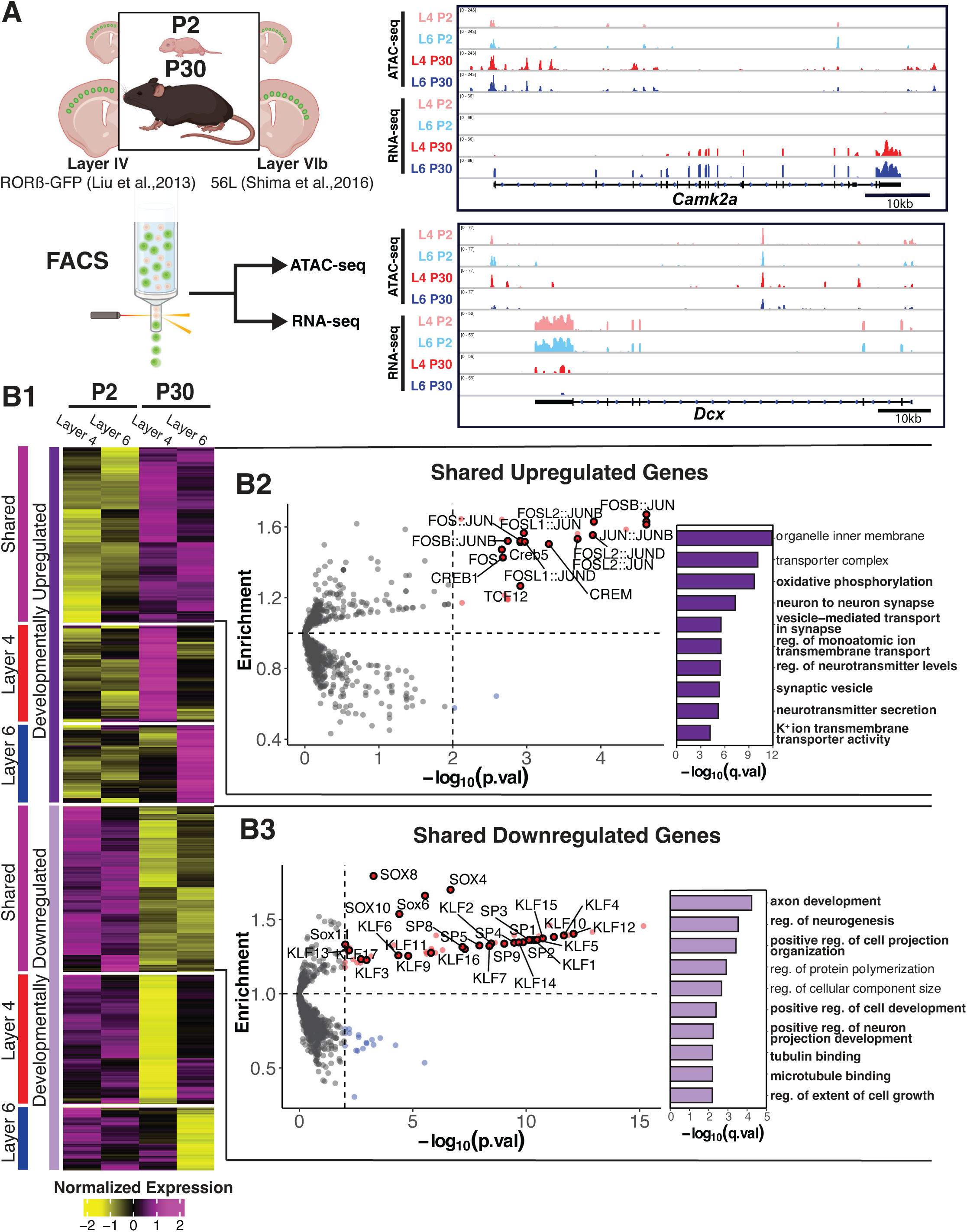
Shared gene expression changes in maturing layer 4 and 6 cortical pyramidal neurons. **A)** Experimental Workflow: Rorb-GFP+/- (L4) or 56L (L6) mice were sacrificed at P2 or P30 and GFP+ neurons were isolated by FACS for RNA-seq or ATAC-seq (left). IGV tracks of Gene Expression (RNA-seq) and Chromatin Accessibility (ATAC-seq) of neonatally expressed Doublecortin (*Dcx*, top right) and adult-expressed *CaMKIIa* (bottom right). **B)** RNA-Seq of FACS-sorted Rorb^+^ Layer 4 neurons and Bmp3^+^ Layer 6 neurons at P2 and P30 (n=4 mice per group). **B1:** Heatmap of All DEGs identified in either layer between P2 and P30 identifies k=6 unique clusters (DEGs defined as |log2FoldChange|>1; p.adj<0.01 by Wald’s Test with BH correction; min TPM>10). Data plotted as group averages of z-scored relative log gene expression. **B2 (right):** Top Gene Ontology terms enriched in shared upregulated genes relative to all genes eligible for Differential Expression (Fisher Exact tests with BH adjustment). **B2 (left):** Transcription Factor Motif Enrichment in ATAC-seq peaks overlapping the Promoter (TSS -1000/+200 bp) of shared upregulated genes relative to all Promoter peaks using the JASPAR 2022 CORE Motif set (Fisher Exact tests with BH adjustment). **B3:** Same as **B2** but for shared downregulated genes.

Although cell type-specific gene regulatory programs are critical for establishing characteristic physiological and morphological properties of neurons in different layers (Lodato & Arlotta, 2015), we focused on the more general program of gene expression and chromatin accessibility changes conserved across the two distinct cell types. RNA-seq and ATAC-seq data demonstrated a substantial effect of postnatal age with a lesser contribution by genetic lineage that was more pronounced in mature neurons consistent with Yuan et al., 2022 (**Figure S1.1A**). RNA-seq identified 5,579 Differentially Expressed Protein-Coding Genes (DEGs; Criteria: absolute Fold-Change≥2, adjusted P value<0.01, TPM≥10) changing between P2 and P30 in at least one cell type. K-means clustering separated these DEGs into 6 groups comprised of a larger group of shared and smaller groups of developmentally up- or down-regulated cell type-specific genes (**Figure 1B1**). Of 5,579 DEGs, 2,684 (48.1%) were similarly regulated in layers 4 and 6 (1,380 up- and 1,304 down-regulated). Gene Ontology (GO) overrepresentation analysis for shared upregulated genes revealed that these genes are involved in processes such as vesicle-mediated transport at the synapse (q-value = 3.32×10^-6^), regulation of neurotransmitter levels (4.18×10^-6^), and regulation of ion transport (3.32×10^-6^), consistent with the synaptic and electrophysiological changes occurring during the first month of life (**Figure 1B2, right**). In contrast, shared downregulated genes were enriched for GO terms axon development (q-value = 6.25×10^-5^), regulation of neurogenesis (3.16×10^-4^), and regulation of projection organization (4.09×10^-4^), consistent with the loss of mitotic and axon growth capacity in the maturing cortex (**Figure 1B3, right**).

We next examined the ATAC-seq data to identify putative cis-regulatory elements and associated Transcription Factor Binding Sites (TFBSs) facilitating the observed changes in gene expression. Of the 242,127 peaks of accessible chromatin identified across conditions, we found 38,321 Differentially Accessible Regions (DARs; Criteria: absolute Fold-Change≥2, adjusted P value<0.01, Reads≥10) that gain or lose accessibility between P2 and P30 in at least one cell type (**Figure S1.1B**). Interestingly, of the 11,434 DARs (29.9%) with significant changes in both cell types, 11,419 (99.6%) changed in the same direction, indicating the presence of a shared regulatory program amidst cell type-specific differences. We found 2,378 promoter DARs which formed 2 clusters representing sites that either gain (1,220 Gained Shared DARs) or lose (1,158 Lost Shared DARs) accessibility in *both* cell types (**Figure S1.2A**). The lack of cell type-specific chromatin changes at promoters is consistent with the general understanding that promoter accessibility does not distinguish between closely related cell types as well as accessibility at distal genomic elements (Gray et al., 2017; Mo et al., 2015; Yoshida et al., 2019). Notably, this property made promoters an attractive genomic feature to focus on when looking for a shared developmental regulatory program. Therefore, we looked for enrichment of TFBSs in gained or lost promoter DARs to identify putative TFs regulating chromatin accessibility changes. This revealed that motifs recognized by the AP-1 family, bHLH, and MEF family of TFs were significantly overrepresented in shared gained promoter DARs (**Figure S1.2B1**). In contrast, we found a strong enrichment for the motif recognized by Homeodomain TFs and the POU family in shared lost promoter DARs (**Figure S1.2B2**), both of which contain members that play significant roles in specifying neuronal fate during prenatal development (Briscoe et al., 2000). An identical TFBS enrichment analysis of Differentially Accessible Regions (DARs) assigned to intronic and intergenic elements was performed, yielding similar results apart from strong MEF enrichment (data not shown).

To relate chromatin accessibility changes to observed changes in gene expression during early postnatal development, we examined the relationship between these features. Surprisingly, fewer than 10% of shared DEGs contained a DAR in their promoter and we observed a weakly discernable change in the aggregate accessibility of promoter elements associated with shared developmental DEGs (**Figure S1.2C**). Promoter elements were also proportionally underrepresented among DARs, suggesting that transcriptional regulation from these sites could instead arise from coordinated transitions in the TFs bound to constitutively accessible promoter sites. To identify putative TFs regulating shared DEGs through chromatin accessibility-independent mechanisms, we performed TFBS enrichment analysis on *all* promoter ATAC-seq peaks associated with developmentally regulated genes. In promoters of shared developmentally upregulated genes, we found significant enrichment for motifs recognized by the AP-1 TFs FOS, C-JUN, and CREB (1.47- to 1.67-fold; P adj. between 2.15×10^-3^ and 2.40×10^-5^) and for the motif recognized by the TCF family (1.26-fold; P adj. = 1.21×10^-3^; **Figure 1B2, left**). The presence of the AP-1 motif in the promoters of genes upregulated during the early postnatal period could reflect the arrival of structured input driving activity-dependent mechanisms associated with the transcriptional maturation of neurons (Stroud et al., 2020; West & Greenberg, 2011).

The most significantly overrepresented motifs associated with downregulated genes were those recognized by members of the Krüppel-Like Factor (KLF)/Sp TF family in the promoters of downregulated genes (P adj. between 6.25×10^-3^ and 7.88×10^-13^; enrichment 1.29-1.40-fold; **Figure 1B3, left**). The KLF/Sp family contains 18 KLFs and 4 Sp factors that share a highly conserved DNA binding domain and preference for the same GC-rich motif, so the observed enrichment likely represents the activity of a subset of KLF/Sp family members (Suske et al., 2005). After simplifying the motif set by using only the root motifs from JASPAR’s motif clustering, the root motif describing the cluster that includes all KLF and Sp TFs remained the most significantly and highly enriched motif (1.35-fold, P adj. = 1.26×10^-6^; **Figure S1.1C**). While members of the KLF/Sp family have been implicated in a variety of neuronal functions, a role for this family in pyramidal cell maturation has not been described with the exception of Klf9, which has been shown to influence dentate granule cell maturation (Scobie et al., 2009). Of note, multiple members of the KLF family have established roles in regulating axon growth, which was one of one of the most enriched GO terms among shared downregulated DEGs (Moore et al., 2009). Altogether, this made the KLFs an attractive TF family for further investigation. Significant enrichment in these promoters was also observed for motifs recognized by members of the SOX family of TFs (1.33- to 1.79-fold; P adj. between 9.24×10^-3^ and 2.09×10^-7^). Sox1-3 are required for the maintenance of Neural Stem Cell pluripotency while Sox4 and Sox11 have established roles in establishing early neuronal identity, so the presence of their recognized motif in downregulated genes likely reflects their regulation of prenatally expressed genes (Bergsland et al., 2006; Bylund et al., 2003).

An identical motif enrichment analysis focused on intronic or intragenic peaks with proximity to DEGs was uninformative, likely due to the difficulty of accurately assigning peaks distal to an annotated TSS to their target gene(s) (Fulco et al., 2019) (data not shown). Together, these results highlight the existence of a developmental program shared by upper and deep layer neurons amidst their cell type-specific gene regulatory networks. The shared program may involve the upregulation of genes required for the acquisition of mature electrophysiological properties concomitant with downregulation of genes required for early developmental processes of cell migration and axon growth (**Figure 1B1**).

### The expression of KLF activators and repressors are oppositely regulated during early postnatal development

The striking enrichment of the KLF/Sp motif in shared downregulated DEG promoters along with the KLF family’s functional agreement with enriched GO terms and their understudied role in the developing cortex made them appealing candidates for loss-of-function study. While Motif Enrichment Analyses can predict TFBSs that are abundant in regions of interest undergoing some notable biological change, they rarely offer insight into the specific TFs underpinning that change since individual motifs are typically recognized by several TFs. This is particularly true for the KLF family, which consists of 18 members with conserved DNA binding domains resulting in a preference for the same motif (Suske et al., 2005). To parse this family, we used 2 common-sense criteria for identifying candidate TFs that impact developmental cortical gene expression: sufficient expression of the TFs in pyramidal cells and developmental regulation of the TFs themselves. We first looked at the expression levels of all KLF paralogs in 6 major excitatory cell types of the adult mouse cortex using additional RNA-seq data published in Sugino et al., 2019. This narrowed our pool of candidate TFs to 6 moderately- to highly-expressed KLFs: *Klf6*, *Klf7*, *Klf9*, *Klf10*, *Klf12*, and *Klf13* (**Figure S2A**). Appropriate to our interest in studying transcriptional networks shared across pyramidal cells, the expression of all KLF paralogs did not differ substantially across excitatory neuron subtypes in different layers in our data or in the Allen Institute’s single-cell RNA-seq data from mouse brain (**Figure S2B**, Yao et al., 2021).

Next, we investigated how the expression of expressed KLFs change during postnatal development. Previous studies have found evidence for downregulation of Klf6 and Klf7 and upregulation of KLF9 during a similar postnatal period in the brain (Laub et al., 2001; Moore et al., 2009; Scobie et al., 2009; Wang et al., 2018). Consistent with these observations, we detected a significant increase in expression of *Klf9* and *Klf13* in both Layer 4 (*Klf9*: 5.67-fold, P adj. = 1.7 ×10^-100^; *Klf13*: 3.96-fold, 8.5×10^-53^) and Layer 6 (*Klf9*: 4.11-fold, 3.32×10^-66^; *Klf13*: 2.55-fold, 3.46×10^-26^) populations and a modest but significant decrease in *Klf7* in both cell types (Layer 6: 3.13-fold, 1.29×10^-40^; Layer 6: 1.49-fold, 3.50×10^-6^) and *Klf6* in Layer 4 (3.31-fold, 4.87×10^-87^) in our RNA-seq data (**Figure S2C**). To examine the expression of the highest expressed activating (*Klf7*) and repressive (*Klf9*) KLFs at finer temporal resolution in other genetically defined excitatory cortical cell types, we leveraged recent RNA-seq data from sorted Layer II/III and Layer 6 neurons across postnatal development published in Yuan et al., 2022. This clearly demonstrated a sharp and synchronous decline in *Klf7* and steep increase in *Klf9* expression between P7 and P21 among cells in Layers II/III and upper Layer 6, suggesting that the switch between KLF activators and repressors is a conserved feature across cortical cell types (**Figure 2A**). We also examined human RNA-seq data from the BrainSpan atlas (Kang et al., 2011) to see if similar trends in KLF expression occur in the human cortex and observed a similar decline in *KLF6* and *KLF7* concurrent with a rise in *KLF9* and *KLF13* expression between birth and the 4^th^ month of postnatal life in four primary sensory cortical regions (**Figure 2B**). This suggests that the developmental regulation of this TF family is a conserved feature of mammalian cortical development.

**Figure 2.**
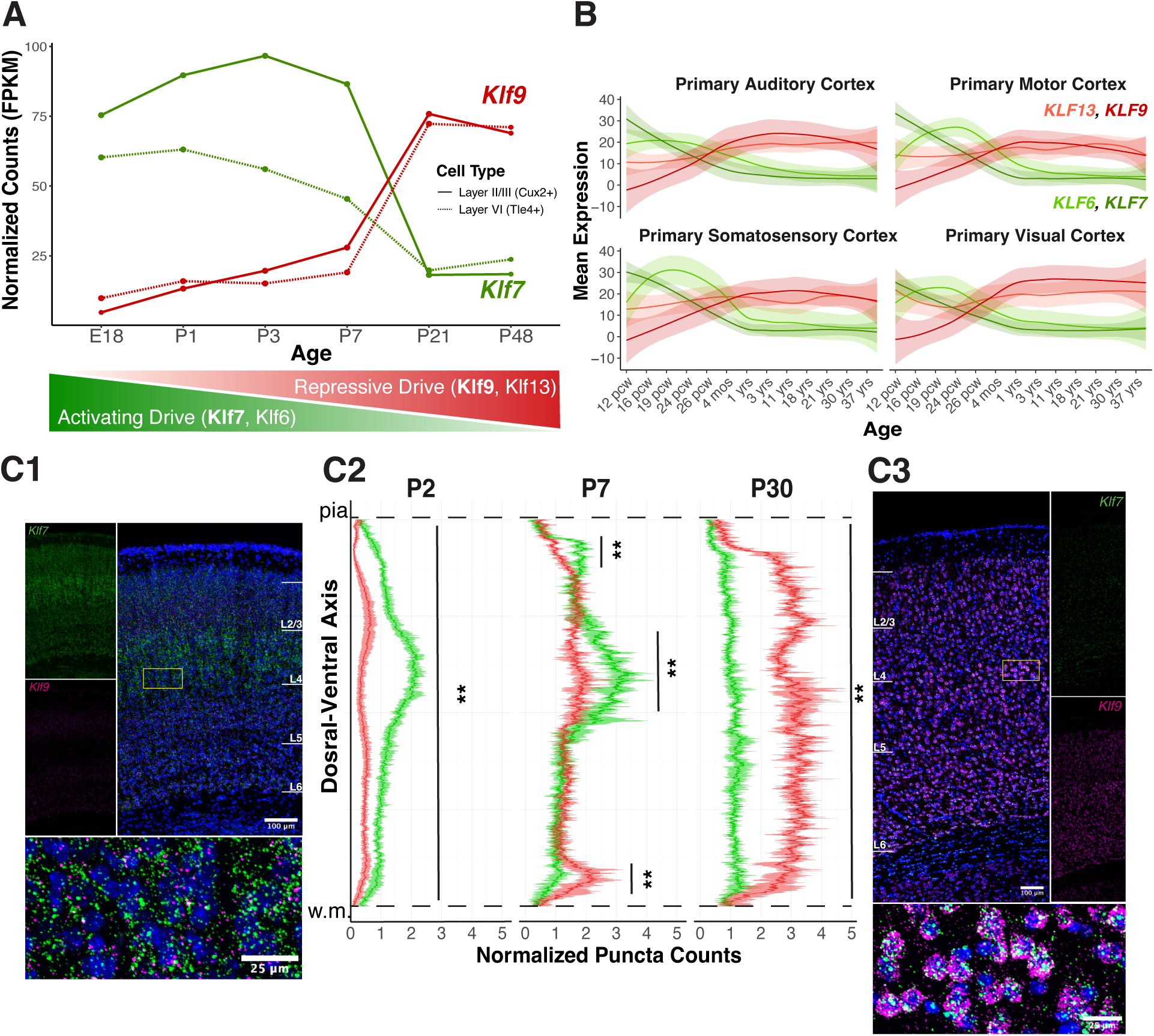
Activating (*Klf6* and *Klf7*) and Repressive (*Klf9* and *Klf13*) members of the KLF family display opposing patterns of gene expression throughout the developing cortex. **A**) Developmental dynamics of *Klf7* and *Klf9* in Layer II/III and Layer VI neurons between E18 and P48. Plotted as mean FPKM from n=2 mice (data replotted from Yuan et al., 2022). **B)** Expression of *Klf6*, *Klf7*, *Klf9*, and *Klf13* mRNA across the human lifespan in 4 sensory cortical regions taken from the human BrainSpan Atlas (Kang et al., 2011). Data are plotted as smoothed loess fits +/- SE. **C**) RNAScope of *Klf7* and *Klf9* mRNA P2, P7, and P30 C57 mouse brains demonstrates *Klf7* mRNA is more abundant than *Klf9* mRNA at P2 (**C1**) while *Klf9* mRNA is more abundant than *Klf7* mRNA at P30, where laminar distribution is more uniform (**C3**). (w.m. = white matter; Scale bar: 100 µm; Insert: 25µm). Data quantified in **C2** (n=3-4 mice per age, plotted as puncta normalized by nuclei density across the dorsal-ventral axis with 95% confidence intervals; **:p<0.001, Mann-Whitney U Test)

We then examined *Klf7* and *Klf9* expression across early postnatal development using Fluorescent *In Situ* Hybridization (FISH) to simultaneously interrogate the temporal and spatial dynamics of the transcriptional switch. As expected, we detected a much higher expression of *Klf7* than *Klf9* mRNA at P2, particularly in Layer 4 (**Figure 2C1**). By P7, both TFs had similar expression across cortical layers, with neurons in Layer 4 maintaining a small but significantly higher level of *Klf7,* potentially due to their unique maturation process <u>(Clark et al., 2020;</u> **Figure 2C2**). At P30 *Klf9* expression had clearly overtaken *Klf7* expression by 1.65–4.39 fold in all cortical layers, and the expression of both KLFs examined appeared uniform across lamina (**Figure 2C3**). Together, these results confirm that members of the KLF family, particularly the activator Klf7 and the repressor Klf9, undergo a coordinated switch in their expression during early postnatal cortical development which coincides with a downregulation of genes enriched for the KLF/Sp motif in their promoters. Notably, the downregulated KLFs (Klf6 and Klf7) are closely structurally related and known to activate transcription of their targets while the upregulated KLFs (Klf9 and Klf13) share similar functional domains and have been primarily described as repressors of gene expression (McConnell & Yang, 2010). We theorized that these opposing expression patterns could enable the KLFs to function as a ‘transcriptional switch’ at a shared set of developmentally downregulated targets through the combined loss of activating drive and increase in repressive drive. Furthermore, this switch could operate without significant changes to the chromatin structure at promoters containing KLF/Sp motifs through a simple competition mechanism between repressors and activators at the same DNA-binding sites.

### Multiplexed Knockdown of *Klf9* and *Klf13 in vivo* using CRISPR Interference

To achieve cell type-specific loss of function of KLF TFs, we crossed homozygous Rosa26-LSL-dCas9-KRAB (CRISPRi) mice to *Emx1*-Cre^+/+^ mice, which generated progeny expressing dCas9-KRAB in excitatory neurons of the forebrain. Using this strategy, cell type-specific knockdown of target transcripts can be achieved through delivery of sgRNA alone. We therefore cloned sgRNA(s) targeting our promoter(s) of interest individually or in tandem into custom AAV transfer vectors also containing a FLOXed fluorescent reporter (mCherry or EGFP) driven by the hSyn promoter to enable reporter expression exclusively in Cre-positive, infected neurons. Transfer vectors were used to generate high-titer rAAV9. To ensure widespread infection, we bilaterally injected neonatal *Emx1*-Cre^+/-^;CRISPRi^+/-^ pups intracerebroventricularly within 12 hours of birth (P0.5), then allowed 18-20 days for viral infection and maximal knockdown (Zheng et al., 2018) before their brains were harvested and processed for purification of reporter-positive neurons by FACS (**Figure 3A**). Cortical infection following neonatal injection was widespread in juvenile mice as previously described (see **Methods**), establishing the technique’s utility for manipulating large populations of neurons with medium throughput (Kim et al., 2013).

**Figure 3.**
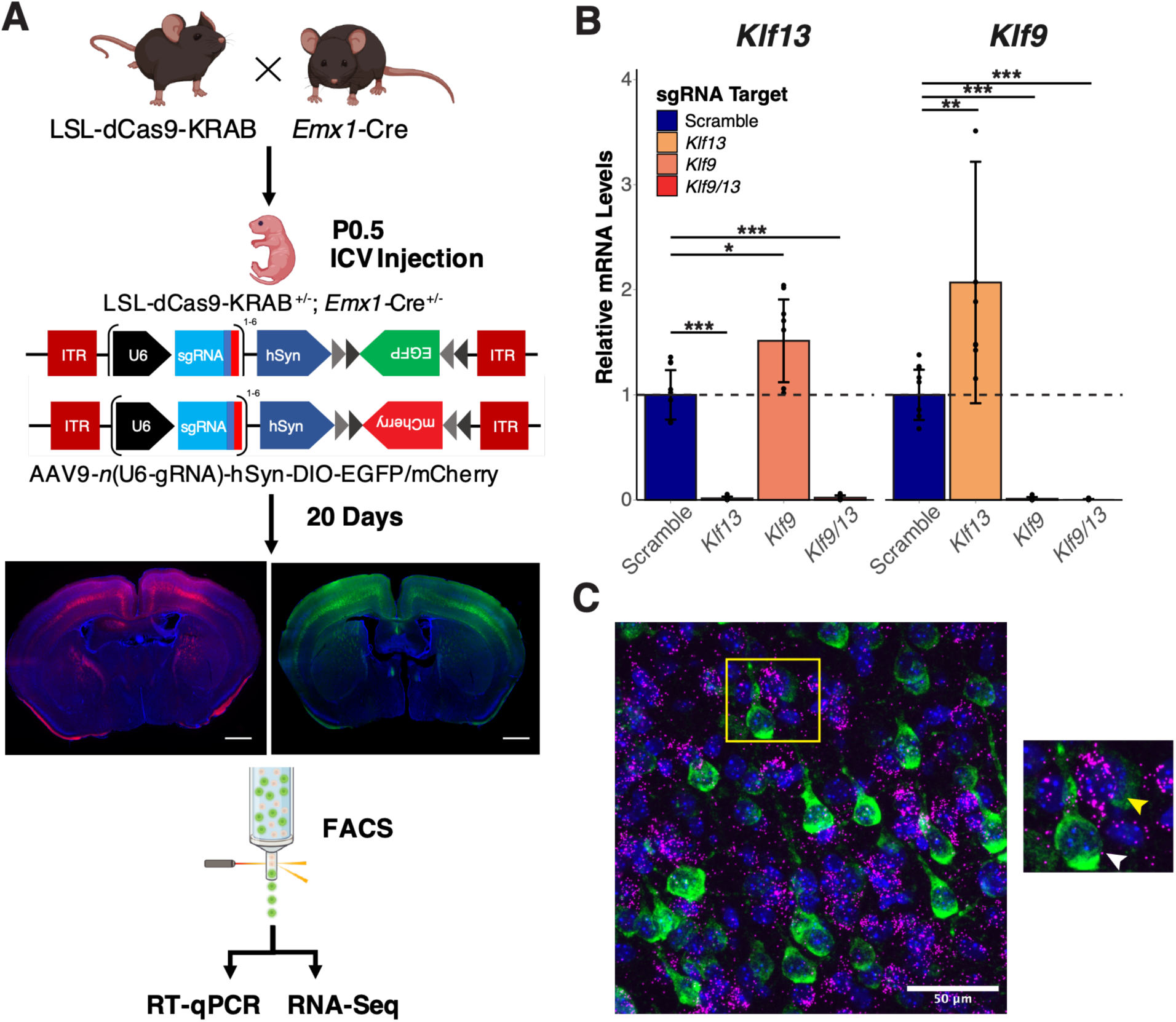
A CRISPR Interference strategy for studying cell-autonomous Transcription Factor knockdown in excitatory cortical neurons. **A)** Schematic of experimental approach: Mice harboring the dCas9-KRAB transgene are crossed with *Emx1*-Cre mice and double heterozygous offspring are intracerebroventricularly injected at Postnatal day 0.5 with AAV9 containing guide RNA’s targeting gene(s) of interest and a floxxed fluorophore. After a 10-20 waiting period, infected neurons are sorted based on fluorescence and processed for RT-qPCR or RNA-seq. (Scale bars = 500µm) **B)** CRISPRi produces consistent and effective knockdown of *Klf9* (99.0±1.7%) and *Klf13* (98.6±1.7%) individually or in combination (*Klf9*: 99.4±0.6%; *Klf13*: 97.9±2.2%) as measured by RT-qPCR (n=6-8 mice/condition; data plotted as average Fold Change relative to Scramble sgRNA Control ± standard deviation, Kruskal-Wallis test with post-hoc pairwise Mann-Whitney U test, * p<0.05 **p<0.01, ***p<0.001) **C)** Example RNA-scope image of P20 mouse cortex infected at P0.5 with AAV9-Klf9^g1,g2^-Klf13^g1,g2^-hSyn-DIO-EGFP and probed for *EGFP* (Green) and *Klf9* (Magenta). (Scale bar = 50µm)

We began by testing this approach against *Klf9*, which is the most abundant KLF in the adult brain and sharply upregulated in excitatory neurons in early postnatal development. We reasoned that this TF was well positioned to effect gene expression in maturing cortex if knocked down prior to its’ rise in expression. We targeted *Klf9* by injecting P0.5 pups with our rAAV9- *Klf9*^sg1,sg2^-hSyn-DIO-mCherry virus (Klf9-mCherry) containing 2 sgRNAs targeting the TSS of *Klf9* to ensure thorough knockdown (Qi et al., 2013). As a control, we injected littermates with rAAV9-Scr^sg1^-hSyn-DIO-mCherry (Scr-mCherry) containing a non-targeting scrambled sgRNA. After 18-20 days, we observed a 99.0±1.7% (P = 0.002) knockdown of *Klf9* by qPCR in neurons from mice injected with the Klf9-mCherry virus relative to mice that received the Scr-mCherry virus (**Figure 3B**).

While Klf9 is the most abundant KLF in the adult mouse brain by far, studies in other tissues suggest that its loss can be partially compensated for by Klf13 (<u>Ávila-Mendoza et al., 2020; Heard et al., 2012;</u> Bernhardt et al., 2022). Because Klf13 exhibits the same developmental upregulation as Klf9 and they are among the most structurally similar KLFs (McConnell & Yang, 2010), we reasoned that a knockdown of both TFs could be more effective at uncovering a transcriptional phenotype. Therefore, we used identical methods to knock down *Klf13* or *Klf9* and *Klf13* simultaneously using AAV9-*Klf13*^sg1,sg2^-hSyn-DIO-mCherry (Klf13-mCherry) or AAV9-*Klf9*^sg1,sg2^-*Klf13*^sg1,sg2^-hSyn-DIO-mCherry (Klf9/13-mCherry) respectively. We detected a 98.6±1.7% (P = 0.0009) knockdown of *Klf13* in mice injected with Klf13-mCherry and a 99.4±0.6% (P = 0.002) knockdown of *Klf9* and a 97.9±2.2% (P = 0.001) knockdown of *Klf13* in mice injected with Klf9/13-mCherry relative to scramble controls, demonstrating the utility of CRISPRi for *in vivo* multiplexed knockdown experiments (**Figure 3B**). Furthermore, we observed a 1.5±0.4-fold (P = 0.025) increase in *Klf13* expression in mice receiving the Klf9-mCherry virus and a similar 2.1±1.2-fold (P = 0.01) increase in *Klf9* expression in Klf13-mCherry samples, suggesting that compensation following loss of a single factor could involve a release of mutual repression. Finally, to confirm that our knockdown was restricted to infected cells, we used RNA-Scope to visualize *Klf9* expression in the brains of P20 mice neonatally injected with an EGFP-expressing rAAV9-*Klf9*^sg1,sg2^-*Klf13*^sg1,sg2^-hSyn-DIO-EGFP virus containing identical sgRNAs as our Klf9/13-mCherry construct. We detected a near-total absence of *Klf9* puncta in EGFP-positive cells interspersed with EGFP-negative nuclei surrounded by dense *Klf9* signal (**Figure 3C**, further quantified in **Figure 8**). This indicates that our manipulation is highly penetrant and restricted to our cells of interest.

### Klf9 and Klf13 are Partially Compensatory Transcriptional Repressors

To identify putative targets of Klf9 and Klf13 and examine the transcriptional consequences of knocking down the major KLF repressors in the juvenile mouse brain, we prepared RNA-seq libraries from sorted neurons receiving sgRNAs targeting *Klf9*, *Klf13*, *Klf9* and *Klf13* or Scrambled controls. Principal Component Analysis of the resultant data showed a clean separation of all treatment groups along PC1 that mirrored the fraction of endogenous *Klf9* + *Klf13* mRNA remaining in these groups relative to the scramble control (*Klf9*/13 > *Klf9 >> Klf13;* **Figure 4A**). Differential Gene Expression analysis revealed 5 DEGs following *Klf13* KD, 28 DEGs following *Klf9* KD, and 212 DEGs in the *Klf9/13* KD samples relative to Scramble controls (DEGs: Protein-coding genes with Absolute Fold-Change³2, Adjusted p. value<0.05; **Figure S4.1A-B**). The gradient observed along PC1 and the discrepancy in DEGs identified among single-versus double-knockdown samples indicates that compensation for loss of Klf9 by Klf13 likely occurs *in vivo*. Volcano plots comparing *Klf9/13* KD samples to Scramble Controls further underscored the efficacy and specificity of CRISPRi knockdown, as *Klf9* and *Klf13* stood out as the most significantly downregulated DEGs (**Figure 4B**). Furthermore, the greater fraction of upregulated DEGs relative to all DEGs following *Klf9/13* KD (85.4%, 181 genes) is consistent with Klf9 and Klf13’s known roles as repressors of gene expression, as upregulated DEGs could plausibly result from a loss of repression.

**Figure 4.**
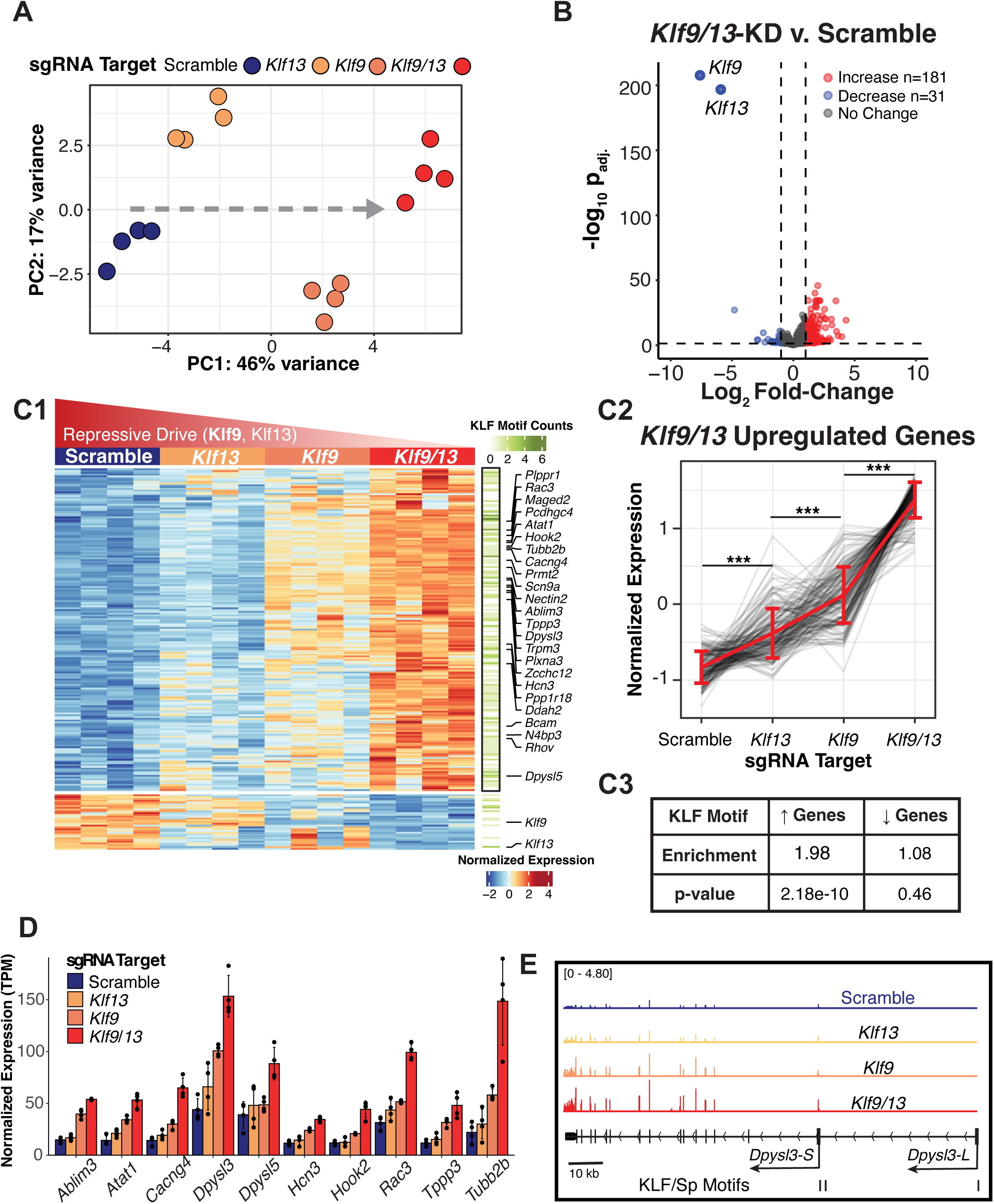
*Klf9* and *Klf13* show compensatory activity and additive regulation of target genes. **A)** Principal component for top 500 most variable genes in all knockdown samples and scramble controls shows graded effect of KLF knockdown along PC1 (n=4 mice per condition). **B)** De-repression of target genes following *Klf9/13* KD. Volcano plot of gene expression changes in *Klf9/13* KD relative to Scramble Control mice (DEGs defined as |log2FoldChange|>1; p.adj<0.05 by Wald’s Test with BH correction). **C1)** Heatmap of all DEGs identified in *Klf9/13* KD (212 genes) across all libraries. Data plotted as z-scored, regularized log-transformed counts. Right bar represents KLF/Sp root cluster motif counts identified in ATAC-seq peaks overlapping respective Promoters (TSS -1000/+200bp) by FIMO (qval<0.05, quantified in **C3,** Fisher’s Exact Test). **C2**: Line plots of all upregulated Klf9/13 targets across all libraries. Grey lines are z-scored, regularized log-transformed RNA-seq counts of individual genes, blue lines represent average of all genes +/- SEM. (***:p<0.0001, Kruskal-Wallis test with post-hoc pairwise Mann-Whitney U test) **D)** Additive regulation of selected targets with roles in cytoskeletal or synaptic function (n=4 mice per group, average TPM +/- standard deviation). **E)** Example IGV traces of RNA-seq coverage at the *Dpysl3* locus. Arrows indicate KLF/Sp motif matches (FIMO, q<0.05).

K-means clustering of *Klf9/13* DEGs across all treatment groups identified 2 clusters representing down- and upregulated gene sets. When visualized as a heatmap, the upregulated gene cluster revealed a gradient in gene expression from Scramble control samples to *Klf9/13* KD samples akin to the gradient observed in PC1 (**Figure 4A**), suggesting that there were subthreshold effects of *Klf9* and *Klf13* KD on *Klf9/13* DEGs (**Figure 4C1**). The increased effect size produced by depleting increasing amounts of KLF repressive drive was particularly evident when we plotted the z-scored expression of the 181 upregulated DEGs across all samples (**Figure 4C2**). Viewed this way, even *Klf13* KD produces a small but highly significant (P = 5.44×10^-26^) effect on Klf9/13 target genes. A similar trend was observed among the 31 downregulated DEGs (**Figure S4.1C**). Together, these results support a model in which Klf9 and Klf13 additively cooperate to repress shared targets and can partially compensate for one another in a dose-dependent manner.

Although Gene Ontology analysis did not return any enriched terms among up- or downregulated gene sets (data not shown), manual inspection of the genes with the greatest ‘KLF Effect’ (absolute loadings along PC1) led to the identification of putative targets with synaptic and cytoskeletal functions as well as ion channels (**Figure 4C1, 4D, 4E**). The abundance of cytoskeletal regulators among targets of Klf9/13 coincides with the established roles of these factors as inhibitors of neurite outgrowth. We did not detect a significant change in *Klf6* or *Klf7* expression in the double knockdown, making it unlikely that the rise in *Klf9* and *Klf13* expression drives the decline in *Klf6* and *Klf7* expression.

The dysregulation of genes related to cytoskeletal regulation and synaptic function prompted us to look for a synaptic phenotype in *Klf9/13* KD cells. However, whole cell patch clamp recordings from infected neurons in *Klf9/13* KD mice and Scramble controls showed no significant difference in miniature Excitatory Postsynaptic Current (mEPSC) amplitude (Klf9/13-mCherry: 12.9±0.614 pA; Scr-mCherry: 12.7±0.47 pA) or frequency (Klf9/13- mCherry: 5.85±0.76 Hz; Scr-mCherry: 5.44±0.20 Hz) between groups (**Figure S4.2A,C**). Intrinsic Excitability of *Klf9/13* KD neurons was also unchanged (**Figure S4.2B**).

Because the KLF family was selected based on enrichment of the KLF/Sp motif in developmental DEG promoters, we reasoned that we should be able to recover this motif enrichment using ATAC-seq peaks overlapping the promoters of DEGs identified from *Klf9/13* KD experiments. Indeed, we found a significant (1.98-fold, P = 2.18×10^-10^) enrichment of the root motif of the cluster containing all KLFs in the JASPAR 2022 CORE collection in promoters of upregulated DEGs but not among downregulated DEGs (p = 0.46, **Figure 4C3**). Because this approach was biased towards the expected motif, we also performed an unbiased motif enrichment analysis on upregulated DEG promoters using the entire JASPAR 2022 CORE collection and found a nearly exclusive enrichment of KLF/Sp family motifs ranging from to 1.47- to 2.76-fold (P adj. << 0.01) (**Figure S4.1D**). These data support the idea that the upregulated genes are likely direct targets of Klf9 and Klf13 and validate CRISPRi as a novel tool to identify TF targets *in vivo*.

### Targets of Klf9 and Klf13 are Developmentally Downregulated Genes

Developmental upregulation of *Klf9* and *Klf13* coincides with downregulation of genes containing an enrichment of the KLF/Sp motif in their promoters. We therefore reasoned that putative targets of Klf9 and Klf13 should themselves be developmentally downregulated. To investigate this, we examined RNA-seq data from sorted Rorb+ Layer 4 neurons published in Clark et al., 2020, which afforded us an additional timepoint between P2 and P30. We observed a clear and significant decrease in the normalized expression of our 181 putative targets of Klf9/13 between P2 and P30, with intermediate expression at P7 (**Figure 5A**). Furthermore, the rise of *Klf9* and *Klf13* during this period is perfectly antagonistic to the decrease in their targets. Plotting genome-wide effects of *Klf9/13* KD against expression changes between P2 and P30 from our original dataset revealed a significant negative correlation (R = -0.56, P < 2.2×10^-16^; **Figure 5B**).

**Figure 5.**
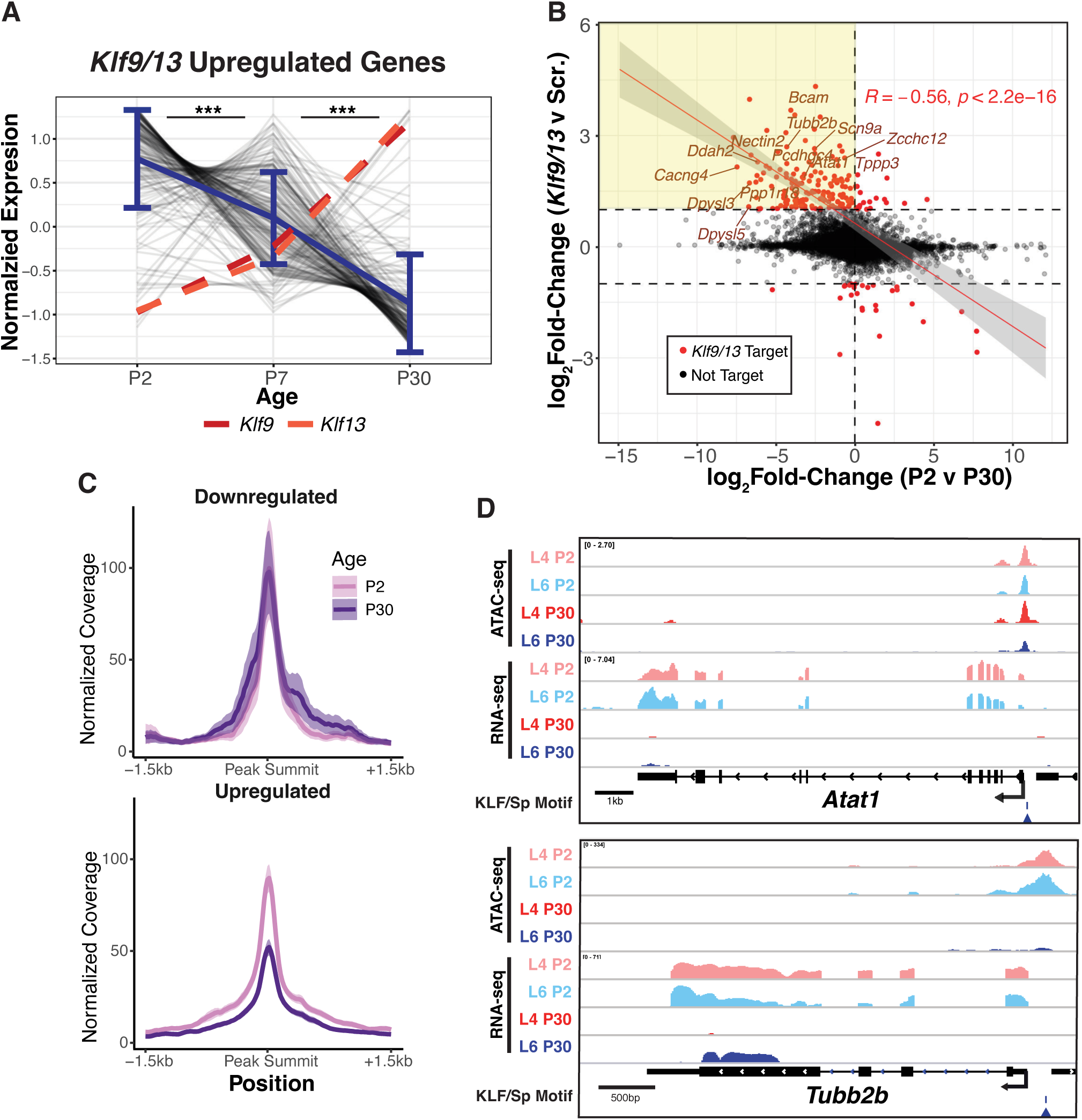
Putative Klf9/13 Targets are developmentally downregulated in the maturing cortex. **A)** Expression of upregulated DEGs in *Klf9/13* KD at early, mid, and late postnatal developmental time points. Grey lines are z-scored, regularized log-transformed RNA-seq counts of individual genes from sorted Rorb+ (Layer 4) excitatory neurons, blue lines represent average of all genes +/- SEM (n=4 mice per group, Kruskal-Wallis test with post-hoc pairwise Mann-Whitney U test, ***p<0.001). **B)** Gene expression changes in *Klf9/13* KD relative to Scramble Controls at P18-20 (n=4 mice per group) relative to gene expression changes between P2 and P30 (n=8 mice per group, aggregate of 2 cell types). Points represent in individual genes, and red points are DEGs identified in *Klf9/13* KD samples. Line is linear fit +/- 95% CI to Klf9/13 DEGs (red). **C)** RiP-normalized coverage of average ATAC-seq signal at Promoter peaks around DEGs identified from *Klf9/13* KD RNA-seq (Shaded region = 95% CI; n=4 libraries per group, aggregate of 2 cell types). **D)** Example IGV traces from P2 & P30 ATAC-seq and RNA-seq for Klf9/13 Targets *Atat1* (**top**, Developmental DEG *without* Promoter DAR) and *Tubb2b* (**bottom**, Developmental DEG *with* Promoter DAR). Arrows indicate putative KLF/Sp binding site identified by FIMO (q<0.05)

Transcriptional repression by Klf9 and Klf13 could occur via direct inhibition of RNA Polymerase II’s binding or activity, or through dynamic changes to the chromatin accessibility at target gene promoters, or through both mechanisms. In support of the latter model, *in vitro* studies on Klf13’s mechanism of action have suggested that it may promote histone modifications leading to heterochromatin formation through its’ Sin3a domain which it shares with Klf9 (Kaczynski et al., 2001). To differentiate between these possibilities, we looked for evidence of changes in chromatin accessibility between P2 and P30 at promoter elements of Klf9/13 targets. Although only a minority (21.6%) of peaks in target promoters met the threshold for DARs, coverage plots of ATAC-seq signal centered on promoter peaks revealed a noticeable loss in accessibility between P2 and P30 in genes upregulated by *Klf9/13* KD (**Figure 5C2**). Furthermore, examination of the relationship between changes in gene expression following *Klf9/13* KD and chromatin accessibility between P2 and P30 revealed a weak negative correlation (R = -0.2) between these variables bolstered by large changes in chromatin accessibility at a minority of highly upregulated Klf9/13 targets (**Figure S5A**). Inspection of ATAC-seq signal at promoters of selected Klf9/13 targets revealed a notable loss of accessibility at promoter peaks containing the KLF/Sp motif upstream of *Tubb2b* and *Dpysl3*, but no change in accessibility in motif-positive peaks upstream of *Plppr1* and *Atat1* (**Figures S5B** and **5D**). These data suggest that a Klf9/13-mediated loss of promoter accessibility could lead to the repression of a subset of targets during early postnatal development, but the mechanism of repression may differ in a target- or site-specific manner.

### Klf6 and Klf7 promote expression of developmentally regulated genes in the perinatal neocortex

The transcriptional activators Klf6 and Klf7 display an inverse pattern in their mRNA and protein expression relative to Klf9 and Klf13, with high prenatal expression that declines gradually after the first week of life. This suggests that Klf6 and Klf7 could function in opposition to Klf9 and Klf13 by promoting the transcription of at least some of same neonatally expressed targets, possibly by binding to the same KLF/Sp motifs in their promoters. To test this possibility, we utilized a similar CRISPRi loss of function approach by creating an AAV construct designed for multiplex knockdown of *Klf6* and *Klf7* (Klf6/7-GFP; AAV9-*Klf6*^sg1,sg2^- *Klf7*^sg1,sg2^-hSyn-DIO-EGFP). We elected to pursue a multiplex KD strategy from the outset following our observation that *Klf9/13* knockdowns have reduced penetrance due to redundancy. Klf6 and Klf7 have highly homologous protein sequence, can compensate for one another in zebrafish, and have non-additive effects on neurite branching *in vitro* (Moore et al., 2009; Veldman et al., 2007).

As before, we bilaterally injected P0.5 offspring of CRISPRi and Emx1-Cre mice with targeting rAAV9 or a scramble control (Scr-GFP) before sorting and collecting mRNA from infected neurons. Because expression of KLF activators is highest in early development and their targets might already be downregulated by KLF repressors by P18-20 (P20), we also sorted neurons from P10-12 mice (P10). This was the minimum latency of infection we deemed appropriate given the time required for mRNA and protein turnover following repression by dCas9-KRAB. We found a near-complete knockdown of both *Klf6* and *Klf7* at P10 (*Klf6*: 99.6±0.5%, P = 0.0005; *Klf7*: 97.6±3.1%, P = 0.0006) and at P20 (*Klf6*: 99.8±0.1%, P =0.0005; *Klf7*: 94.7±12.6%, P = 0.004) relative to scramble controls, further demonstrating the general applicability of CRISPRi for manipulation of a range of targets (**Figure 6A**). In addition, we were able to detect a 2.76-fold (P = 0.0007) reduction in *Klf6* and 2.89-fold (P = 0.0009) reduction in *Klf7* expression between P10 and P20, indicating that the KLFs are still undergoing their switch during this period.

**Figure 6.**
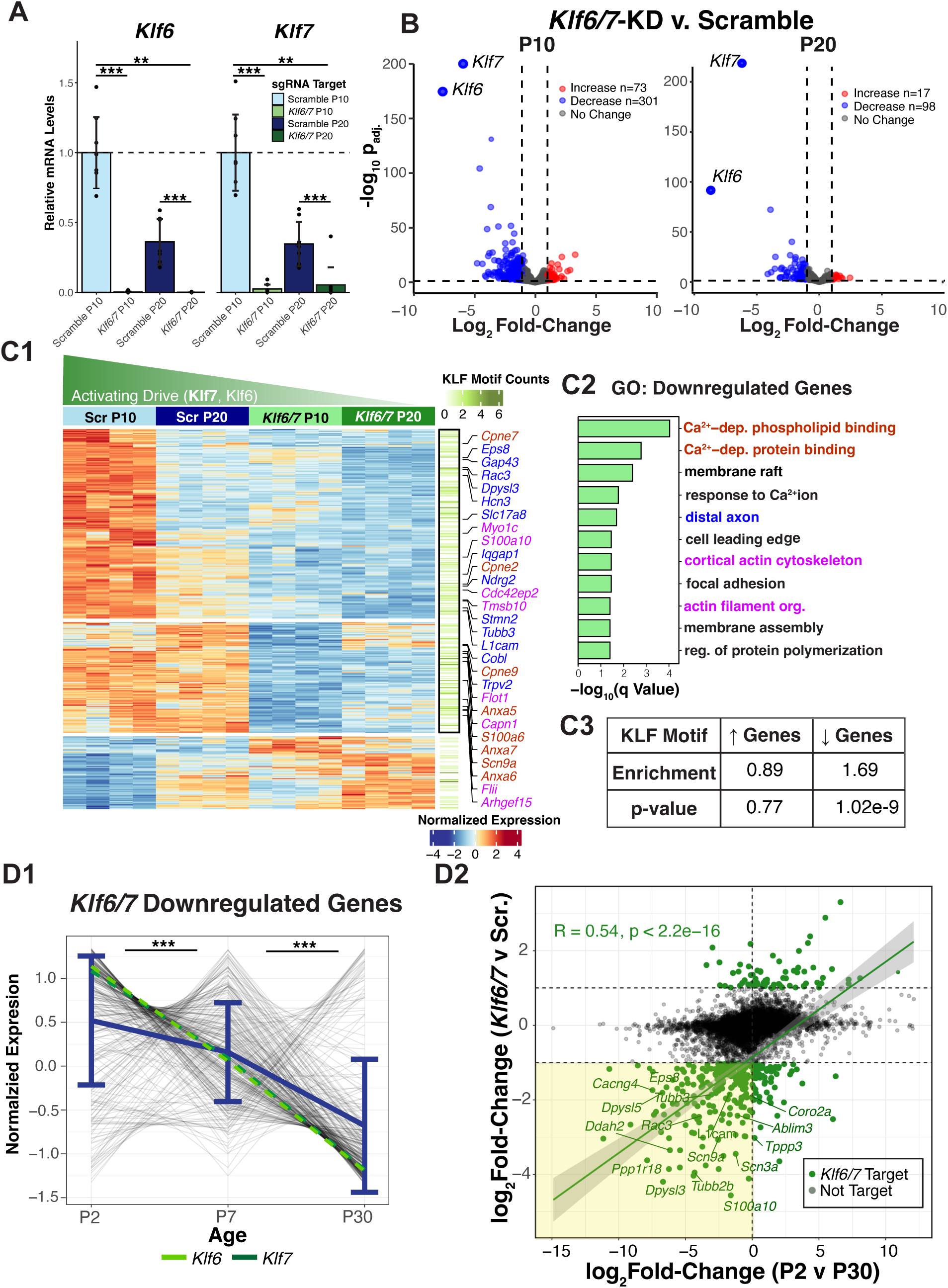
*Klf6* and *Klf7* Promote expression of Developmentally Regulated Genes in the Perinatal Cortex. **A)** RT-qPCR quantification of *Klf6* and *Klf7* in mice injected with *Klf6* and *Klf7*-targeting sgRNA’s at P0, collected at early (P10-12) or late (P18-20) postnatal timepoints (n=7-9; data plotted as average fold-change relative to P10 Scramble control +/- standard deviation, Kruskal-Wallis test with post-hoc pairwise Mann-Whitney U test with BH adjustment, **p<0.005, ***p<0.001) **B)** Volcano plot of DEGs identified in *Klf6/7* KD samples relative to Scramble Controls at P10 (**B1**) and P20 (**B2,** DEGs defined as |log2FoldChange|>1; p.adj<0.05 by Wald’s Test with BH correction). **C1)** Heatmap of all DEGs identified in *Klf6/7* KD at P10 (374 genes, k=3). Data plotted as z-scored, regularized log-transformed counts. Right bar represents KLF motif counts identified in ATAC-seq peaks overlapping respective Promoters (TSS -1000/+200bp) by FIMO (qval<0.05, quantified in **C3,** Fisher’s Exact Test). Gene Ontology overrepresentation analysis of downregulated targets shown in **C2.** Gene names in **C1** colored by GO term membership. **D)** Developmental regulation of Klf6/7 Targets **D1:** Expression of downreguated DEGs in *Klf6/7* KD at early, mid, and late postnatal developmental time points. Grey lines are z-scored, regularized log-transformed RNA-seq counts of individual genes from sorted Rorb+ (Layer 4) excitatory neurons, blue lines represent average of all genes +/- SEM (n=4 mice per group, Kruskal-Wallis test with post-hoc pairwise Mann-Whitney U test, ***p<0.001). **D2:** Gene expression changes in *Klf6/7* KD relative to Scramble Controls at P10-12 (n=4 mice per group) relative to gene expression changes between P2 and P30 (n=8 mice per group, aggregate of 2 cell types). Points represent in individual genes, and green points are DEGs identified in *Klf6/7* KD samples. Lines are linear fits +/- 95% CI to all points (black) or Klf6/7 DEGs (green) with correlation indicated in the respective colors.

We next conducted RNA-sequencing of P10 and P20 excitatory neurons infected Klf6/7- GFP and Scr-GFP AAV with the goals of identifying likely targets of the KLF activators and examining how their regulatory role changes across early postnatal development. Using identical criteria as above, we identified 374 DEGs in P10 *Klf6/7* KD neurons and 115 DEGs at P20 relative to age-matched scramble controls, with both sgRNA targets once again being the most significantly affected DEGs (**Figure 6B**). In accordance with Klf6 and Klf7’s reported functions as transcriptional activators, the majority of DEGs were downregulated at both ages (80.5% at P10, 85.2% at P20). Furthermore, the enhanced effect of *Klf6/7* KD at P10 is consistent with an elevated KLF activating drive at this age resulting from higher expression of Klf6 and Klf7. The elevated Klf9/13 repressive drive at P20 could have repressed Klf6/7 targets that would otherwise have exhibited loss of activation in the KDs as at P10. Therefore, downstream analysis focused on the 374 DEGs identified at P10 when the role of Klf6 and Klf7 is most pronounced.

The inclusion of an additional developmental time point allowed us to ask how Klf6 and Klf7’s regulatory activity and the expression of their putative targets may differ between ages. To identify DEG sets with different trajectories across groups, we performed k-means clustering of our 374 DEGs and visualized the results as a heatmap. We detected 3 clusters: downregulated DEGs that are also developmentally downregulated between P10 and P20, downregulated DEGs that do not change between P10 and P20, and genes upregulated by *Klf6/7* KD (**Figure 6C1**). Notably, the largest cluster of 190 DEGs downregulated by age and *Klf6/7* KD displayed a gradient from high expression at P10 in Scramble control neurons to low expression at P20 in *Klf6/7* KD neurons, approximating the ratio of KLF activator to repressor abundance across groups (**Figure 6C1**). The upregulated gene cluster exhibited the opposite trend, and P20 targets were largely consistent with those identified at P10.

Gene Ontology analysis of all downregulated DEGs identified enrichment for functions related to neurite growth and cell motility such as actin filament organization (q-value = 0.040), membrane rafts (0.004), and focal adhesion (0.035), consistent with previously described roles for Klf6 and Klf7 *in vivo* (**Figure 6C1, 6C2**) (Blackmore et al., 2012; Hong et al., 2023; Kajimura et al., 2007; Wang et al., 2018). Genes contributing to these terms include the actin nucleator *Cobl,* the actin-binding proteins *Ablim3, Cd44, Dpysl3,* and *Iqgap1,* the actin capping protein *Eps8*, the growth-cone enriched microtubule destabilizing factor *Stmn2* (STG10) tubulin isoforms *Tubb3*, *Tubb5*, *Tuba1a*, and *Tubb2b*, and the myosin motor *Myo1c*. Most notably, putative Klf6/7 targets were enriched for genes associated with the distal axon (q-value = 0.020), which includes previously identified targets of Klf7 such as *Rac3*, *Stmn2, Gap43*, *Cdkn1a* (p21), and *L1cam.* This indicates that CRISPRi can reproduce molecular phenotypes of null mutants *in vivo* (Hong et al., 2023; Kajimura et al., 2007).

If DEGs identified by RNA-seq are enriched for direct targets of Klf6/7, we would expect to see an enrichment of the KLF/Sp motif in ATAC-seq peaks overlapping the Promoters of DEGs. Indeed, we found the KLF cluster root motif was 1.69-fold enriched in downregulated gene promoters (p = 1.02×10^-9^) but not in upregulated gene promoters (p = 0.77), consistent with logic that direct targets of transcriptional activators should be downregulated following their KD (**Figure 6C3**). Enrichment analysis for the full JASPAR 2022 CORE motif set also demonstrated a nearly exclusive enrichment for the KLF/Sp motif in downregulated DEG promoters ranging from 1.35- to 2.25-fold (p adj. << 0.01, **Figure S6A**).

*Klf6* and *Klf7* expression declines during early postnatal development, so we wondered if putative targets of these TFs change in parallel as was the case for targets of Klf9/13. We examined changes in gene expression at P10 caused by *Klf6/7* KD against developmental gene expression changes between P2 and P30. This revealed a strong positive correlation (R = 0.54, P < 2.2×10^-16^) among putative targets of Klf6/7, suggesting that Klf6 and Klf7 promote the expression of a subset of neonatally expressed genes that are downregulated by a coordinated developmental loss of KLF activators and phenocopied by their acute KD (**Figure 6D2**). Normalized trend line plots of downregulated DEGs at P2, P7, and P30 also demonstrated significant and progressive decline of putative target expression that parallels reduced levels of *Klf6* and *Klf7* (**Figure 6D1**). Since targets of Klf6/7 are developmentally downregulated, we wondered if they also exhibit changes in promoter accessibility between P2 and P30 that coincide with this change. As with targets of Klf9/13, we found a modest change in overall accessibility at promoter elements in Klf6/7 targets, although only a small percentage (14.8%) of targets contained a promoter DAR (**Figure S6B,C**). Globally, the developmental change in promoter accessibly was weakly but positively correlated with the effect of *Klf6/7* KD on gene expression (R = 0.17), and this effect was most prominent among the most downregulated genes (**Figure S6C**). Together, these data suggest that Klf6 and Klf7 promote neonatal expression of cytoskeletal genes that are repressed in maturing neurons through both chromatin accessibility-dependent and -independent mechanisms.

### KLF Activators and Repressors Bidirectionally Regulate a set of Shared Targets

If the KLF activators and repressors truly act as a transcriptional switch at a subset of targets, they should share the following features: they should be upregulated following *Klf9/13* knockdown, downregulated following *Klf6/7* knockdown, and developmentally downregulated in accordance with the shift in from KLF activator to repressor drive across early postnasal development. As a group, these genes would be expected to be enriched in genes that regulate axonal development or cytoskeletal function and should contain an overrepresentation of the KLF/Sp motif in their promoter.

Inspection of all RNA-seq libraries by Principal Component Analysis revealed a clean separation of all groups sequenced along the first two Principal Components (PCs) corresponding to Age and sgRNA Targets respectively. Intriguingly, *Klf6/7* KD samples and *Klf9/13* KD samples were found at opposite extremes of PC2 (‘KLF Effect’) with Scramble controls in the middle, indicating that genes contributing to this PC were bidirectionally affected by KLF KD (**Figure 7A**). Furthermore, we observed a noticeable shift along PC1 (‘Age Effect’) depending on the sgRNA targets, with *Klf9/13* KD libraries shifted left towards the P10 samples, and *Klf6/7* KD samples shifted right past P20 libraries (**Figure 7A**). This aligns with our hypothesis that the KLF family regulates a developmental gene expression program through an activator-repressor balance: depleting the late-expressed repressors Klf9 and Klf13 returns neurons to a younger transcriptional state while knocking down early-expressed activators Klf6 and Klf7 transcriptionally ‘ages’ these same cells.

**Figure 7.**
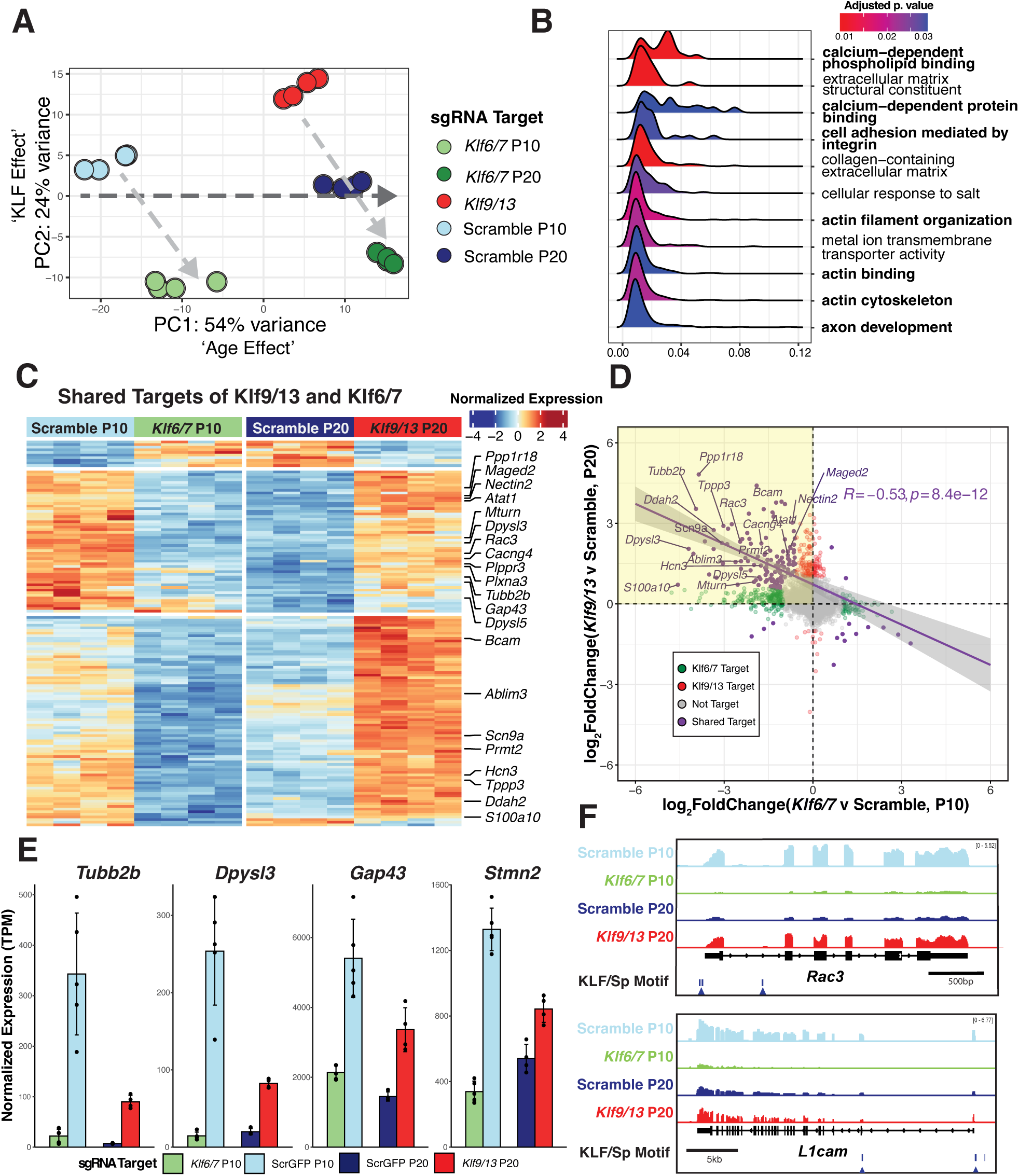
The transition from KLF activators to repressors constitutes a developmental switch at shared targets. **A)** Principal component analysis of all RNA-seq libraries shows that Age and KLF valence account for >75% of variance (n=4 mice each). **B)** GSEA of genes ranked by the bidirectional effect of KLF KD (defined as absolute loadings of PC2 from A) **C** Heatmap of all DEGs shared between *Klf9/13* KD and *Klf6/7* KD ( |log2FoldChange|>1; p.adj<0.05) shows bidirectional regulation by the KLF family and developmental regulation of all shared targets. Data plotted as z-scored, regularized log-transformed counts. **D**) Gene expression changes in *Klf6/7* KD relative to Scramble Controls at P10 are significantly negatively correlated with gene expression changes in *Klf9/13* KD relative to Scramble controls at P20, especially at shared DEGs (p.adj<0.05 by Wald’s Test with BH correction). **E)** Bidirectional regulation of novel (*Dpysl3*, *Tubb2b*), and established (*Gap43*, *Stmn2*) targets Klf7. (Data plotted as TPM +/- standard deviation, n=4 mice) **F)** Representative IGV tracks of RNA-seq data at targets of Klf7 *Rac3* (top) and *L1cam* (bottom) across treatment groups. Arrows indicate the presence of a KLF/Sp motif identified by FIMO (q<0.05).

To identify shared targets of our KLF activator and repressor pairs, we integrated Differential Gene Expression Analysis results from *Klf9/13* KD neurons at P20 and *Klf6/7* KD neurons at P10 compared to their respective age-matched scramble controls. Separate ages were selected to represent the developmental periods when KLF activator drive (P10) or repressive drive (P20) is higher. We found that 41.7% (156/374 DEGs) of putative Klf6/7 P10 targets are significantly affected by *Klf9/13* KD (p adj. < 0.05), with the vast majority (96.1%, 150/156) being oppositely regulated by KLF pairs. Similarly, we found that 47.4% of Klf9/13 P20 targets exhibit significant changes following *Klf6/7* KD, 97.4% of which were bidirectional effects. We refined this to a set of 144 high-confidence shared targets affected by manipulation of both KLF paralog pairs out of 553 total DEGs between them (26.0%; Shared Targets: protein-coding genes with an absolute Fold Change ³ 2 in at least one comparison and ³ 1.5 in the other, adj. p value < 0.05 in both). K-means clustering (k=3) demonstrated that 141/144 (97.9%) of genes meeting these criteria were bidirectionally regulated by the KLFs and the vast majority (134, 93.1%) were upregulated by *Klf9/13* KD and downregulated by *Klf6/7* KD (**Figure 7C**). In addition, most shared targets were downregulated 1.5-fold between P2 and P30 in both Layer 4 and Layer 6 (72.2 and 69.4% respectively), indicating that KLF targets are components of the shared developmental gene regulatory program described above (**Figure S7.1, S7.3**). Notably, knockdown of *Klf9/13* returned expression of shared targets to levels more similar to their expression at P10 in Scramble controls, while *Klf6/7* KD accelerated the developmental decline in these genes to levels closer to those measured at P20 (**Figure S7.2B**). The fact that KD of *Klf9/13* did not fully restore expression of many genes to P10 levels by P20 is consistent with push-pull regulation by developmentally increasing levels of repressors and decreasing levels of activators, although other repressing factors were not ruled out.

In further support of the view that these KLF paralog pairs make up a bidirectional transcriptional switch, we detected a significant negative correlation (R = -0.53, P = 8.4×10^-12^) between effects of *Klf6/7* KD and *Klf9/13* KD at shared targets (**Figure 7D**). Furthermore, the scarcity of targets affected in the same direction by both KD conditions suggests that there are few transcriptional effects attributable to multiplex knockdown alone. As before, we looked for enrichment of the core KLF/Sp motif in accessible promoter regions of shared targets. Reassuringly, we found a significant (2.23-fold, p = 1.12×10^-10^) enrichment of the cognate motif, higher than detected for either *Klf9/13* or *Klf6/7* KD alone. This enrichment was only observed for genes downregulated by *Klf6/7* KD and upregulated by *Klf9/13* KD. Furthermore, motifs recognized by KLF/Sp family TFs were the only overrepresented motifs identified in promoters of shared targets in when this analysis was conducted using the full JASPAR 2022 Core Vertebrate motif collection (2.01- to 3.55-fold enrichment, p adj. < 2.74×10^-6^; **Figure S7.2A**). The magnitude of this effect may indicate that putative shared targets are more likely to represent genuine direct targets of the KLF family than DEGs identified by either knockdown individually. Next, we looked at changes in the accessibility of promoter peaks upstream of shared targets using our developmental ATAC-seq data. We detected a notable loss of ATAC-seq signal at shared KLF targets that was less apparent in non-DEG control genes with similar developmental expression (Figure **S7.2C**).

Although Gene Ontology analysis of shared targets failed to identify any enriched terms, we conducted the threshold-free Gene Set Enrichment Analysis (GSEA) using the absolute loadings of genes along the PCA axis that separated *Klf9/13* from *Klf6/7* KD libraries (PC2). We reasoned that genes contributing to this axis exhibit the strongest bidirectional regulation by the KLFs. GSEA identified enrichment of terms identified among Klf6/7 targets and among shared downregulated genes identified between P2 and P30, including Actin Filament Organization (q-value = 0.017) and Axon Development (q-value = 0.037) (**Figure 7B**). Shared targets with known cytoskeletal and axonal roles were among those with the largest bidirectional effect sizes. Two of the most potently regulated genes in either condition were the actin and microtubule-polymerizing factor *Dpysl3* and the β-tubulin subunit *Tubb2b* with respective roles in Autism Spectrum Disorder (ASD) and polymicrogyria (Stottmann et al., 2013; Tsutiya et al., 2017a) (**Figure 7E**). Other notable shared targets included the actin regulators *Ppp1r18, Prmt2, Rac3,* and *Ablim3*, the tubulin polymerizing protein *Tppp3*, the PTEN inhibitor *Plppr3*, axon guidance mediator *Plxna3* (PlexinA3), inhibitor of microtubule polymerization *Dpysl5*, and the microtubule acetylase *Atat1.* We also identified *Gap43*, a common growth cone marker, strong promoter of axon regeneration and known target of Klf7, as a shared target (Bomze et al., 2001; Kajimura et al., 2007; Skene et al., 1986) (**Figure 7E**).

We also identified significant but subthreshold upregulation (p adj. 0.05; -log_2_(1.5) < LFC < 0) of known Klf6/7 targets and regulators of axon growth *L1Cam*, *Cdkna1*, and *Stmn2* following *Klf9/13* KD, indicating that the list of *bona fide* shared targets may be more extensive (**Figure 7E** and **7F**). A full list of functionally relevant targets of interest is available in **Supplementary Table 1**.

### Bidirectional Regulation of *Dpysl3* and *Tubb2b* by the KLF Family

Because *Tubb2b* and *Dpysl3* were among the most potently bidirectionally regulated genes identified and represent novel disease-relevant targets through which KLFs could exert opposing effects on axon growth, we evaluated effects of KLF KD on their expression using an orthogonal method: RNA FISH. This technique also provides cellular resolution not available when studying transcriptional regulation by bulk RNA-seq. Therefore, to confirm that KLF knockdown and downstream gene expression changes are occurring in the same cells, we used RNAScope to visualize the knockdown of *Klf9* and upregulation *Dpysl3* and *Tubb2b* following neonatal knockdown of *Klf9* and *Klf13* via CRISPRi (**Figures 8A** and **S8A**). As expected, we found a near-complete absence of *Klf9* puncta in GFP+ cells from P20 *Klf9/13* KD mice (Scramble: 64.54±4.43 puncta/cell; Klf9/13: 2.60±1.02). In the same infected neurons, *Tubb2b* and *Dpysl3* were potently upregulated in a manner consistent with changes observed by RNA-seq (**Figure 8B**; *Dpysl3*: Scramble: 18.65±1.98 puncta/cell; Klf9/13: 39.21±1.10; *Tubb2b*: Scr: 6.23±0.29 puncta/cell; Klf9/13: 28.11±2.23). This confirms that *Klf9* KD and *Dpysl3* and *Tubb2b* upregulation are occurring within the same cells, likely as a direct effect of the loss of repression by Klf9 and Klf13.

**Figure 8.**
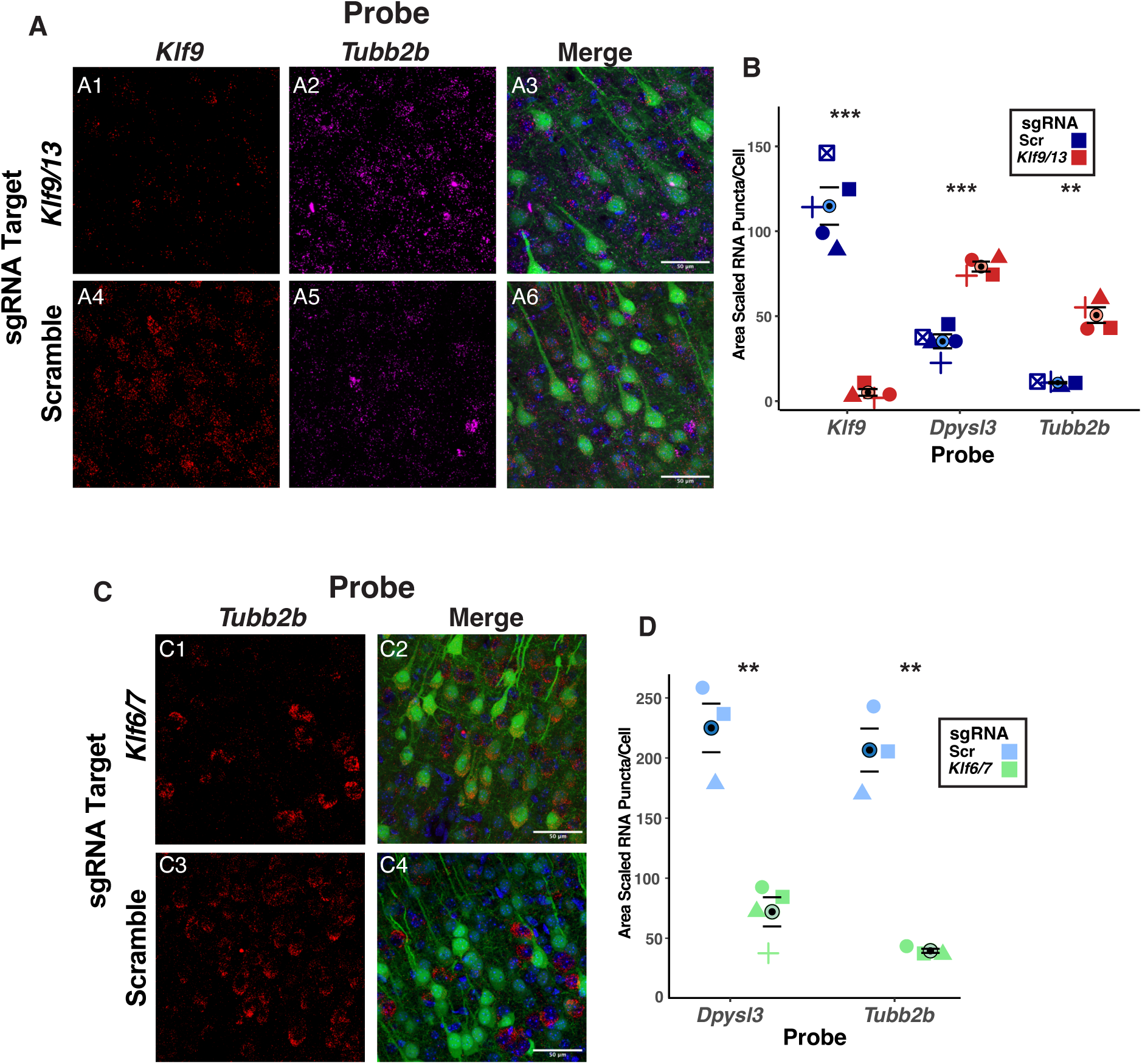
FISH validation of bidirectional regulation of *Tubb2b* and *Dpyls3* by KLF activators and repressors. **A)** Maximum intensity projections of slices taken from P20 mice receiving P0.5 i.c.v. injections of Scramble sgRNA or *Klf9/13*-targeting sgRNA AAV9 probed for *Klf9* (**A1,A4**) and *Tubb2b* (**A2,A5**). Infected cells receiving sgRNA are EGFP-positive (**A3,A6**). Scale bars: 50µm. **B**) Quantification of results in **A**. Data plotted as average number of RNA puncta per cell scaled by cell pixel area ± s.e.m. (n=4 mice per condition, n=5 for ScrGFP *Dpysl3* and *Klf9;* ***: p<0.001, **: p<0.005 independent samples t-test). **C)** Maximum intensity projections of slices taken from P10 mice receiving i.c.v. injections of Scramble sgRNA or *Klf6/7*-targeting sgRNA AAV9 at P0.5 probed for *Tubb2b* (**C1,C3**). Infected cells receiving sgRNA are EGFP-positive (**C2,C4**). Scale bars: 50µm. **D**) Quantification of results in **C**. (n=3 mice per condition, n=4 for Klf6/7 *Dpysl3;* **: p<0.01 independent samples t-test). Shaded shapes represent individual animals.

Since *Tubb2b* and *Dpysl3* are also predicted targets of Klf6/7, we next investigated whether we could detect their downregulation by RNAScope following *Klf6/7* KD. We therefore probed P10 slices from mice injected at P0.5 with our Klf6/7-GFP KD virus or Scr-GFP control for *Dpysl3* and *Tubb2b* (**Figures 8C** and **S8B**). As expected, we saw significant reductions in *Dpysl3* and *Tubb2b* RNA puncta in neurons infected with *Klf6/7* KD virus relative to scramble controls (**Figure 8D**; *Dpysl3*: Scramble: 99.39±7.32 puncta/cell; Klf6/7: 24.75±2.06; *Tubb2b*: Scr: 106.72±7.14 puncta/cell; Klf6/7: 16.77±0.84). Qualitative comparison of the relative *Dpysl3* and *Tubb2b* abundance in Scramble control neurons between P10 and P20 further validates that a dramatic downregulation of *Tubb2b* and *Dpysl3* occurs during early postnatal development, concomitant with the switch in KLF activator and repressor drive. Together, these results validate our RNA-seq findings *in situ* and further argue that bidirectional *Tubb2b* and *Dpysl3* regulation is a direct product of the balance between KLF activators and repressors.

## Discussion

Here, we utilize a multiplexed CRISPRi approach to demonstrate that four paralogs from a diverse TF family function redundantly and antagonistically regulate a common set of targets during early postnatal cortical development. Specifically, we show that Klf9 and Klf13 compensate for one another in a concentration-dependent manner to repress a set of neonatally expressed genes. We provide evidence that this repression functions in opposition to Klf6 and Klf7, which promote the expression of an overlapping set of genes in early postnatal development. This co-regulated gene set is enriched for genes with axonal and cytoskeletal functions, consistent with the known roles of KLF paralogs in regulating axon growth (Moore et al., 2009). We propose a model whereby the KLF family transitions from an activating mode of action driven by high Klf6 and Klf7 expression and low Klf9 and Klf13 expression to a repressive mode of action driven by high Klf9/13 and low Klf6/7 during the second to third postnatal week (**Figure 9**). The temporal alignment of this transition with the conclusion of the postnatal period for cortical neuron axon growth suggests that the shift in the KLF family’s regulatory valence could contribute to the termination of this neonatal program (Fenlon & Richards, 2015; Schreyer & Jones, 1982). Finally, we offer the first demonstration of CRISPRi as a tool for identifying novel targets of TFs in the mammalian brain *in vivo* and demonstrate its potential for studying and overcoming redundancy in paralogous gene families.

**Figure 9.**
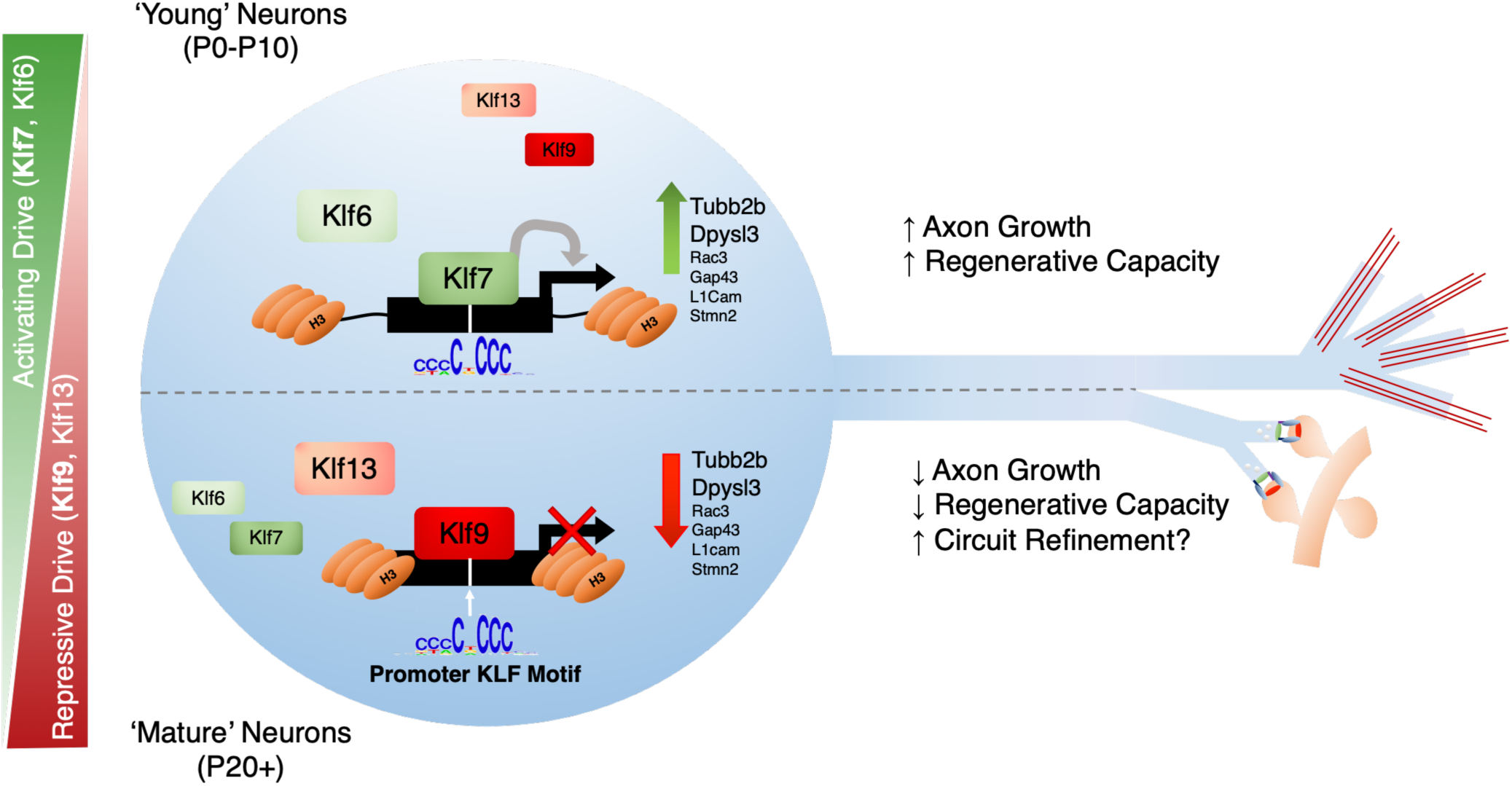
A model for the postnatal loss of axon growth capacity through competition between KLF paralogs. During the first week of postnatal development when Klf6 and Klf7 expression are high, the KLF family functions to activate transcription of genes with roles in axon growth including (but not limited to) *Tubb2b*, *Dplys3*, *Rac3*, and *Gap43* by binding GC-rich regions in target gene promoters (Top). During the second and third week of postnatal development, the expression of transcriptional repressors Klf9 and Klf13 increases while Klf6 and Klf7 expression decreases, leading to displacement of KLF activators at target promoters and repression of pro-growth transcripts (bottom)

Our focus on shared transcriptional features of postnatal cortical development stems from an appreciation for the conservation of this program across species and cell types since clusters representing shared changes in gene expression and chromatin accessibility represent nearly 50% of all DEGs and DARs, especially at Promoters (Bakken et al., 2016; Shi et al., 2021; Yuan et al., 2022; Zhu et al., 2018). That the primary axis of variance for both gene expression and chromatin accessibility was associated with age and not cell type suggests the primacy of a shared developmental program, an observation that has been made in some recent studies (Mayer et al., 2018; Yuan et al., 2022). Other studies of chromatin accessibility, gene expression, and DNA methylation in the prenatal and adult brain have emphasized how cellular diversity is achieved (Di Bella et al., 2021; Liu et al., 2021; Yao et al., 2023; Zu et al., 2023), a compelling transcriptional explanation for the uniformity of the physiological and morphological transitions exhibited by cortical neurons across species and layers is comparatively lacking.

Analysis of TFBSs associated with shared DARs and DEGs generated several predictions for TFs involved in neuronal development. Specifically, the ubiquity of AP-1 motif enrichment in promoters of upregulated DEGs and gained DARs of all classes examined is reminiscent of a recently described role for FOS in directing activity-dependent enhancer formation in maturing interneurons (Stroud et al., 2020). FOS is necessary to facilitate lasting activity-induced chromatin accessibility changes in the hippocampal formation, and the AP-1 motif was recently detected in ATAC-seq peaks gained during postnatal maturation of human and mouse neurons *in vivo* and *in vitro* (Ciceri et al., 2024; Patel et al., 2022). Additionally, the ability of AP-1 TFs to serve as pioneer-like factors could prime maturing neurons for activity-dependent gene expression changes by increasing chromatin accessibility in promoters of genes with synaptic functions (Malik et al., 2014; Stroud et al., 2020; Su et al., 2017; Vierbuchen et al., 2017).

The hypothesized role for the KLF family as regulators of a shared cortical maturation program was motivated by three lines of convergent evidence. First, we noticed a striking correspondence between the biological functions enriched among shared downregulated genes (axon development, cell projection organization) and the known or purported roles of KLF paralogs predicted to interact with their promoters (**Figure 1B3**) (Moore et al., 2009). Second, enrichment of the highly conserved KLF/Sp motif in promoters was consistent with established preferences of Klf9, Klf13, and Klf7 for binding TSS-adjacent regions (Ávila-Mendoza et al., 2020; Knoedler et al., 2017; Tian et al., 2022). We reasoned that promoter elements were a rational site to look for TFs regulating the shared maturation program given their relative lack of cell type-specific patterns of accessibility (Gray et al., 2017; Mo et al., 2015). Finally, the developmentally regulated expression of KLF paralogs (**Figure 2**) suggested they could regulate gene expression through coordinated changes in TF concentration.

Our data and others indicate that the switch in KLF paralog composition over postnatal development is a general and evolutionarily conserved feature of pyramidal neurons (Ávila-Mendoza et al., 2024). While the magnitude and delay of this switch appear to exhibit slight variability across cortical layers, its ubiquity suggests a general role in mammalian cortical development. The upstream signal driving the developmental increase in KLF repressors has yet to be determined, but there is intriguing evidence supporting a role for Thyroid Hormone (T3). Klf9 was originally characterized as a T3-responsive TF in the rat hippocampus, and its’ developmental increase temporally aligns with the postnatal rise in circulating T3 levels (Denver et al., 1999; Denver & Williamson, 2009). Notably, in Purkinje Cells of the cerebellum, T3-driven upregulation of *Klf9* is necessary and sufficient to mediate the inhibitory effects of T3 on axon growth (Avci et al., 2012). This mechanistic link between T3, Klf9, and axon growth is further supported by the excessive axon growth phenotype observed in hypothyroid rats that presumably fail to express Klf9 (Berbel et al., 1993; Li et al., 1995). In support of this, analysis of recently published snRNA-seq data of adult mice treated with T3 (Hochbaum et al., 2024) revealed that *Klf9* is robustly upregulated by T3 treatment in all annotated excitatory cortical cells type clusters, while select shared KLF targets including *Gap43* and *Dpysl3* are reciprocally downregulated in clusters with the greatest upregulation of *Klf9* (**Figure S9**).

A T3-dependent postnatal increase in Klf9 and Klf13 expression with equivalent timing has been reported in oligodendrocyte precursor cells, and this appears to regulate their differentiation into myelinating oligodendrocytes (Bernhardt et al., 2022; Dugas et al., 2012). Myelin and myelin-associated inhibitors (MAI) present another substantial barrier to axon growth and regeneration, offering another layer of robustness to the growth-inhibitory actions of the KLF repressors (Geoffroy & Zheng, 2014). Rising T3, through the KLFs, may coordinate the loss of axon growth through multiple independent mechanisms. This orthogonal role for the KLFs in oligodendrocytes emphasizes the necessity for cell type-specific manipulations when studying this TF family. The selectivity of our CRISPR-interference strategy for excitatory cortical neurons was therefore essential given the unavailability of conditional knockout lines for Klf6, Klf7, and Klf13.

By measuring gene expression changes following KD of KLF activator or repressor pairs, we identified hundreds of putative targets, many of which were bidirectionally regulated and subject to strong postnatal repression (**Supplementary Table 1**) and bidirectional regulation of two of these targets, *Tubb2b* and *Dpysl3*, was further validated *in situ* (**Figure 8**). In addition to being among those most potently affected by either manipulation, both genes are powerfully and globally downregulated during postnatal development and exhibit a developmental loss of chromatin accessibility at promoter elements containing putative KLF binding site(s) (**Figure 5D, S5B1**). TUBB2B is a ß-tubulin isoform expressed in the in the pre- and neonatal brain (Breuss et al., 2015). Mutations in the human TUBB2B gene broadly result in polymicrogyria with additional deficits in cognitive and motor function that vary with the site of mutation (Cederquist et al., 2012; Jaglin et al., 2009). In particular, mutations producing dominant-negative forms of TUBB2B have been shown to interfere with axon guidance, fasciculation, and arborization (Cederquist et al., 2012; Romaniello et al., 2012). *Klf6/7* KD targets also included multiple α- and β-Tubulins beyond *Tubb2b*, including *Tuba1a*, *Tubb3*, and *Tubb5,* all of which are weakly yet significantly upregulated by *Klf9/13* KD. Disruptions to any of these genes results in distinct yet overlapping clinical tubulinopathies characterized by cortical malformations frequently arising from defective neurogenesis or neuronal migration (Bahi-Buisson et al., 2014; Keays et al., 2007) that bear notable resemblance to the phenotype described in the Emx1-Cre;*Klf7*^fl/fl^ cortex (Hong et al., 2023). Furthermore, ß-tubulins were among the first growth cone-enriched proteins identified in regenerating rat sensory neurons, but the high sequence homology between ß-tubulin isoforms precludes immunohistochemical staining of specific isoforms (Hoffman, 1989; Hoffman & Cleveland, 1988) and leaves open the question of why loss of TUBB2B is not adequately compensated by other isoforms. Dpysl3 (also known as Crmp4), along with Dpysl5 (Crmp5; **Fig 7C**) are members of the Collapsin Response Mediator Protein family that collectively transduce extracellular signals to regulate growth cone guidance, retraction, and collapse in the central and peripheral nervous system (Schmidt & Strittmatter, 2007; Wang & Strittmatter, 1996). DPYSL3 mutations have been identified in human ASD patients, resulting in its’ designation as a SFARI Category 2 ASD risk gene (Tsutiya et al., 2017b). Mechanistically, Dpysl3 has been shown to promote microtubule assembly and actin bundling *in vitro* and developing Dpysl3-null mice show decreased axon extension and reduced growth cone size (Cha et al., 2016; Khazaei et al., 2014; Rosslenbroich et al., 2005).

Knockdown of *Klf6/7* also resulted in significant downregulation of genes with established roles in axon growth and maintenance previously identified as dysregulated in the olfactory epithelium of Klf7^-/-^ mice including *Gap43*, *Cdkn1a* (P21), *Stmn2* (SCG10), and *L1Cam* (Kajimura et al., 2007; **Figure 7E, F**). L1CAM, GAP-43, and STMN2 are neonatally lethal due to severe defects in the formation of major axon tracts (Yuanjun Li et al., 2023; Shen et al., 2002; Strittmatter et al., 1995) or are associated with a severe neurodevelopmental disorders (Camp et al., 1993; Fransen et al., 1998). GAP-43 is also necessary and sufficient to enhance regeneration following spinal cord injury (Bomze et al., 2001; Doster et al., 1991; Mason et al., 2002). Additional putative KLF targets of with known roles in development and disease are listed in Supplementary Table 1.

Targets of the KLF family are downregulated by coordinated withdrawal of activating drive and increasing repressive drive. This redundancy appears to have been maintained throughout mammalian evolution and may ensure its robustness. Given that regeneration of CNS axons would seem to be advantageous, why should repressing this capacity be so robustly conserved? A likely explanation is that axon growth and circuit refinement may be at odds with one another, and thus retaining developmental mechanisms that allow axons to freely grow could potentially disrupt the coherence of mature circuits. Related ideas have been suggested in the context of axon regeneration (Delpech et al., 2024; Kiyoshi & Tedeschi, 2020; Wang et al., 2015). Following the targeting of axons to their appropriate sites through chemical cues, precision in postnatal circuit development is achieved by activity-based synaptic pruning and retraction of exuberant axon branches and terminals (Kantor & Kolodkin, 2003; Luo & O’Leary, 2005). Disruptions to either of these processes can lead to altered circuit structure, but compromised refinement resulting from failures of synaptic pruning is believed to underlie human neuropsychiatric disorders such as ASD and schizophrenia (Riccomagno & Kolodkin, 2015). In early neonatal development when KLF activating drive is dominant, axons in the CNS exhibit stereotyped patterns of excessive outgrowth termed ‘axon exuberance’ (Innocenti & Price, 2005). This frequently takes the form of supernumerary axon terminals invading otherwise appropriate target regions, but some cortical neurons transiently extend entire axon collaterals to contralateral or subcortical regions that they later retract. Notable examples include the retraction of callosal projections from Layer 2/3 neurons in primary visual (V1) and somatosensory (S1) cortices with cell bodies distal to the V1/V2 (or S1/S2) border (Innocenti, 1981; Ivy & Killackey, 1982) and elimination of axon collaterals from Layer 5 neurons in V1 that transiently extend into the cervical spinal cord (Stanfield et al., 1982). Most of these stereotyped retraction and pruning events occur in the second to third postnatal week when KLF repressive drive becomes dominant and are complete by P30. This teases the possibility that the KLF switch may be important for mediating a transition from axon growth to circuit refinement in early adolescence through repression of pro-growth genes. In support of this, hypothyroid rats with presumably dysregulated Klf9 expression fail to trim the V1 to Spinal cord collateral, indicating that axon retraction is impaired in these animals (Li et al., 1995), and show impaired refinement of the corpus callosum (Gravel et al., 1990). Furthermore, 3 bidirectional targets of the KLF switch - *Dplys3*, *Dpysl5*, and *Plxna3* (Plexin-A3) – are transducers of the response to SEMA3A, one the best studied effectors of synapse retraction (Bagri et al., 2003; Cheng et al., 2001; Khdour et al., 2022; Low et al., 2008). Thus, manipulations of the KLF family that tip the balance in favor of its’ activators to promote axon regeneration may come at an unforeseen cost of synaptic specificity.

This study represents the first reported application of CRISPR interference for identifying TF targets in eukaryotic organisms. Our data demonstrates that this method is particularly well-suited for overcoming redundancy inherent in TF families to facilitate the prediction of shared and specific targets of TF paralogues. Furthermore, cre-dependent CRISPRi offers a flexible workaround to the lack of conditional knockout lines targeting many essential and redundant genes and the challenge of maintaining and breeding transgenic strains for every gene and cell type of interest. By pairing a modular cloning strategy for creating AAVs capable of multiplexed sgRNA delivery with neonatal ICV injections, we developed a streamlined gene knockdown approach that boasts remarkably high penetrance and apparent specificity. This method reliably yielded >95% knockdown of our genes of interest as early as 10 days post-injection, representing an improvement over many reports of knockdown efficiency *in vivo* or *in vitro*. This could be a product of the abundance of endogenous dCas9-KRAB in LSL-dCas9-KRAB mice and the high viral penetrance produced by ICV injections, which overcomes any limitations on CRISPRi imposed by insufficient enzyme or sgRNA concentration (Fontana et al., 2018; X. Li et al., 2016). By using CRIPSRi to knock down KLF repressor and activator pairs, we identified specific genes through which the KLF family could mediate its bidirectional effect on developmental gene expression and axon growth (Moore et al., 2009). While this provides insight into the genetic mechanisms underlying the loss of intrinsic axon growth capacity in maturing neurons, it is possible that postnatal repression of this program by KLF repressors is essential for circuit stability and precision in the mature brain.

## Supporting information

Shared and Disease-Associated putative targets of KLF Activators and Repressors

## Acknowledgements

We thank the Brandeis Light Microscopy Core Facility for assistance with image acquisition. This work was supported by R01NS109916 (SBN) and a grant from the W.M. Keck Foundation (S. Birren and SBN).

The authors declare no competing financial interests

## Methods

### Animal Husbandry and Transgenic Lines

All animals were bred, housed, and cared for in Foster Biomedical Research Laboratory at Brandeis University (Waltham, MA, USA). Animals were provided with food and water *ad libitum* and kept on a 12 hr:12 hr light:dark cycle. Cages were enriched with huts and tubes. All experiments were approved by the Institutional Animal Care and Use Committee of Brandeis University, Waltham, MA, USA (Protocol 22015A). Both sexes were used for all experiments.

*Rorb^GFP^* (*Rorb^1g^*) mice were obtained from Dr. Douglas Forrest (Liu et al., 2013) and maintained as heterozygotes, which labels Layer 4 RORβ+ neurons with GFP without disrupting layer 4 neuron identity or function (Clark et al., 2020).

56L mice, which label cortico-thalamic layer 6 neurons with mCitrine, were created through lentiviral enhancer trapping and characterized in Shima et al., 2016.

Emx1-IRES-Cre (Emx1-Cre) mice (Gorski et al., 2002) were obtained from Jackson labs (JAX #005628).

Rosa26:LSL-dCas9^KRAB^ (CRISPRi) mice (Gemberling et al., 2021) were obtained from Jackson labs (JAX #033066). To produce cell type specific dCas9-KRAB expression in the Emx1+ lineage, homozygous CRISPRi mice were bred with homozygous Emx1-Cre mice and F1 offspring (dCas9-Krab^+/-^;Emx1-Cre^+/-^) of both sexes were used for all experiments.

### Developmental RNA-seq Library Preparation

For RNA-seq of Layer 4 neurons and P2 Layer 6 neurons, libraries were prepared using Ovation Trio RNA-Seq library preparation kit with mouse rRNA depletion (0507–32) according to manufacturer’s specifications and sequenced on the Illumina NextSeq 550/500 platform as published in Clark et al., 2020. For P30 Layer 6 libraries, RNA was reverse transcribed using the NuGEN Ovation RNA-Seq System V2 (#7102; NuGEN) followed by fragmentation to an average of ∼200 bp and ligation of Illumina sequencing adaptors with the Encore Rapid Kit (0314; NuGEN). These libraries were sequenced on the Illumina HiSeq 2500 platform as published in Sugino et al., 2019.

### ATAC-seq Library Preparation

ATAC-seq libraries were prepared as described in Clark & Nelson, 2019. Briefly, tissue from 2 male and 2 female mice of the same age and strain were sorted together to collect 30,000-50,000 cells per library. Nuclei were tagmented for 30 minutes and libraries were prepared in accordance with (Corces et al., 2016) followed by paried end (PE) sequencing on the NextSeq Illumina platform.

### Developmental RNA-Seq Analysis

The adaptor sequence AGATCGGAAGAGCACACGTCTGAACTCCAGTCAC was trimmed from libraries using *cutadapt* v. 3.4 and libraries were mapped to the UCSC mm10 genome using STAR v. 2.7.10 (Dobin et al., 2013) using the ENCODE Long Read RNA-seq parameters. Reads mapping to exons of known genes were quantified using featureCounts in the *Rsubread* package (Liao et al., 2019).

Differential expression analysis was conducted using DESeq2 (Love et al., 2014) with model design ∼ Group, where group is the Age:Layer identity of each sample. Genes with significant changes in expression between P2 and P30 were identified independently for either cortical layer using cutoffs |log_2_FC| > 1, adjusted p value < 0.01, and TPM > 10 in at least one sample, and combined as a master list of 5,579 unique developmental DEGs for downstream analysis. Adjusted p values for each gene were reported from a Wald test with Benjamini-Hochberg (BH) correction for multiple comparisons, and log_2_FC values were shrunk using the *ashr* method. To identify shared and cell-type specific gene sets and plot heatmaps, a scaled counts matrix was generated by z-scoring the condition-averaged relative log (*rlog*) expression. The Gap Statistic method from R package *cluster* was used to identify k=6 as the optimal number of clusters, and genes were clustered using k-means clustering with 100 starts. Assignment of clusters as ‘Shared’ or ‘Layer *x* Specific’ was manually determined by inspecting plots of normalized gene expression within clusters as in **Figure 1** and **Figure S1**.

For comparison of Layer 4 and Layer 6 RNA-seq with other cortical cell types from Sugino et al., 2019, batch effects corresponding to sequencing location (Brandeis v. Janelia) were removed using limma’s *removeBatchEffect*() function prior to log_2_CPM conversion in edgeR.

Bigwig files for IGV visualization were generated from BAM files of 2 merged replicates using deepTools v. 3.5.4 *bamCoverage* normalized to counts per million mapped reads (CPM) while excluding reads mapped to the mitochondrial genome.

### ATAC-Seq Data Analysis

ATAC-seq libraries were processed using an abbreviated version of the ENCODE ATAC-seq pipeline. First, Illumina adaptors were trimmed using *cutadapt* v 3.4 and trimmed reads were mapped to the mm10 genome using bowtie2 v. 2.4.2 (<u>Langmead & Salzberg, 2012)</u>. Reads mapping to the mitochondrial genome, unannotated chromosomes, or ENCODE mm10 blacklist regions, or reads where MAPQ < 30, were removed using *samtools* v. 1.11 (Danecek et al., 2021), followed by removal of PCR duplicates with Picard tools v. 2.23.5 (Broad Institute). Reads were shifted and converted to PE BED files in *samtools* and MACS2 v. 2.2.7 was used to call peaks in each library (Y. Zhang et al., 2008). Only peaks identified in both replicates were retained in the final peak set. Peaks called in all 4 groups were merged into a non-redundant set of 242,127 peaks using the method described in <u>Corces et al., 2018</u>.

DiffBind was used for differential accessibility analysis. First, reads overlapping the non-redundant peak set were counted and filtered to remove peaks containing less than 10 counts in at least 1 library and peaks overlapping mm10 v2 blacklist regions (Amemiya et al., 2019) resulting in 99,995 peaks eligible for differential accessibility. Resulting peaks were then normalized using the TMM method (Reske et al., 2020; Robinson et al., 2010). Peaks with differential accessibility between P2 and P30 were identified independently for either cortical layer using cutoffs |log_2_FC| > 1 and FDR < 0.01, and combined as a master list of 38,321 unique developmental DARs for downstream analysis.

Peaks were assigned to genes and regulatory elements using the ChiPSeeker package in R using mm10 Ensembl annotations with the TSS region defined as -1000bp upstream of the TSS to 200bp downstream. Gene assignments for intronic/exonic peaks were adjusted to reflect the gene that they were within (as opposed to their nearest TSS). For subsequent analysis, the full list of all DARs was subdivided into Promoter Peaks, Intronic Peaks, and Intragenic Peaks according to their ChIPSeeker annotation.

To identify shared and cell-type specific peaks and plot heatmaps, a scaled counts matrix was generated by z-scoring the condition averages of Reads in Peaks-normalized counts. Optimal k was determined using the R package *cluster* as described above for RNA-seq analysis, resulting in k=2 Promoter Peak sets, k=5 Intronic Peak sets, and k=6 Intragenic Peak sets. Assignment of clusters as ‘Shared’ or ‘Layer *x* Specific’ was manually determined.

ATAC-seq coverage plots were created from RiP-normalized peaks using the *dba.plotProfile* function in DiffBind. Summit-centered regions of interest were taken by extending peaks in DAR clusters or peaks assigned to DEG sets out 1.5kb in either direction. For TSS-centered regions, the Ensembl TSS with the highest cumulative coverage across libraries was selected as the ‘primary’ TSS and extended 1.5kb in either direction. Normalized coverage at each position were averaged across genes within a sample, then across age groups for plotting chromatin changes between P2 and P30.

Bigwig files for IGV visualization were generated from BAM files of merged replicates using deepTools v. 3.5.4 *bamCoverage* normalized to reads per genomic content (RPGC) while excluding reads mapped to the mitochondrial genome or ENCODE blacklist regions.

### Motif Overrepresentation Analysis

Motif matches for the 836 motifs in the non-redundant JASPAR 2022 Vertebrate CORE collection (Castro-Mondragon et al., 2022) were identified in the merged, non-redundant peak set using FIMO from the MEME suite v. 5.5.0 (Bailey et al., 2015) with an adjusted q value cutoff of 0.05 using the nucleotide frequency in the full peak set as background. Motif matches passing threshold were counted in the original peak set using custom R script and used for Overrepresentation analysis. Peaks were identified as belonging to a test set based on cluster membership or ChIPSeeker gene assignment for DAR and DEG analysis respectively, and all remaining peaks were identified as the control set. Motif counts were binarized and a 2x2 contingency table was constructed for each motif. The effect size and significance of motif overrepresentation was calculated using Fisher’s Exact Tests followed by BH adjustment for multiple testing. As above, all Overrepresentation analysis was conducted separately for peaks assigned to different regulatory features (Promoter, Intron, or Intragenic). Motif matching and enrichment analysis using the 137 root motifs from the JASPAR 2022 Vertebrate CORE clusters (Castro-Mondragon et al., 2017) was conducted using identical methods.

### Gene Ontology Analysis

Gene ontology analysis was conducted using the *clusterprofiler* package in R (Yu et al., 2012). Overrepresentation of terms of interest in DEG clusters or genes assigned to DAR clusters was measured relative to in all genes eligible for inclusion in the test group using the *enrichGO* function with an adjusted p value cutoff of 0.05. For bar plots, redundant GO terms were simplified using a cutoff of 0.5.

### Gene Set Enrichment Analysis

To identify terms enriched among genes bidirectionally regulated by the KLF family by GSEA, Principal Component Analysis was performed on the *rlog* counts matrix of all KLF KD, which separated samples by Age (PC1) and ‘KLF Effect’ (PC2, as in **Figure 7A**). The absolute loadings from the second Principle Component (PC2) representing the magnitude of the ‘KLF Effect’ were used as input to *clusterprofiler*’s *gseGO* function with 10,000 permutations. For plotting, redundant GO terms were simplified using a cutoff of 0.5.

### sgRNA Design and Cloning

Single-guide RNA’s (sgRNA) were cloned into custom AAV transfer vectors using a custom cloning strategy. Briefly, sgRNA’s were selected from among the top 5 scoring sgRNA’s for the selected target in the mouse CRISPRi v2 library <u>(Maechler et al., n.d.)</u>. Guides were aligned to the target’s TSS region and inspected for overlap with other guides and to peaks identified in the developmental ATAC-seq dataset. Preference was given to sgRNA’s with >=50bp of separation from another sgRNA’s and located within an identified region of open chromatin. In the case where other guides overlapped the top scoring guide, an additional sgRNA was predicted using the CRISPick tool (Sanson et al., 2018).

Guides were ordered as oligonucleotides from Azenta (Genewiz) with additional bases appended to facilitate seamless Golden Gate Cloning into the Lenti-CRISPR-v2 vector (Addgene #52961) in frame with a U6 promoter sequence and sgRNA scaffold. Following annealing and phosphorylation of gRNA pairs as in Sanjana et al., 2014, guides were cloned into the Lenti-CRISPR-v2 vector by Golden Gate Cloning using BsmBI (NEB; R0739). Competent DH5a cells (Zymo; T3007), were transformed with the reaction product, and plasmid DNA was isolated from resultant colonies and sequenced to verify successful cloning. sgRNA cloned in frame with the U6 promoter and sgRNA scaffold were amplified from the vectors by PCR using primers in **Table 1** containing a 5’ recognition element for the Type IIS Restriction endonuclease SapI and unique overhang sequences to facilitate cloning of tandem arrays. PCR products were purified and cloned in tandem by Golden Gate Cloning using SapI (NEB; R0549) into a custom vector modified from the pAAV-hSyn-DIO-EGFP (Addgene #50457) or pAAV-hSyn-DIO-mCherry (Addgene #50459) AAV transfer plasmids. Custom sgRNA delivery AAV vectors were generated by inserting a dual-SapI cassette between the MluI and ApaI sites upstream of the hSyn promoter in either plasmid. Plasmids isolated from competent DH5a cells transformed with the reaction product were sequenced to confirm successful cloning of 2-4 guides.

**Table 1:**
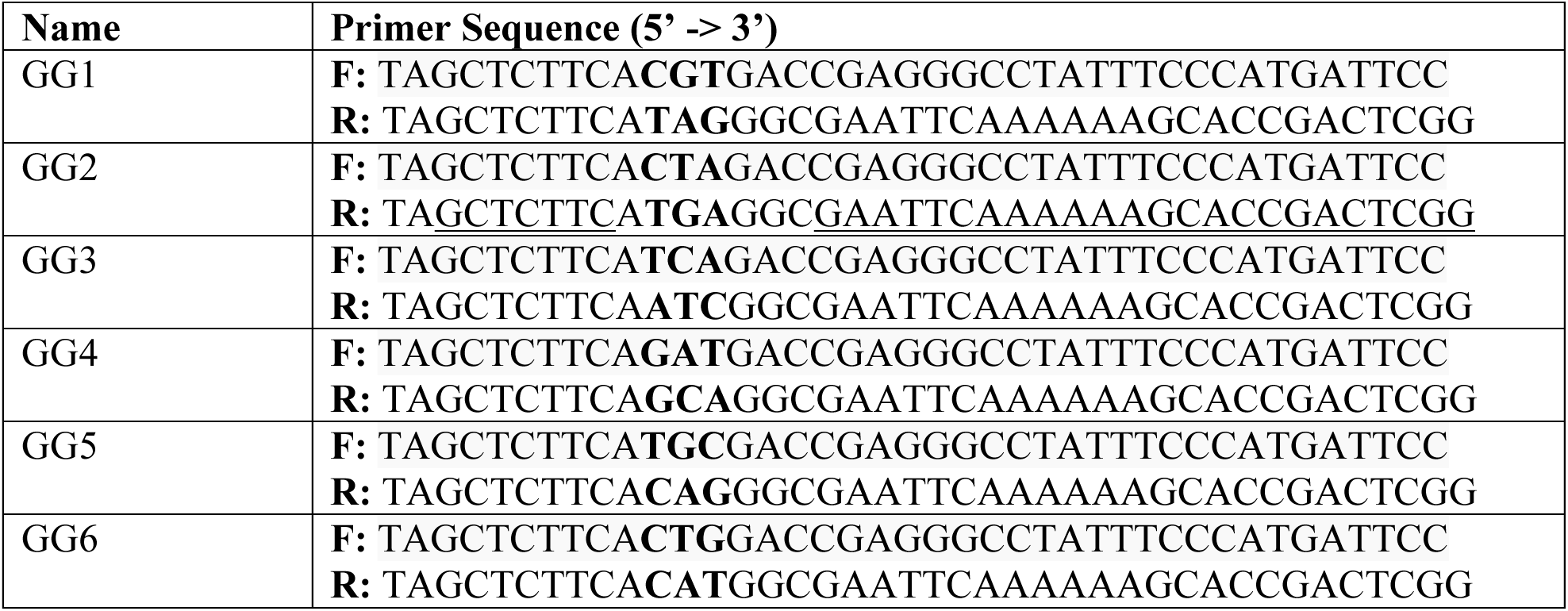
sgRNA Golden Gate Primers.

**Table 2:**
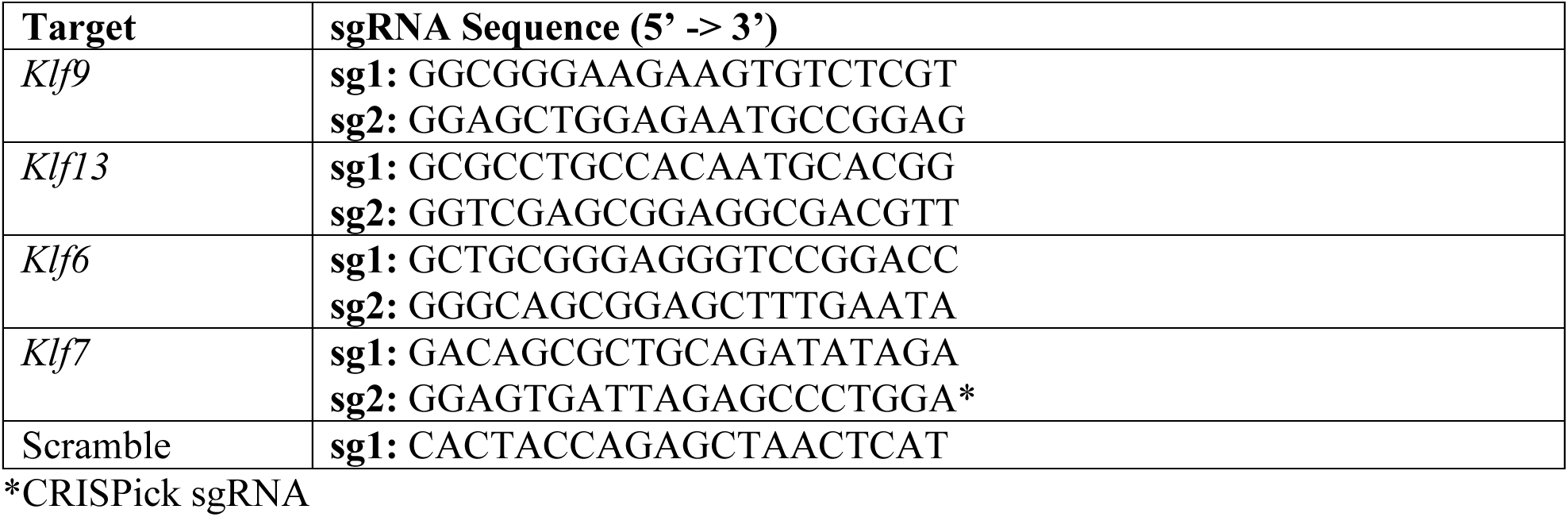
sgRNA Sequences.

**Table 3:**
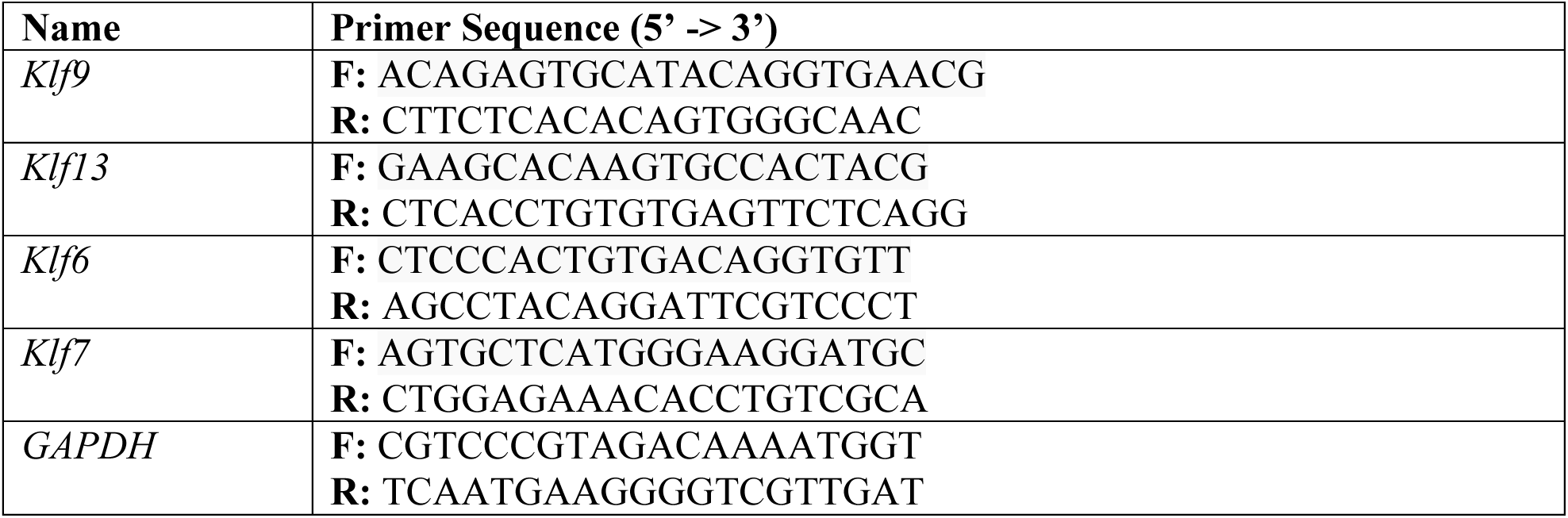
qPCR Primers.

### HEK293T Cell Culture AAV Production

HEK293T cells (ATCC; CRL-3216) were maintained at low passage numbers under standard cell culture conditions (37°C, 5% CO_2_) in DMEM, high glucose, pyruvate, (Gibco; 11-995) supplemented with 10% FBS (Gibco; 10438026) and 1% Penicillin/Streptomycin (Gibco; 15140122). For viral production, HEK293T cells were plated in 3-6 large 150mm cell culture plates (Corning; CLS430599) per virus and transfected with 10µg pAdDeltaF6 (pHelper, Addgene #112867), 6µg pAAV2/9n (Rep2/Cap9, Addgene #112865), and 6µg transfer plasmid (Custom, described above) using PEI MAX (PolySciences Inc.; 24765). Transfected cultures were maintained in 5% FBS media for 72 hours until harvesting, and pooled pellets were resuspended in 1x PBS-MK and stored at -80°C. rAAV was purified from frozen pellets using the IDX gradient method described in (Zolotukhin et al., 1999). Following ultracentrifugation, the viral fraction was repeatedly passed through an Amicon Ultra 100kDa dialysis column (Millipore; UFC910008) by centrifugation to remove IDX carryover and concentrate the virus in Lactated Ringer’s Solution. Resultant viral titer was between 2.0-8.0 x 10^13^ vg/ml as measured by qPCR using an AAV Titer kit (ABM; G931).

### Intracerebroventricular Injections

dCas9-KRAB^+/+^ mice were housed in harems with Emx1-Cre^+/+^ mice and dams were checked twice weekly for pregnancy. Pregnant mice were house individually with extra bedding and paper huts and monitored daily for new litters. At P0.5, mice were injected intracerebroventricularly with AAV using the protocol established in Kim et al., 2013. Briefly, pups were separated from their mothers in groups of 4 and anesthetized on ice.

Once movement could not be detected, pups were placed on a cooling block and injected bilaterally with 1µl AAV9-[U6-gRNA]-hSyn-DIO-[EGFP/mCherry] at 2.0-5.0x10^13^ viral genome copies/ml into their lateral ventricles. Pups were then marked by tail or toe clips and immediately warmed by hand until movement was restored, then placed in soft bedding on a heat pad until the entire cohort had been injected. Pups were returned to their mother once all pups had been injected, and mice were housed overnight on a heating pad. Injected pups were monitored the next day to monitor viability and ensure that maternal care was being provided, and cross-fostered with experienced dCas9-KRAB^+/+^ mothers if needed.

### Fluorescence Activated Cell Sorting (FACS)

Cells were sorted from acute coronal slices as in Clark & Nelson, 2019 and Clark et al., 2020. Briefly, animals were fatally anesthetized with 0.1ml/10g of a ketamine (20 mg/mL), xylazine (2.5 mg/mL), and acepromazine (0.5 mg/mL) (KXA) mixture. After loss of toe pinch response, animals P10 and older were transcardially perfused with 10-20ml of cold Choline Chloride Solution (CS) containing blockers of neuronal activity. Brains were swiftly extracted and sliced on a vibratome (Leica; VT1000S) at 300µm in cold CS, then slices were transferred to filtered inserts to equilibrate at 37°C for 15 minutes.

Inserts containing slices were transferred to room temperature Artificial Cerebrospinal Fluid (ACSF) containing blockers of neuronal activity and Protease for 90 minutes to digest tissue. Relevant cortical layers (for Layer 4/6 experiments) or densely infected regions (for CRISPRi experiments) were microdissected using a 18G syringe tip (BD Biosciences) under a fluorescent dissection microscope (Zeiss; MZFLIII) and kept on ice in ACSF containing 1% FBS (Gibco; 10438026). Dissected tissue was triturated using progressively smaller flame-polished glass pipettes and filtered through a cell strainer (Falcon; 352235), then centrifuged in a 10% Percoll gradient (Sigma-Aldrich; P1644) to generate a pellet enriched in pyramidal cells. The Pellet was resuspended in fresh ACSF + 1% FBS and stored on ice prior to isolation by FACS (BD FACSAria Flow Cytometer). Neurons in the top 10-25% of fluorescent cells above the autofluorescence background were collected. 1,000-5,000 cells were sorted into PicoPure Extraction buffer, incubated at 42°C for 30 minutes, and stored at -80°C until RNA extraction.

### RNA Extraction

RNA was isolated from sorted cell extract using PicoPure RNA Isolation Kit (Arcturus; KIT0204) with on-column DNAse digestion step (QIAGEN; 79254), and quantified on Qubit fluorometer using the RNA High Sensitivity reagents (Thermo; Q32852).

### cDNA Synthesis and qPCR

Reverse transcription was performed with 10-20ng of RNA using the iScript Reverse Transcription Supermix (BioRad; 1708891) according to manufacturer’s instructions. cDNA was diluted to a 0.25ng/µl equivalent and used as a template for qPCR with SsoAdvanced Universal SYBR Supermix (BioRad; 1725272) according to manufacturer’s instructions on a Corbett RotorGene 6000 thermocycler. All reactions were performed in triplicate.

qPCR primers were designed to span 3’ exon-exon junctions using Primer3plus and assessed for homo- and heterodimer formation using IDT’s oligoanalyzer tool. Primers were tested for efficiency and specificity against serial dilutions of whole cortex cDNA and only primer pairs with 95-105% efficiency were retained.

qPCR data was analyzed using the Ct method. To account for technical differences across several qPCR runs, relative expression was calculated using Ct’s obtained from all samples across multiple runs. The control Ct for each measured gene was computed by average Ct’s from the control or baseline group (Scramble sgRNA at P10-12 for Klf6/7 data, Scramble sgRNA at P18-20 for Klf9/13) and subtracted from the Ct to obtain the Ct. As primer efficiency was ∼100% for all genes measured, relative expression was simply calculated as. For Klf9/13 qPCR, significance was measured using a Wilcoxon rank sum test. For Klf6/7 qPCR, significance for the effect of Age and Virus on each measured gene was calculated using a 2-way ANOVA using formula Relative expression ∼ Age*Virus, followed by Wilcoxon rank sum tests.

### Perfusion

Animals were fatally anesthetized with KXA mixture. After loss of toe pinch response, mice were transcardially perfused with 10ml of cold 1x PBS followed by 10ml of cold 4% PFA. Brains were carefully removed from the skull and incubated in 4% PFA for 24-48 hours on a rocker at 4°C before being transferred to 1x PBS for storage at 4°C.

For P2 brains, wild-type pups were anesthetized on ice until there was no toe pinch response and then swiftly decapitated. The skin of the head was carefully peeled back and the brain and skull were washed with ice cold 1x PBS every 30 seconds while the skull and pia were peeled away with fine forceps. Brains were immediately transferred to 5ml tubes containing ice cold 1x PBS and washed twice with cold PBS to remove residual blood, then resuspended in ice cold 4% PFA and incubated overnight at 4°C on a rocker.

### Vibratome Sectioning and Epifluorescence Imaging

The cerebellum of fixed brains was removed using a scalpel and brains were imbedded in 4% Agarose dissolved 1x PBS. Brains were mounted on a pedestal and sliced on a vibratome (Leica; VT1000S) in groups of 2-4 at a thickness of 50 µm. Slices were transferred to 24 well plates containing 1x PBS + 0.05% Sodium Azide using a paintbrush and stored protected from light at 4°C.

For imaging of representative slices, selected slices were incubated in 1µg/ml DAPI (Thermo; 62248) in 1x PBS for 10 minutes at Room temperature, washed once with PBS for 5 minutes, and mounted on slides with VectaMount Mounting Medium (SouthernBiotech; 0100-01). Slides were sealed with nail polish and stored at 4°C prior to imaging. Whole brain slices were imaged on a Keyence BZ-X epiflourescent microscope in a single z plane with exposure settings calibrated to prevent saturation of the brightest infected cells and stitched using Keyence’s native stitching algorithm. Consequently, infection appears biased towards thick-tufted Layer 5 neurons while the true laminar distribution is much broader (Kirk et al., 2025*, in preparation*).

### CRISPRi RNA-Sequencing

Prior to library preparation, RNA quality was measured High Sensitivity RNA ScreenTape (Agilent; 5067-5579). For experiments using AAV containing mCherry as a reporter, RNA-seq libraries were prepared using the SMARTer Stranded Total RNA-Seq Kit v3 (Takara Bio; 634485) with 5-10ng of RNA as input according to manufacturer’s instructions. Library fragment size was evaluated using D1000 High Sensitivity ScreenTape (Agilent; 5067-5584). For experiments using EGFP as a reporter, libraries were prepared using the SMART-Seq mRNA LP kit (Takara Bio; 634762) with 5-10ng of RNA as input followed by 2ng of cDNA according to manufacturer’s instructions. cDNA concentration and library fragment size distribution were measured using D5000 High Sensitivity ScreenTape (Agilent; 5067-5593). For all libraries, concentration was measured using KAPA qPCR Library quantification kit (Roche; KK4824). Samples were pooled for PE sequencing on the Illumina NovaSeq platform (Novogene) or Illumina NextSeq 550/500 platform.

### RNA-Sequencing Analysis

Illumina TruSeq adapter sequences were trimmed from all libraries using *cutadapt* v 3.4, and reads with MAPQ < 30 were removed. For v3 libraries containing UMIs, 8bp UMIs were trimmed from Read 2 and reads were marked using *umitools* v. 1.1.2, and reads were trimmed to 75bp for faster mapping using *cutadapt*. Reads were then mapped to the GENCODE M27 genome using STAR v. 2.7.8a with ENCODE Long Read parameters, and mapped reads were deduplicated using *umitools*. For LP libraries, *cutadapt* was used to trim bases added by the SMART-Seq TSO ATTGCGCAATGNNNNNNNNGGGG from Read 1 prior to mapping to the GENCODE M27 genome by STAR as above. Paired end reads mapping to exons of known genes were quantified using featureCounts in the *Rsubread* package.

Differential expression analysis was conducted using DESeq2 with model design ∼ Batch + Group, where Group is the Age:Virus identity of each sample controlling for batch effects across sequencing runs. DEGs in each KD condition were identified relative to Scramble gRNA controls of the same age with cutoffs |log_2_FC| > 1 and adjusted p value < 0.05. Adjusted p values for each gene were reported from a Wald test with Benjamini-Hochberg (BH) correction for multiple comparisons, and log_2_FC values were shrunk using the *ashr* method.

For plotting heatmaps and normalized line plots, scaled counts matrices were generated by z-scoring the condition-averaged relative log (*rlog*) expression and corrected for batch effects using the removeBatchEffect function from *limma* (Ritchie et al., 2015). Optimal cluster numbers for heatmaps were determined as above, but implemented in *ComplexHeatmap* (Gu et al., 2016) using 100 random starts. Motif counting and enrichment in Promoter peaks were conducted using the methods described above.

For coverage plots, control genes were chosen among those unaffected by either KLF manipulation with a quantile-matched distribution of log2-FoldChanges between P2 and P30 in order to account for expected effects of developmental expression on chromatin accessibility. For analysis of the effect of Thyroid Hormone on KLF target expression, snRNA-seq data from T3-treated and control mice published in Hochbaum et al., 2019 were downloaded from the CELL x GENE repository (https://cellxgene.cziscience.com/collections/c450e15d-321a-42d6-986b-11409d04896d), converted to a Seurat object in R, and filtered for excitatory neuronal cell type clusters annotated by the authors. Differentially expressed genes were identified using Seurat’s FindMarkers() function comparing T3 and Control neurons within each cluster using a Wilcox test with a log fold-change threshold of 0.1. The resulting data was subset by high-confidence shared KLF targets identified as a DEG in at least one cluster for plotting.

### Electrophysiology

Recordings were performed as described in Valakh et al., 2023. Mice were fatally anesthetized with KXA between P18-20 and their brains were swiftly removed and allowed to acclimate in ice cold oxygenated ACSF containing (in mM) 126 NaCl, 3 KCl, 1 NaH_2_PO_4_, 25 NaHCO_3_, 2 MgCl_2_, 2 CaCl_2_, and 10 Glucose. 300µm acute coronal slices were made using a vibrating microtome (Leica; VT1000S) and slice were incubated at 37°C in oxygenated ACSF for 30 minutes to 1 hour. Slices were subsequently maintained in oxygenated room temperature ACSF and recordings were conducted at 34-35°C. Neurons were visualized on an Olympus upright epifluorescence microscope 40× water immersion objectives and infected cells were identified based on presence of an mCherry reporter. Whole-cell patch-clamp recordings were made using near-infrared differential interference contrast microscopy. Recording pipettes of 3–5 MΩ resistance contained internal solution containing 20 KCl, 100 K-gluconate, 10 HEPES, 4 Mg-ATP, 0.3 Na-GTP, 10 Na-phosphocreatine, and 0.1% biocytin. Recordings were amplified (Multiclamp 700B, Molecular Devices) and digitized at 4 kHz using a National Instruments Board under control of IGOR Pro (WaveMetrics).

For Intrinsic Excitability recordings, synaptic currents were blocked with 25µM picrotoxin, 25µM DNQX, and 35µM APV. Cells were maintained at -70mV by injecting the required current and steady state series resistance was compensated. 500ms currents between 50–700 pA were randomly injected every 10s (at least 2 times per cell) with a 25 pA seal test before each trial.

To record miniature Excitatory AMPA currents, action potentials and non-AMPA synaptic currents were blocked with 25µM picrotoxin, 500nM TTX, and 35µM APV. Cells were clamped at -70mV with no compensation for the liquid junction potential and at least 3 10-second sweeps were recorded for each cell with a seal test before each trial.

Threshold-based detection of mPSCs and action potentials was implemented using custom-written R scripts with routines for baseline subtraction, custom filtering, and measurements of intervals, amplitudes, and kinetic properties. Comparisons between sgRNA treatment groups were made by a 2-way ANOVA for the effects of Current and sgRNA for intrinsic excitability, and a Mann-Whitney U test for mEPSC parameters.

For visualization of patched cells by biocytin staining, slices were fixed overnight in 4% PFA at 4°C followed by 2 1X PBS washes and 2 hours in Blocking Buffer (1x PBS supplemented with 5% Normal Goat Serum (NGS, Gibco), 3% Bovine Serum Albumin (BSA, Thermo), and 0.5% Triton-X-100 (Sigma)). Slices were then incubated at 4°C overnight with Alexa-594-Streptavidin diluted 1:500 in Blocking Buffer. The following day, slices were washed 3 times in 1x PBS, stained with 1µg/ml DAPI (Thermo; 62248) in 1x PBS for 10 minutes at Room temperature, washed once with PBS for 5 minutes, and mounted on slides with VectaMount Mounting Medium (SouthernBiotech; 0100-01). Slides were sealed with nail polish and stored at 4°C prior to imaging. Slides were imaged on a Nikon AX-R Confocal Microscope with a 20x air objective by taking Z-stacks spanning the entire dendritic arbor at a 5µm step size. Images were converted into maximum intensity projections in Fiji.

### RNAScope

Following overnight incubation in 4% PFA, brains were transferred to 30% Sucrose/PBS overnight followed by 15% Sucrose/PBS overnight to dehydrate tissue. The olfactory bulb and cerebellum were removed using razor blades in a mouse brain matrix (AgnTho’s; 69-1175-1) and brains were flash frozen in OCT (Fisher; 23-730-571) molds by submerging them in a 100% Ethanol and dry ice bath for 5 minutes. Frozen tissue blocks were immediately transferred to -80°C.

Before slicing, brains were equilibrated to -20°C for ∼30 minutes and excess OCT was removed. Brains were mounted onto a freezing microtome sliced at 16µm with 100µm between each slice with slices centered around primary somatosensory cortex (S1). Slices were transferred to PBS using a paintbrush and mounted on charged slides (Fisher; 12-550-15). Slides were dried at 4°C overnight and transferred to a sealed slide box at -80°C the following day.

For RNAScope of brains infected with AAV, we began the RNAScope Multiplexed Fluorescent Assay v2 protocol (ACD; PN 323110) just before the protease treatment step to avoid denaturing viral EGFP during the Target Retrieval step. Slides with 1-2 slices each were removed from - 80°C, washed once with PBS, and allowed to air dry before application of a hydrophobic barrier. Slides were stored overnight at 4°C and probed following the standard RNAScope protocol beginning with Protease III treatment. P20 brains were probed with Mm-C2-Klf9 (ACD; 488371-C2) and *either* Mm-C1-Dpysl3 (ACD; 496611) or Mm-C1-Tubb2b (ACD; 1039621-C1) and stained with Opal 620 Dye (Akoya; FP1495001KT) at a 1:750 dilution for the C2 channel and Opal 690 (Akoya; FP1497001KT) at 1:750 for C1. P10 brains were probed with Mm-C2-Klf7 (ACD; 570961-C2) and *either* Mm-C1-Dpysl3 or Mm-C1-Tubb2b and stained with Opal 620 Dye at a 1:1500 dilution for the C1 channel and Opal 690 at 1:750 for C2.

For RNAScope of *Klf9* and *Klf7* expression across development, samples were processed following the RNAScope Multiplexed Fluorescent Assay v2 protocols with 2 exceptions: for P2 and P7 samples, the Target Retrieval step was reduced to 5 minutes and brains were treated with with Proteaste Plus for 15 minutes to prevent over-digestion of fragile neonatal tissue. All slices were probed with Mm-C1-Klf7 (ACD; 570961) and Mm-C2-Klf9 and stained with Opal 520 Dye (Akoya; FP1487001KT) at a 1:1500 dilution for the C1 channel and Opal 620 at 1:1500 for C2.

### RNAScope Image Acquisition and Analysis

All slides were imaged on a Zeiss 880 confocal microscope. For CRISPRi samples, z-stacks were acquired from at least 2 locations in each hemisphere centered on cortical Layer 5 (minimum 4 total) under a 40x oil-immersion objective. For P2 brains, stitch-stacks spanning the entire height of the cortex were acquired under a 40x oil-immersion objective and stitched using the native stitching algorithm in Zen (Zeiss). For P7 and P30 brains, stitch-stacks containing all cortical layers were acquired under a 20x air-immersion objective and stitched as above.

Image analysis and quantification was conducted using custom Python scripts. For RNAScope of *Klf9* and *Klf7* expression across development, images were first rotated so that the pial surface was parallel to the bottom and top borders of the image, then cropped so images spanned from the bottom of Layer 6 to the epithelial cell layer above Layer 2/3 (Dorsal-Ventral axis). The nuclear DAPI channel was downscaled as above and used to segment and label nuclei with CellPose v. 3.0.1 (Stringer, et al., 2021) in 2.5-D mode using a stitch threshold of 0.3, cell probability threshold of 3, and a diameter of 25 pixels for 40x images and 12 for 20x images.

Artifacts were removed by excluding labels with a total pixel volume in the bottom 2% of the measured range. RNA puncta were segmented and labeled as above. The number of unique nuclei and RNA labels in each x-by-z slice along the y (Dorsal-Ventral) axis were counted and exported to R for downstream analysis. To normalize RNA puncta counts for differences in cortical thickness across animals and ages, a Binned Dorsal-Ventral Axis was created by binning tables containing counts of nuclei and RNA along the y-axis into n=2000 bins for P7 and P30 samples and n=1500 bins for P2 samples. Finally, RNA puncta counts were normalized by nuclei counts within each bin to create a measure of RNA quantity scaled by cell density. 95% confidence intervals for both mRNA species were estimated by bootstrapping with 1000 random resamples using the R package *boot*. Significance of the difference between *Klf9* and *Klf7* abundance for each age group at each position was measured by a Mann-Whitney U test with BH correction.

For CRISPRi RNAScope, images were split into their composite channels and the EGFP channel was downscaled by 4x and a gaussian blur was applied with sigma=1.3. The downscaled image was used to segment and label cells with CellPose v. 3.0.1 (Stringer et al., 2020) in 2.5-D mode using the *cyto2* model with a stitch threshold of 0.5 and a diameter of 40 pixels. Presumptive false positives were removed by excluding labels with a summed fluorescence signal in the bottom 10% of the measured range or with more than 5% of their pixel volume along the edges of the image (including top and bottom). Remaining labels were upscaled 4x to the dimensions of the original EGFP image. The raw image intensities from the 2 RNA channels were reciprocally subtracted from one another to eliminate bleed-through artifacts, and RNA puncta were labeled and segmented using *pyclesperanto*’s *veronoi_otsu_labeling* with a spot sigma of 2 and outline sigma of 1. The overlap (in pixels) of every RNA puncta with every cell label was calculated and exported to R for downstream analysis. Puncta below 20 pixels in size or puncta overlapping cells by less than 75% of their pixel volume were excluded. The number of unique RNA puncta overlapping each cell label was quantified and scaled by the area of the cell label in pixels×10^-6^ to produce area scaled puncta counts. Significance for the difference between each gene across treatment groups was measured by an independent samples t-test.

---

**Figure S1.1.**
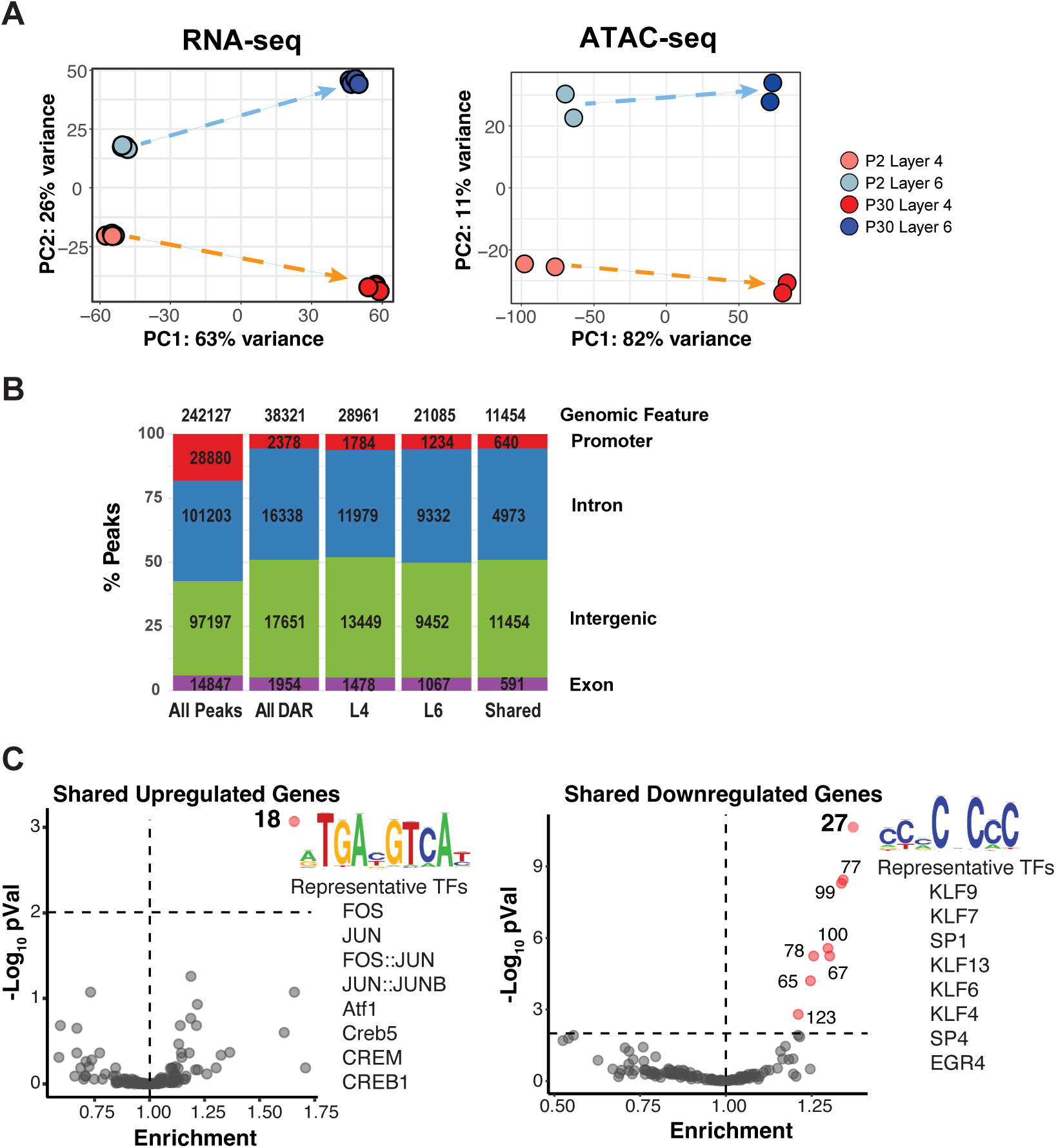
Shared gene expression and chromatin accessibility changes in maturing layer 4 and layer 6 pyramidal neurons. **A)** Principal Component Analysis of RNA-Seq (A1, n=4 per condition) and ATAC-seq (A2, n=2 per condition) samples used in this analysis **B)** Distributions of ATAC-seq Peak Annotations for All Peaks, All DARs, Layer 4 DARs, Layer 6 DARs, and DARs identified in both cell types. **C)** Transcription Factor Motif Enrichment in ATAC-seq peaks overlapping the Promoter (TSS -1000/+200 bp) of shared up- (**left**) or down-regulated (**right**) genes relative to all Promoter peaks using the JASPAR 2022 Cluster Root Motifs (Fisher Exact tests with BH adjustment). Cluster 18 includes AP-1 family Transcription Factors. Cluster 27 contains all KLF/Sp Transcription Factors. Consensus motif and representative members are indicated on the right for each gene set.

**Figure S1.2.**
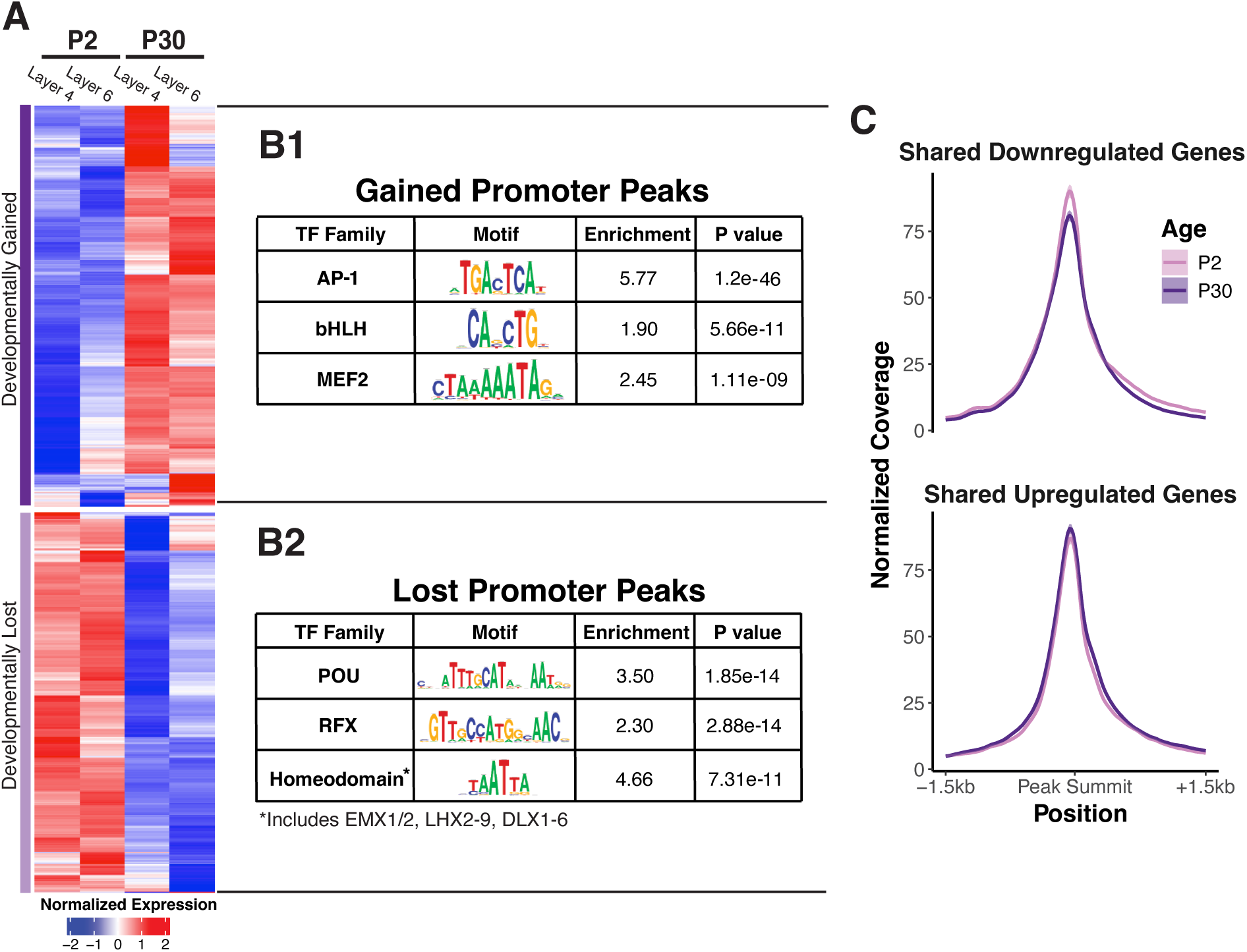
Shared chromatin accessibility changes in maturing layer 4 and layer 6 pyramidal neurons. **A)** Heatmap of all DARs in gene Promoters identified in either layer between P2 and P30, separated into k=2 clusters. (DARs defined as |log2FoldChange|>1; p.adj<0.01 by Wald’s Test with BH correction; min(TMM)>10). Data plotted as group averages of z-scored log2(TMM). **B)** Transcription Factor Motif Enrichment in Gained **(B1)** and Lost **(B2)** ATAC-seq Promoter peaks (TSS -1000/+200 bp) relative to all Promoter peaks using the JASPAR 2022 Cluster Root motif set (Fisher Exact tests with BH adjustment). **C**) RiP-normalized coverage of average ATAC-seq signal at Promoter peaks +/-1.5kb of shared downregulated (top) or shared upregulated (bottom) DEGs in all P2 or P30 samples (Shaded region = 95% CI; n=4 libraries per group, aggregate of 2 cell types).

**Figure S2.**
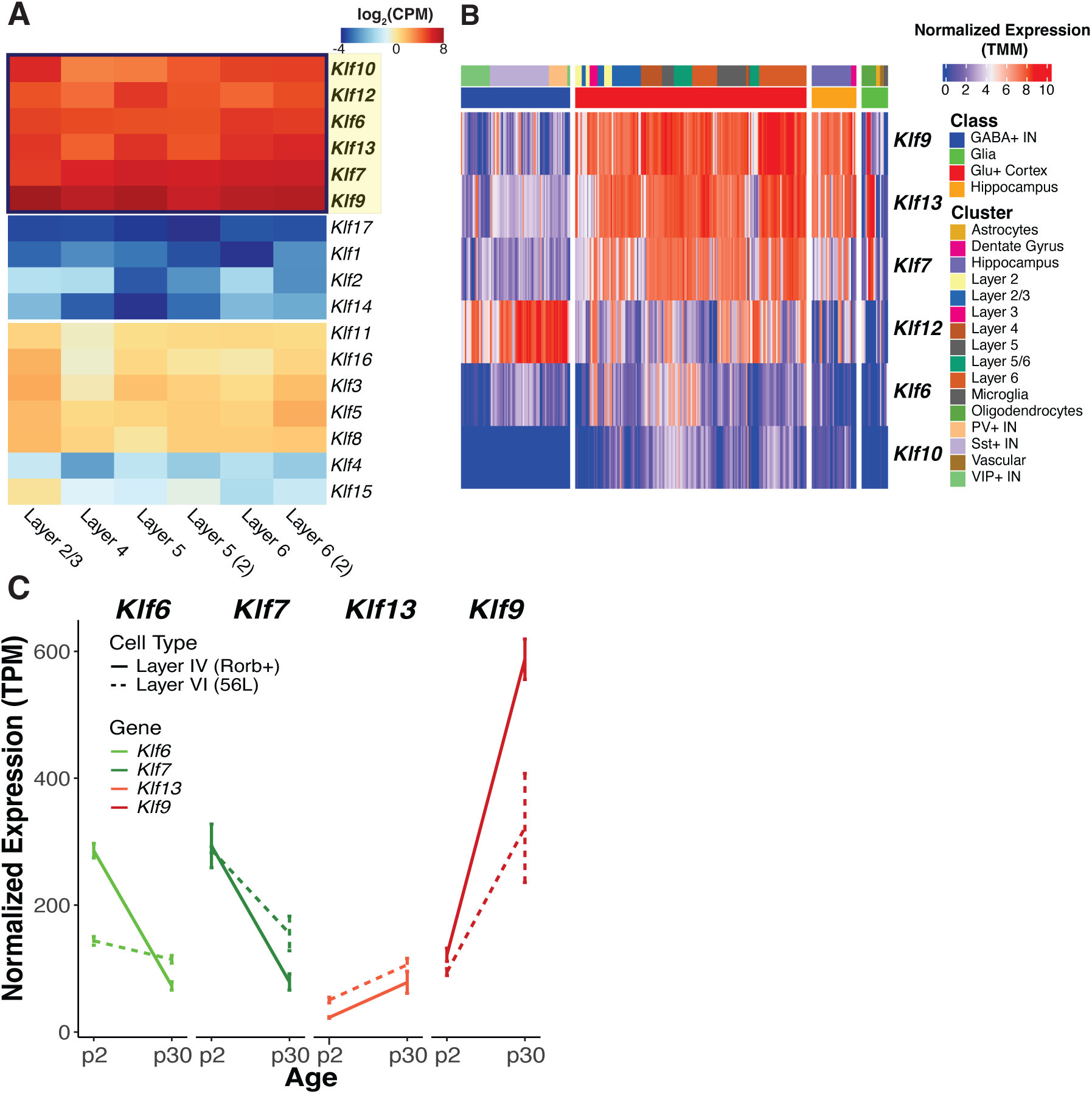
Activating (*Klf6* and *Klf7*) and repressive (*Klf9* and *Klf13*) members of the KLF family display opposing patterns of gene expression throughout the developing cortex. **A**) Expression of all KLF family members in major excitatory neuronal cell types of the cortex obtained by bulk RNA-seq (**left**, Sugino et al., 2019) and in the principal cell clusters of the mouse cortex and hippocampus obtained by scRNA-seq (**right**, Yao et al., 2021) **B**) Expression patterns of developmentally regulated KLFs between P2 and P30 in Layer 4 and Layer 6 neurons (RNA-seq, n=4 mice per group).

**Figure S4.1.**
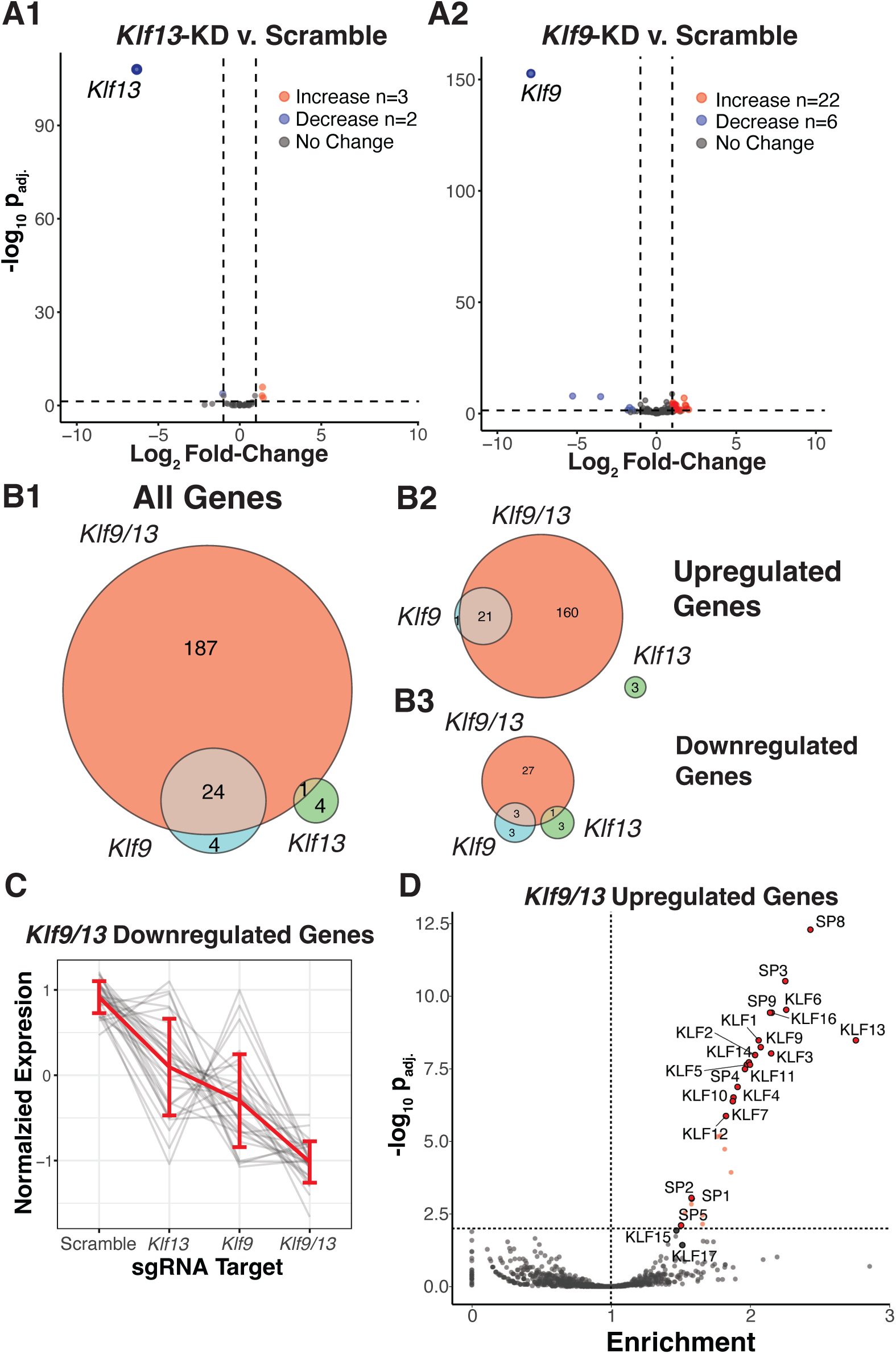
*Klf9* and *Klf13* show compensatory activity and additive regulation of target genes. **A)** Volcano plots of gene expression changes in *Klf9* KD (**A1**) or *Klf13* KD (**A2**) relative to Scramble Controls (DEGs defined as |log2FoldChange|>1; p.adj<0.05 by Wald’s Test with BH correction) **B)** Venn Diagrams comparing All DEGs (**B1**), Upregulated DEGs (**B2**), and Downregulated DEGs (**B3**) identified in *Klf9*, *Klf13*, and *Klf9/13* KD samples relative to scramble controls **C)** Line plot of all downregulated Klf9/13 targets across all libraries. Grey lines are z-scored, regularized log-transformed RNA-seq counts of individual genes, red lines represent average of all genes +/- SEM. (***:p<0.0001, Kruskal-Wallis test with post-hoc pairwise Mann-Whitney U test) **D)** Transcription Factor Motif Enrichment in ATAC-seq peaks overlapping the Promoter (TSS -1000/+200 bp) of Klf9/13 targets (upregulated DEGs) relative to Promoter peaks in non-DEGs using the JASPAR 2022 CORE Motif set (Fisher Exact tests with BH adjustment).

**Figure S4.2.**
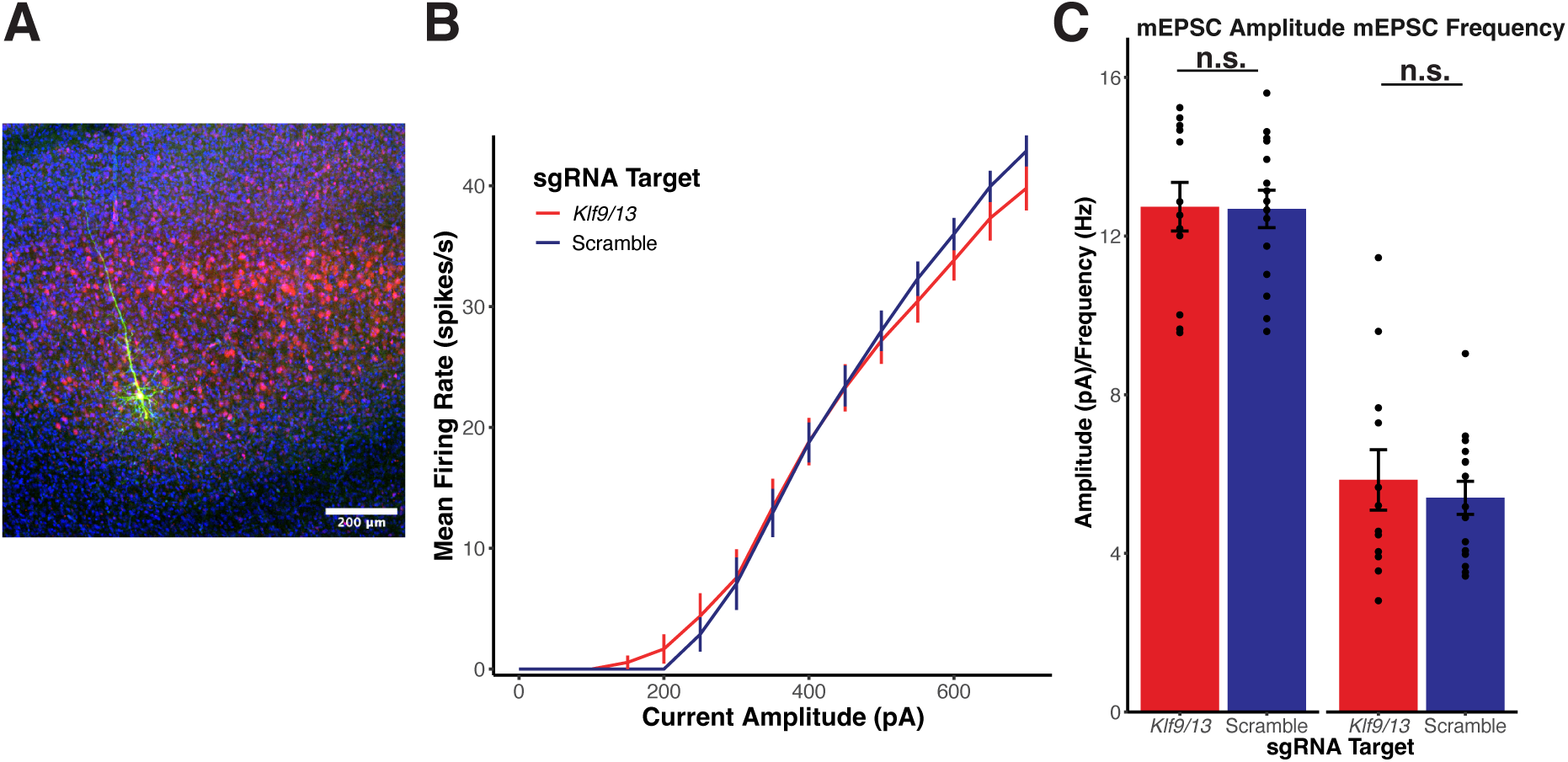
Normal intrinsic and synaptic properties of *Klf9/13* KD neurons. **A)** Example biocytin fill of Layer 5 pyramidal neuron (green) infected with Klf9/13-mCherry KD virus (red) at P0.5 (Scale bar = 200µm) **B)** Intrinsic excitability is unchanged between *Klf9/13* KD neurons (N=15) and Scrambled controls (N=10). (p = 0.787; Virus::Current effect from 2-Way ANOVA for Current and Virus) **C)** Miniature Excitatory Postsynaptic Current (mEPSC) properties of *Klf9/13* KD neurons (red, N=12) and Scrambled controls (blue, N=15) are not different (top) (Amplitude: Klf9/13-mCherry: 12.9±0.614 pA, Scr-mCherry: 12.7±0.47 pA, p=0.90, Mann-Whitney U test; Frequency: Klf9/13-mCherry: 5.85±0.76 Hz; Scr-mCherry: 5.4±42 Hz p= 0.83, Mann-Whitney U test)

**Figure S5.**
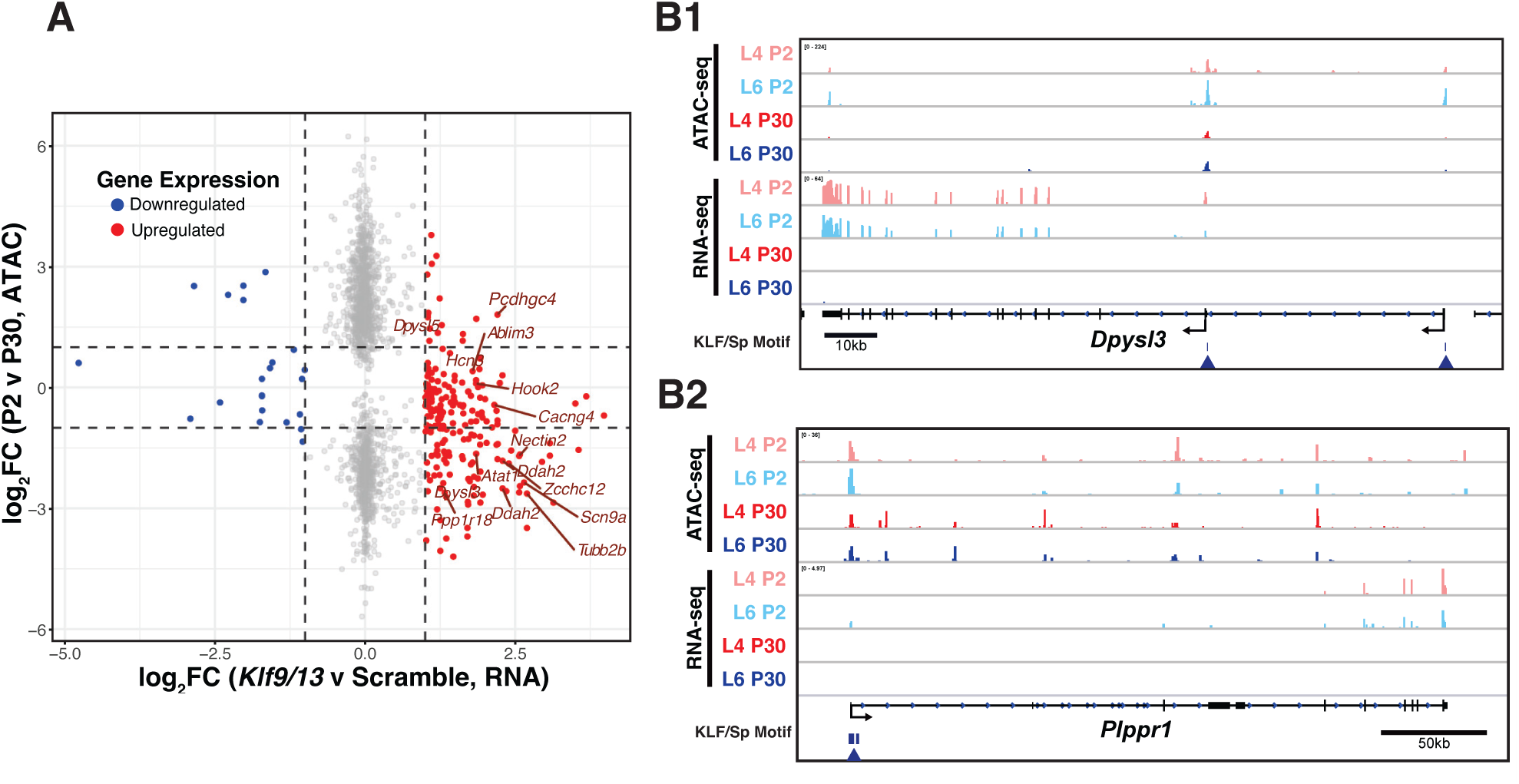
Putative Klf9/13 Targets are developmentally downregulated in the maturing cortex. **A)** Gene expression changes in *Klf9/13* KD relative to Scramble Controls at P18-20 (n=4 mice per group) relative to chromatin accessibility changes between P2 and P30 (n=4 mice per group, aggregate of 2 cell types). Only DEGs or genes with at least one Differentially Accessible Promoter peak are plotted. Points represent in individual genes, and colors indicate the direction of change in Klf9/13 KD samples. Genes with more than one Promoter peak may appear multiple times. **B)** Example IGV traces from P2 & P30 ATAC-seq and RNA-seq for Klf9/13 Targets *Plppr1* (**B1**, Developmental DEG *without* Promoter DAR) and *Dpysl3* (**B2**, Developmental DEG *with* Promoter DAR). Arrows: putative KLF/Sp binding site identified by FIMO (q<0.05)

**Figure S6.**
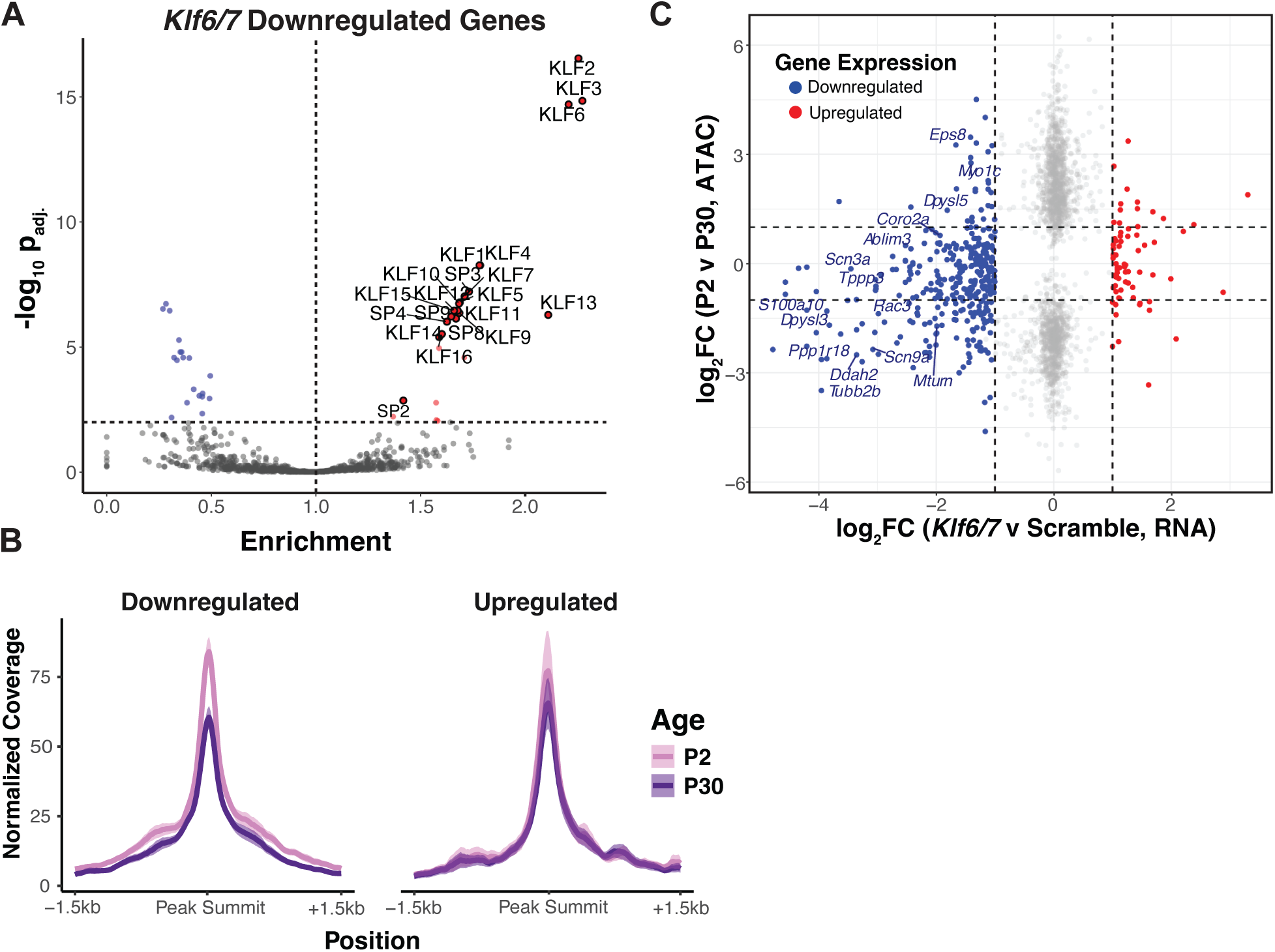
*Klf6* and *Klf7* promote expression of developmentally regulated genes in the perinatal cortex. **A)** Transcription Factor Motif Enrichment in ATAC-seq peaks overlapping the Promoter (TSS -1000/+200 bp) of Klf6/7 P10 targets (downregulated DEGs in Clusters 1 and 2) relative to all Promoter peaks in non-DEGs using the JASPAR 2022 CORE Motif set (Fisher Exact tests with BH adjustment). **B)** RiP-normalized coverage of average ATAC-seq signal at Promoter peaks around DEGs identified from *Klf6/7* P10 KD RNA-seq (Shaded region = 95% CI; n=4 libraries per group, aggregate of 2 cell types). **C)** Gene expression changes in *Klf6/7* KD relative to Scramble Controls at P10-12 (n=4 mice per group) relative to chromatin accessibility changes between P2 and P30 (n=4 mice per group, aggregate of 2 cell types). Only DEGs or genes with at least one Differentially Accessible Promoter peak are plotted. Points represent in individual genes, and colors indicate the direction of change in *Klf6/7* KD samples. Genes with more than one Promoter peak may appear multiple times.

**Figure S7.1.**
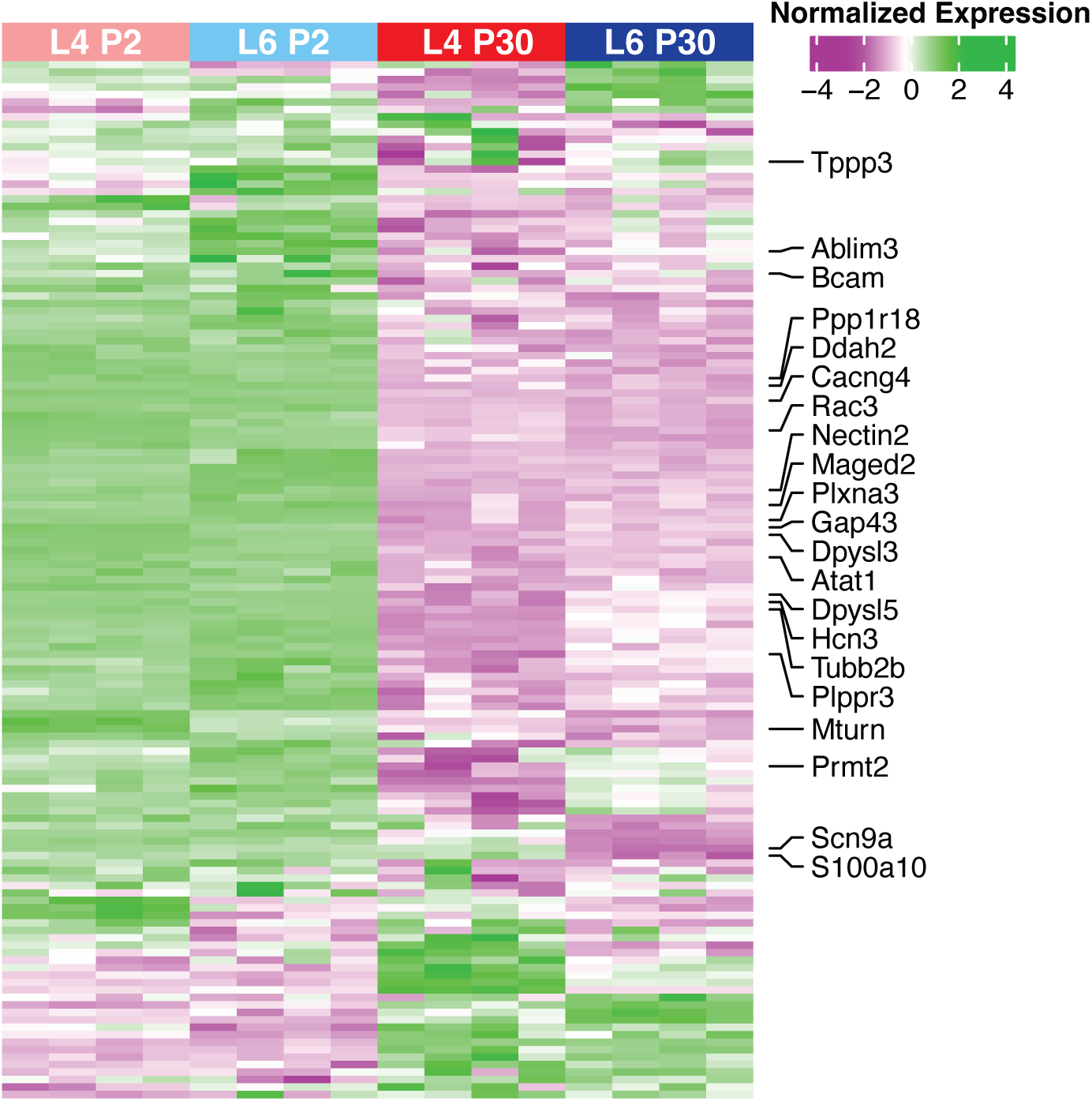
Shared KLF targets are similarly expressed and developmentally regulated in layer 4 and layer 6 neurons. **A)** Heatmap of scored, log-normalized gene expression of 144 Shared KLF Targets in Layer 4 and Layer 6 neurons at P2 and P30 (n=4 mice per group). Candidate genes with significant axonal or neuronal roles are highlighted on the right.

**Figure S7.2.**
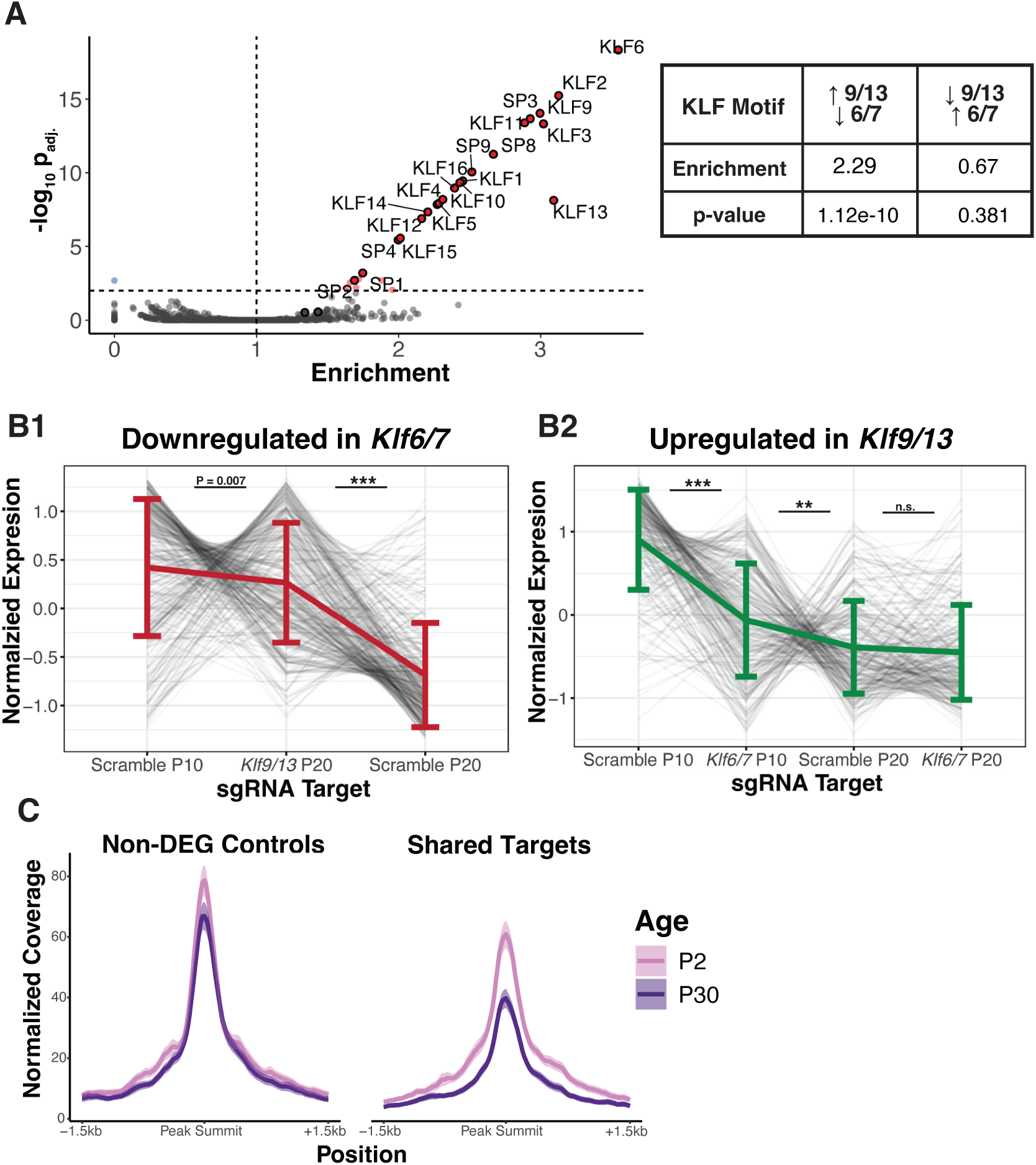
The transition from KLF Activators to Repressors constitutes a Developmental switch at shared targets. **A)** Shared Targets are highly enriched for the KLF/Sp Motif. Transcription Factor Motif Enrichment in ATAC-seq peaks overlapping the Promoter (TSS - 1000/+200 bp) of shared targets for the root KLF/Sp motif (top, Fisher Exact Test) and the full JASPAR 2022 CORE Motif set (bottom, Fisher Exact tests with BH adjustment). Enrichment is measured relative to promoter peaks in all non-DEGs. **B)** Klf6/7 Targets are upregulated by *Klf9/13* KD (**left**) and Klf9/13 Targets are downregulated by *Klf6/7* KD (**right**). Grey lines are z-scored, averaged regularized log-transformed RNA-seq counts of individual genes and dark line represent average of all genes +/- SEM (n=4 mice per group, Kruskal-Wallis test with post-hoc pairwise Mann-Whitney U test with BH adjustment, *:p<0.01; **p<1e-6 ***: p<2e-16). **C)** Loss of chromatin accessibility around at Shared Targets. RiP-normalized coverage of average ATAC-seq signal at Promoter peaks around Shared Targets and unaffected control genes (left; Shaded region = 95% CI; n=4 libraries per group, aggregate of 2 cell types).

**Figure S7.3.**
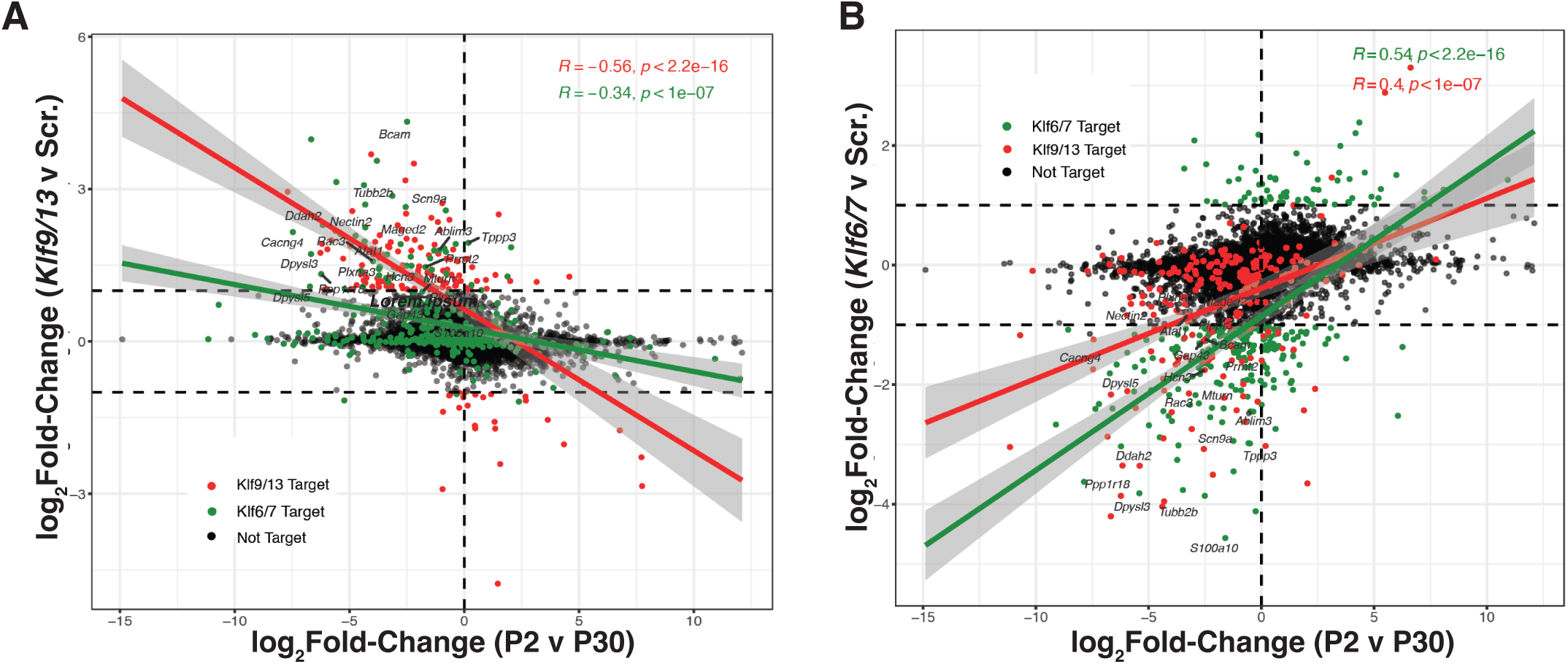
The transition from KLF Activators to Repressors constitutes a Developmental switch at shared targets. **A)** Gene expression changes in *Klf9/13* KD relative to Scramble Controls at P18-20 plotted against gene expression changes between P2 and P30 (as in **Figure 5B**). Red points are DEGs identified in *Klf9/13* KD samples. Green points are DEGs uniquely identified in *Klf6/7* KD samples at P10. Lines are linear fit +/- 95% CI to Klf9/13 (red) or Klf6/7 (green) DEGs. **B)** As in **A**, but changes in *Klf6/7* KD relative to Scramble Controls at P10-12 against gene expression changes between P2 and P30.

**Figure S8.**
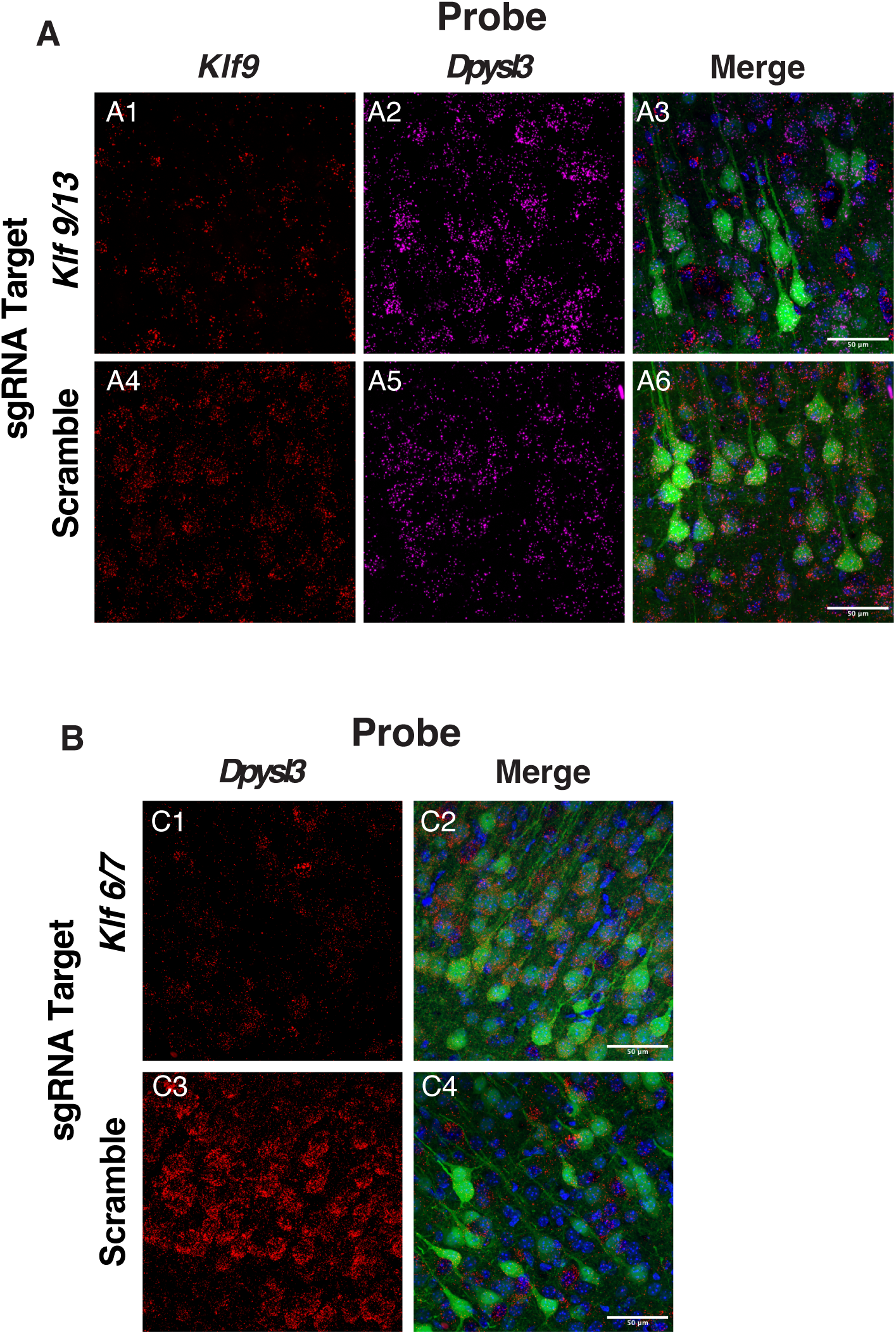
FISH validation of bidirectional regulation of *Tubb2b* and *Dpyls3* by KLF. **A)** Maximum intensity projections of slices taken from P20 mice receiving i.c.v. injections of Scramble sgRNA or *Klf9/13*-targeting sgRNA AAV9 at P0.5 probed for *Klf9* (**A1,A4**) and *Dpysl3* (**A2,A5**). Infected cells receiving sgRNA are EGFP-positive (**A3,A6**). Scale bars: 50µm. **B)** Maximum intensity projections of slices taken from P10 mice receiving i.c.v. injections of Scramble sgRNA or *Klf6/7*-targeting sgRNA AAV9 at P0 probed for *Dpysl3* (**B1,B3**). Infected cells receiving sgRNA are EGFP-positive (**B2,B4**). Scale bars: 50µm.

**Figure S9.**
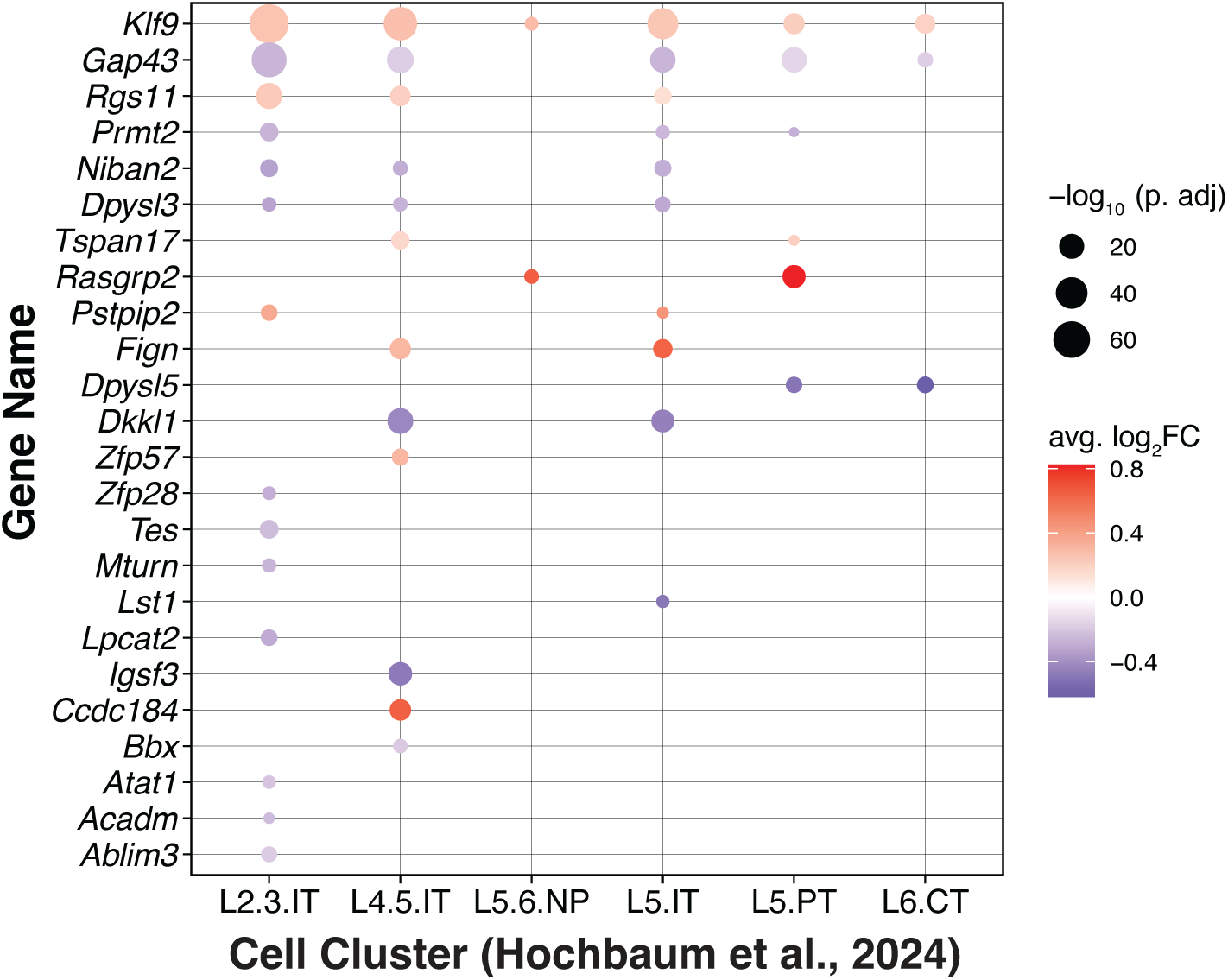
Thyroid Hormone (T3) treatment elevates *Klf9* and reduces the expression of KLF Targets in Cortical Neurons. Differential expression of KLF shared targets between T3- and vehicle-treated mice within excitatory cortical neuron cell type clusters annotated by Hochbaum et al., 2024. Differentially expressed KLF shared targets identified within a cluster are indicated by dots colored by fold-change and scaled by adjusted p value. (IT = intratelencephalic; NP = near-projecting; PT = pyramidal tract; CT = corticothalamic)

**Supplementary Table 1:**
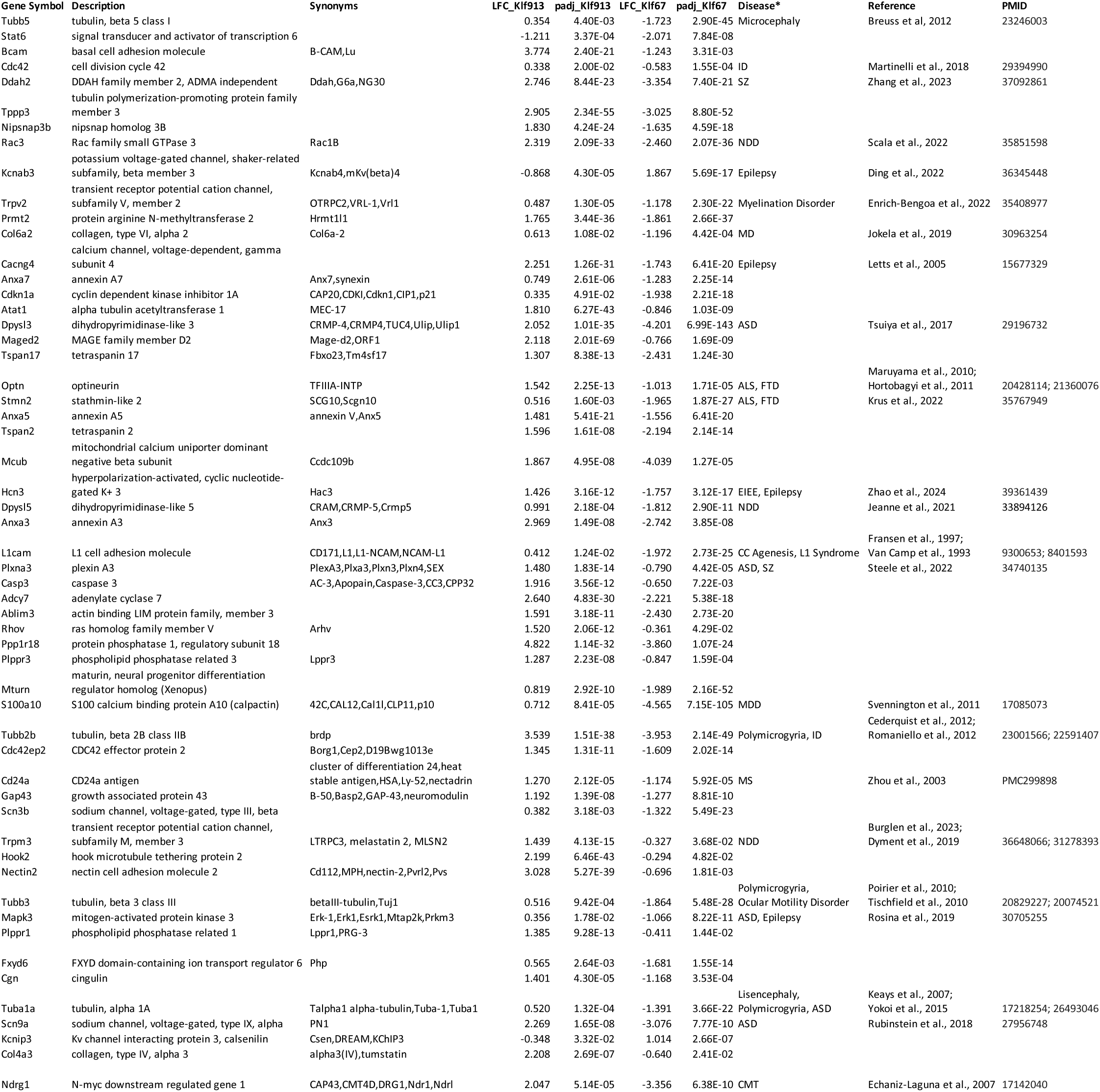
Disease-Associated targets of KLF Activators and Repressors A curated list of putative KLF targets with links to neuronal development and disease. Log2Fold-Change (LFC) and adjusted p value (padj) taken from DESeq2 comparisons to age-matched scrambled controls. \***Disease Categories:** ASD: Autism Specturm Disorder; CC: Corpus Callosum; CMT: Charot-Marie Tooth Syndrome; EIEE: Early Infintile Epilepsy; FTD: Frontotemporal Dementia; ID: Intellectual Disability; MDD: Major Depressive Disorder; MD: Muscular Dystrophy; NDD: Neurodevelopmental Disorder; SZ: Schizophrenia

